# A maximum entropy model for *m*^6^*A* prediction: from human transcripts to influenza A viral segments

**DOI:** 10.64898/2026.10.26.761909

**Authors:** Ivan Lecce, Pierre-Guillaume Brun, Simona Cocco

## Abstract

N6-methyladenosine (*m*^6^*A*) is the most frequent mRNA modification in eukaryotic cells and is involved in the regulation of gene expression. Since viruses can hijack the host machinery to perform this chemical editing, *m*^6^*A* is also involved in viral replication and innate immune system activation. Nowadays only few *m*^6^*A* viral maps are available, making complementary methods crucial to accelerate the investigation of these mechanisms. Here we propose MEMAR, a Maximum Entropy model, to predict the methylation frequency, depending on the eight-mers nucleotide motif surrounding the central adenosine. MEMAR is a fast, simple and modular model: it can be trained on human methylation maps and refined including the methylation level per transcript and the position of motifs relatively to the coding regions. This modularity allows strong adaptability, leading MEMAR to better perform over influenza A segments than the mammalian state-of-the-art DeepSRAMP method. MEMAR generalization potential is assessed on i) human transcripts from GLORI and miCLIP experiments ii) HEK293T and HeLa cell lines iii) positive and negative influenza A viral segments and iv) altered-methylation experiments. A two-state dynamical model between *m*^6^*A* writers and erasers explains MEMAR predictive power as a function of the global methylation percentage, and allows us to interpret anomalies in altered conditions through variations of methylation or de-methylation binding constants.

## Introduction

All cellular RNA transcripts are heavily modified in the cell, including base editing outside of the canonical A, C, G and U. In particular, messenger RNAs (mRNA) can display a large number of modified bases, such as pseudo-uridine [1], Adenosine-to-Inosine editing [2], N5-methylcytosine [3, 4], or the widespread N7-methylguanosine also known as mRNA cap [5]. These modifications are involved in mRNA splicing, maturation and localization [6], and have also been identified as interesting therapeutic targets to interfere with transcript translation and restore functional proteins [7].

Amongst all these modification, N6-Methyladenosine (*m*^6^*A*) constitutes the most frequent base editing found in mRNA [8]. The methylation of adenosines is carried out by the METLL3/METLL14 complex, mostly in the cellular nucleus after transcription [9, 10]. This complex, which is a methylation writer, has higher affinity to short motifs in transcripts known as DRACH (G/A/U - G/A - A - C - C/A/U) *via* a zinc-finger domain involved with RNA binding, carried by the METTL3 N-terminal segment [11]. The specificity to some RNA sequences is potentially further increased by Arginine-Glycine repeats on the C-term of METTL14 [12] and other regulatory proteins of the complex [9].

N6-methylation of adenosines has many roles in the cell, arising from a variety of proteins known as readers, including YTHDC1, YTHDF1/2/3, IGF2BP1/2/3 and eIF3 [13], that can recognize *m*^6^*A* to regulate transcripts nuclear export and localization, transcript splicing, stability, translation efficiency [13, 14], gene silencing (such as the X-silencing process [13, 15]) or to be recruited by active transporters to promote distant translation [14, 16]. The methylation can also be negatively regulated by eraser proteins, such as the ALKBH5 protein and the Fat mass and Obesity-associated protein FTO, which mutations are known to be implied in fat regulation [17]. Finally, disruption of the *m*^6^*A* homeostasis has implication both *in vivo* and *in vitro*, e.g. defects in cell fate such as haematopoiesis, tumour development and cortical lamination [17, 13, 18], cell signalling, e.g. formation of deleterious double-stranded RNA triggering the immune system response [19, 20], signalling of G protein-coupled receptors [21], or cell toxicity, disrupting glucose metabolism [22] or the mitochondrial respiratory chain [23].

Despite all of these phenotypic observations, much less is known about the molecular mechanisms bridging the regulation of N6-adenosine methylation and gene expression. A major effort has been put over the last decade to develop experimental techniques for *m*^6^*A* detection in the cell transcripts, with antibodies-based methods being a crucial turning point. Amongst these, MeRIP [24] and PA-*m*^6^*A*-seq [25] at low resolution (around 100-200 and 30-50 nucleotides respectively), or miCLIP [26, 27] (at single-nucleotide resolution) being the most popular. New techniques have also emerged recently at single-nucleotide resolution, either by direct third-generation sequencing [28], or by the enzymatical and chemical modification of bases (e.g. DART-seq [29], SELECT [30], *m*^6^*A*-SEAL [31]). The latest works have focused on the *in situ* observation and localization of *m*^6^*A via* the FISH techniques, e.g. DART-FISH [32].

Amongst all these experimental techniques, a recent method is GLORI [33, 34], which offers interesting properties and produces highly reliable *m*^6^*A* mapping. The protocol relies on the glyoxal- and nitrite-mediated deamination of adenosine in the total mRNA. Following reverse transcription and mapping on a reference transcriptome, *m*^6^*A* resist the deamination and are read as adenosine during sequencing while non-methylated adenosines are deaminated into inosines and are read as guanosines. The chemical treatment is able to detect much more methylated sites at single-nucleotide resolution, displaying high reproducibility between experimental replicates and allowing to estimate the methylation frequency of adenosines, namely the *stoichiometry*. GLORI achieves a chemical reaction efficiency of 98 to 99% A-to-G conversion. It is expected to be less prone to site bias than antibody and enzyme reliant techniques and more robust to sequencing errors than third-generation sequencing (despite noticeable recent progress on this issue).

These experimental developments are complemented by computational methods for predicting *m*^6^*A* in transcriptomes, developed in the last decade, starting with iRNA-Metyl [35]. Amongst these, the most popular predictors are WHISTLE [36], DeepPromise [37], MASS [38] or DeepSRAMP [39], which all made use of single nucleotide CLIP data, lacking methylation frequency signal, achieving promising performance at the cost of deep learning architectures. These predictors are assessed on datasets composed of 10% methylated DRACH sites and 90% non-methylated DRACH. For fair comparison, the imbalance is thus fixed independently of the experimental detection of *m*^6^*A* sites. Efficiency assessment on single mapped transcripts, keeping the unbiased number of methylations, remains although crucial to evaluate the real predicting capacity while generalizing to different datasets, and to distinguish computational limitations from potential biological effects, such as extrinsic features, leading to dynamical *m*^6^*A*s.

Recently, a great interest has also been devoted to the implication of *m*^6^*A* on viral replication, host’s machinery hijacking and triggering of the immune response (such as RNA viruses HIV-1 [40], Influenza A Virus [41], SARS-CoV-2 [42], Hepatitis C virus [43] and DNA viruses Hepatitis B [44, 45] or Herpesviruses [44]) holding a great potential for future cutting-edge therapeutic treatments. A very limited amount of single-nucleotide maps are nowadays available, and detection remains challenging due to several reasons, such as difficulties in distinguishing unique profiles at different infection stages [46]. This prompts the need for the development of complementary analysis to carry on the investigation through accurate and most importantly general and adaptable *m*^6^*A* prediction algorithm.

In this direction, Maximum Entropy Models come at hand as unbiased ways to describe data. They follow the Laplace principle of Insufficient Reason, where all events are equally probable in absence of relevant evidence [47, 48]. These energy-based machine learning models allow to match a set of measurable features by maximizing the entropy of the model. They are fast to infer, easy to train on a limited amount of data, and interpretable thanks to a low number of explicitly designed parameters. Their adaptable architecture has allowed a variety of applications, from the investigation of viral entropic forces under sequences’ constraints [49] to the inference of neural networks from recording data [50, 51] and of generative models from protein sequence data [52]. In this work, we introduce **M**aximum **E**ntropy **M**odel for *m*^6^**A R**ecognition (**MEMAR**), a predictor of methylation probability of 8-mer motifs surrounding a central adenosine, going beyond the DRACH-motif preference, built on GLORI data. To reflect the observed variability of methylation in the transcriptome and the higher density of *m*^6^*A* near the STOP codon and 3’UTR [24] we include in the model a flexible *prior* knowledge of the methylation level per transcript and along the transcripts.

At first MEMAR performance are assessed over high quality GLORI *m*^6^*A* data in human HEK293T cells, and results are generalized to human HeLa cell line and different experimental conditions impacting methylation levels to assess model generalization and the influence of different biological contexts on predictions. Subsequently comparison of performance are shown with respect to less extensive miCLIP data over HEK293T to further estimate the experimental technique impact. Our results, both for accuracy and runtime, are later compared to DeepSRAMP [39], which is a popular state-of-the-art predictor that achieves great performance on mammalian transcriptomes. MEMAR full adaptability capacity is then shown over a strain of Influenza A, a negative-sensed virus for which experimental methylation maps, produced in infected human cells, are available. Finally we present MEMAR robustness to out-of-sample predictions. We show how results numerically scale with *m*^6^*A* percentage, establishing its generality from a theoretical standpoint with a two-state dynamical model between *m*^6^*A* writers and erasers. Anomalies to this scaling, in altered experimental conditions, are further explained by non-uniform variations of methylation or de-methylation binding constants for particular motifs

## Material and Methods

### Data

#### Mapping *m*^6^*A* positions and filtering

Since experimental methods exploit polyadenylation selection (PolyA^+^ [53]) to retrieve cellular mRNAs, we mapped detected *m*^6^*A* sites onto the longest isoforms of *mature* RNAs. Genome reference GRCh38 was used for GLORI [33] [34] and GRCh37 for miCLIP data [26]. We assigned to all adenosines a methylation label *m* to distinguish between mapped methylated sites (*m* = 1) and non-methylated ones (*m* = 0). Genes containing any adenosine mismatch at the mapped *m*^6^*A* position were fully discarded.

Two replicate experiments, with 64.7% overlap [34], were used for training the models (and later for validation), over GLORI data (Tabs. 1, 3). All *m*^6^*A* sites identified in these experiments were pooled together. The stoichiometry of a site was computed as the ratio between the sum of methylated reads and the total sum of reads bearing that site in the two replicates.

**Table 1:** Training Dataset specifications.

| Training | Genes | $m^6A$ | A | Accession |
| --- | --- | --- | --- | --- |
| GLORI HEK293T | 6 796 | 52 381 | 2 000 000 | GSM7438377, GSM7438378 |
| CIMS miCLIP HEK293T | 1 170 | 1 881 | 1 102 536 | GSM1556679 |

#### Methylated and non-methylated 8-mer dataset

From experimental *m*^6^*A* sites, mapped over mature transcripts, we selected all shorts motifs of four nucleotides before and after methylated (*m*^6^*A*) and non-methylated adenosines (A). By defining the DNA alphabet, without any loss of generality, *A* = *{A, C, G, T }*, the motifs can be formalized as a vector of letters 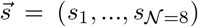, with *s*_*i*_ ∈ *A*. All 8-mers surrounding methylated *m*^6^*A* or non-methylated *A* sites were grouped respectively in the dataset *D*_*methylated*_ (of size *M*), or *D*_*non*-*methylated*_ (of size *N*). For GLORI maps, we additionally defined the vector 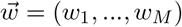 having as components the *stoichiometry* of each methylated site.

#### Transcript-dependent methylation rate

For a given experimental dataset, we define 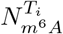 as the amount of methylated adenosines in transcript *T*_*i*_ and 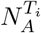 as the total number of adenosines. We then compute the methylation rate *p*_0_(*m* = 1) in the transcript *T*_*i*_ as:

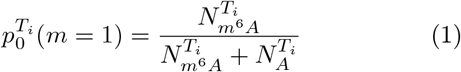

#### Overall methylation rate

We compute the overall methylation rate in a given dataset by averaging the transcript-dependent methylation rate (Eq. 1) amongst all *N*_*T*_ transcripts:

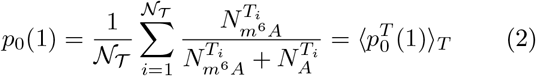

#### *m*^6^*A* position-dependence along the transcript

For all coding transcripts in a dataset, adenosines sites were projected onto a meta-transcript profile made by 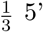-UTR, 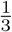 CDS and 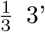-UTR. Denoting the mapped positions in this meta-transcript as *x*, after averaging over windows of 20 nucleotides, and setting a bin size 0.001, a density distribution for methylated adenosines *d*(*x* |*m* = 1) was computed. The same was done to compute a density distribution of non-methylated adenosines *d*(*x* |*m* = 0) (for resulting profiles over training sets see Suppl. Fig. S7).

#### Stoichiometry analysis on DRACH motifs

We consider a methylation GLORI dataset *D* and a DRACH motif 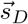. Defining 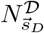 as the total number of times the methylated motif 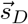 is present in the dataset, and 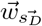 as the vector of associated experimental stoichiometry, we compute the average experimental stoichiometry as: 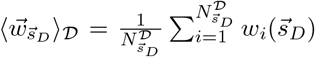. In order to retain statistically meaningful means, only methylated DRACH motifs present at least 50 times are considered and for those 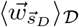 are compared against the mean predicted methylation probability 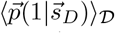. For a fair comparison, since the components of 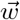 are always higher than 0.1 due to experimental filtering [34], we restrict to predicted probabilities with the same condition. To this end, we define 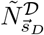 as the number of times the methylated motif 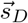 is present in the dataset having predicted methylation probability higher or equal than 0.1 and 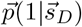 the vector of predictions associated for each considered motif. The mean predicted stoichiometry is therefore computed:

### MEMAR Model

Two maximum entropy models can be built to reproduce the statistics of the 8-mers surrounding a central adenosine from *D*_*methylated*_ and *D*_*non*-*methylated*_ datasets. Recalling the *methylated* label (*m* = 1) and the *non-methylated* one (*m* = 0), the model probability of an 8-mer in each dataset can be written as:

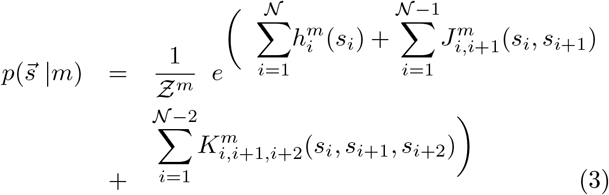

with 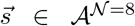and *Z*^m^ a normalization constant which can be found analytically (for full details see Appendices A1-A4). The parameters of the models 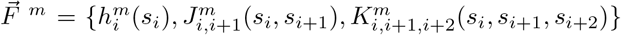 correspond to a set of site-dependent *selective forces* [49], which constrain, at each position the average frequency of, respectively, all mono-, di-, tri-nucleotides from the probability distribution to reflect the ones in *D*_*methylated*_ and *D*_*non*-*methylated*_ data respectively (Suppl. Eqs. S2-S4). This model is a site-dependent generalization of a previously developed Maximum Entropy model, deployed for viral-host recognition task [54]. Once the parameters are inferred from the data, the model can be used to predict the probability of an adenosine surrounded by a given 8-mer to be methylated or not. We have developed four different model classes for this prediction, depending on the additional *prior* knowledge of the tested sequence: MEMAR-C, MEMAR-CT, MEMAR-X and MEMAR-XT.

#### MEMAR-C

*The prediction requires the 8-mer motif and the prior knowledge of the overall methylation rate*. We write the probability of methylation (Appendix A5) as:

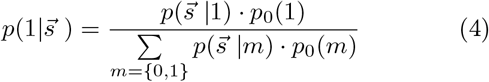

where the *prior* value *p*_0_(1) is the overall methylation imbalance (Eq. 2), quantified by the average level of methy-lated adenosines and *p*_0_(0) = 1 − *p*_0_(1) of non-methylated adenosines in a whole dataset. See Priors paragraph and Suppl. Tab. S1 for *p*_0_(1) values used.

#### MEMAR-XC

*The prediction requires the 8-mer motif, its relative position along the meta-transcript and the prior knowledge of the overall methylation-level*. If the position of the 8-mer in the mature coding transcript is known, the model methylation probability can be modulated by a probability density *d* depending on the position *x* of the motif 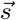along the meta-transcript either from *D*_*methylated*_ and *D*_*non*−*methylated*_ (see ***m***^**6**^***A*** position-dependence along the transcript paragraph):

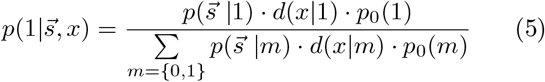

This position-dependent function *d*(*x m*) can be applied in testing transcripts only if the codon START and STOP positions are known.

#### MEMAR-T

*The prediction requires the 8-mer motif and the prior knowledge of the transcript-dependent methylation rate*. A third strategy applies when additional information is known regarding the transcript-dependent imbalance between *m*^6^*A* and unmodified adenosine. By defining 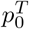 (1) (Eq. 9), which quantifies the methylation ratio for a given transcript, we write the probability to methylate as follows:

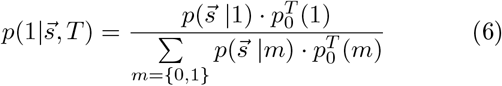

#### MEMAR-XT

*the prediction requires the 8-mer motif, its relative position along the meta-transcript and the prior knowledge of the transcript-dependent methylation rate*. At last, if both the knowledge of methylation imbalance and the relative position of the 8-mer in the transcripts are known, a complete probability distribution can be assigned following the expression below:

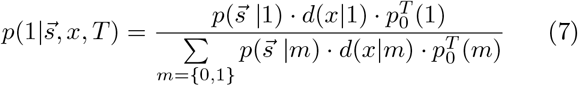

This final class combines all three information: sequence statistics, position and transcript-dependent imbalance.

#### Marginalization

To compute the methylation probability of motifs smaller than 8 nucleotides (such as the DRACH motifs), we marginalize the probability distribution learnt from data, 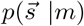, following the general expression:

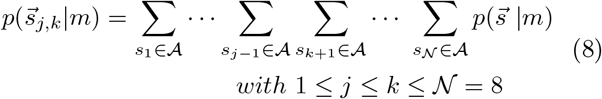

Where 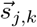 represents the sub-vector of 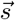 component *j* to *k*.

#### Performance indicators

We consider different ways to assess model performance. Choosing a threshold *h* (from 0 to 1), motifs are predicted to be methylated if their methylation probability *p* satisfies the following condition: *p* ≥*h*. Motifs are predicted as non-methylated otherwise. Moreover motifs are distinguished based on their experimental label, methylated (P) or non-methylated (N). Predictions are True (T) if they correspond to correct label, False (F) otherwise.

This procedure allows to plot two curves, the Receiver Operating Characteristic Curve (ROC) and the Precision Recall Curve (PRC) and to compute the Area Under the Receiver Operating Curve (AUROC) and the Area Under the Precision Recall Curve (AUPRC). Both quantities provide complementary measures of classification performance. The ROC curve is obtained by computing the True Positive Rate, 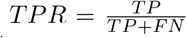, and False Positive Rate, 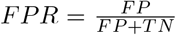, while the PRC is constructed by considering the *recall*, which is equivalent to the *T PR*, and the *precision* 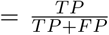. The ROC curve is insensitive to class imbalance between methylated and non-methylated sites because both *FPR* and *T PR* are normalized within each class by the true number of methy-lated (*T P* +*FN*) and non-methylated (*FP* +*T N*) sites. In contrast, the Precision–Recall curve is influenced by the class imbalance, making it more reliable, in this context, for performance evaluation [55]. In particular, precision is normalized by the number of sites predicted as positive by the model (*T P* + *FP*), which includes both correctly and incorrectly predicted methylated sites and mixes sites belonging to the two classes.

A third indicator is the F1 score: *F* 1 = 2·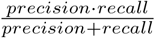, which corresponds to the harmonic mean between precision and recall. Maximizing F1 allows to set model best classification threshold in terms of accuracy and retrieval of true methylated sites.

#### Training

Training was performed over sets *T* ^*′*^ of genes, by selecting all shorts motifs as described in Methylated and non-methylated 8-mer datasets. We computed the frequency of all mono-, di-, tri-nucleotides (at each position of the 8-mer motifs) from *D*_*non*−*methylated*_ and *D*_*methylated*_. In the case of training from GLORI dataset, the frequency of sub-motifs in the methylated dataset is weighted by the stoichiometry 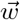 (Suppl. Eq. S3). In order to speed the model’s parameters inference, the numerical computation of the gradient (Appendix A2) was optimized performing automatic differentiation exploiting the ReverseDiff package and momentum algorithm [56] for the minimization step. Lastly, the number of non-methylated adenosine *N* was limited to two millions (randomly selected) in training dataset from GLORI data (Tab. 1).

#### Priors

The *prior* knowledge of the overall methylation rate was computed using Eq. 2 with training datasets from GLORI or miCLIP data (Tab. 1, Suppl. Tab. S1).

#### Transcript-dependent priors

To avoid bias, we compute the transcript-dependent *prior* using an independent dataset different from both the train and test one. Defining the gene in the test set as *T*_*i*_ and the same gene in the *prior* dataset as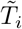, we generalize Eq. 1 with the following expression:

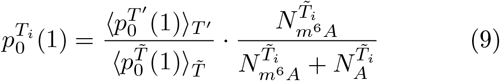

where averages correspond to the constant *priors* defined in Eq. 2 of the training set at the numerator, and of the independent *prior* set at denominator. This was chosen for GLORI experiments on HEK293T cell line from the deep sequencing data [33] while for miCLIP experiments over the same cell line using Cross-linking-Induced Truncations data (CITS) opposed to training over Cross-linking-Induced Mutation Site (CIMS) [26] (Tab. 2).

**Table 2:** Prior Dataset specifications.

| Transcript-dependent *Prior* | Genes | Accession |
| --- | --- | --- |
| GLORI HEK293T | 13 836 | GSM6432590 |
| CITS-miCLIP HEK293T | 1 798 | GSM1556678 |

#### Validation

An independent set (Tab. 3) was used to determine the models’ hyper parameters, which correspond to L2-regularization strength [52] and tolerance for gradient computation (Appendix A2). Both quantities allow the models to avoid over-fitting, and were chosen to maximize AUPRCs over all *m*^6^*A* motifs following Eq. 4 (Suppl. Fig. S1).

**Table 3:** Validation Dataset specifications.

| Validation | Genes | $m^6A$ | A | Accession |
| --- | --- | --- | --- | --- |
| GLORI HEK293T | 1 246 | 9 309 | 1 512 833 | GSM7438377, GSM7438378 |

#### Testing

*Test datasets were built exclusively from single experiments*, by taking a set of coding genes *T* different from the training ones, with at least one detected methylated site (Tab. 4). The model assigned a probability to 8-mers 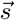surrounding an A to be methylated. All adenosines at site *i* near the transcript extremities, such as *i* ≤4 and *i* ≥*L*_*T*_ −3, where *L*_*T*_ is the tested gene length, were assigned a null score .

Test sets were chosen: i) keeping real data imbalance between all methylated and non-methylated motifs, ii) only considering DRACH motifs, iii) rebalancing datasets with 10% sites from *D*_*methylated*_ and 90% from *D*_*non*−*methylated*_ without any bias in position along the transcripts, iv) rebalancing datasets as (iii) but varying the amount of methylated motifs in the test set.

For the comparison with miCLIP [26], only a common set of genes between miCLIP and GLORI datasets [34] were retained for testing at first. This condition was relaxed for the comparison with DeepSRAMP. For performance analysis over different cell lines, we used GLORI data with higher sequencing depths [33]. All genes (different from the training ones) were kept as test ones from HEK293T in control conditions (corresponding to unaltered methylation levels). We made the same for HeLa in control conditions. For altered datasets (impacting the methylation levels), if a gene was not present in the experimental data while it was in the control condition of the corresponding cell line, all its 8-mer motifs were considered as non-methylated. Averages and standard deviation were computed by separating testing genes in three independent sets.

#### Performance evaluation against DeepSRAM

Two transcripts’ datasets containing 1259 and 301 genes were used to compare the performance of DeepSRAMP [39] and MEMAR from GLORI and miCLIP data respectively (Tab. 4). Genes that were part of the MEMAR training set were excluded to ensure a fair comparison.

DeepSRAMP was run using the “full_400_mature” model provided by authors at the [DeepSRAMP Web-server](https://www.cuilab.cn/deepsramp/). The training of this model was performed over mature transcripts incorporating both genomic and sequence features extracted from an 800-nucleotide window surrounding each methylated site. The human reference assembly GRCh38 and its associated annotation file were used to detect genomic features, following the authors’ recommendations.

To evaluate runtime performance, 200 genes were randomly selected from test dataset. Predictions were performed independently for each gene. Since DeepSRAMP includes an initial BLAST step against the human genome, whereas MEMAR directly uses a reference table with gene names, the time required for the BLAST and sequence alignment steps were excluded from DeepSRAMP’s total runtime. This ensures a fair comparison of models testing-time.

For the comparison with IAV segments, the [Deep-SRAMP Webserver](https://www.cuilab.cn/deepsramp/) was directly used to produce predictions, selecting as organism “Other Species” and as mode: “Mature mRNA mode”.

#### Baseline DRACH model

In order to compare MEMAR predictions to a background model, we considered a simple baseline DRACH binary classifier, which divides motifs into methylated and non-methylated classes.

This classifier assigns probability 1 to all canonical DRACH motifs, and 0 to all other motifs containing a central adenosine. As a consequence, PRCs and ROCs collapse into single points, as varying the probability threshold does not alter the classification. The F1 score is therefore the only relevant performance indicator for this model.

### Two-state dynamical model

We consider a simple dynamical process (Appendix A6) based only on *m*^6^*A* writers and erasers leading to methy-lated *m* = 1 or non-methylated *m* = 0 states. Within this model the probability for a given motif 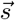 to be methy-lated, resulting from a two-state Markov model continuous in time [57], reads:

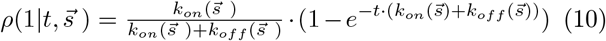

where *k*_*on*_ corresponds to the the reaction rate of the writer complex to perform the methylation and *k*_*off*_ corresponds to the reaction rate of an eraser to de-methylate the A. These are assumed to depend only on the motif 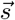 surrounding the A.

By assuming that the detected *m*^6^*A* sites in a given experiemental dataset are obtained after the equilibrium condition is reached:

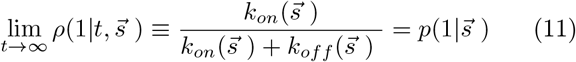

leading to the equilibrium relationship:

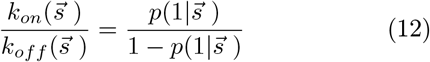

which links the kinetic parameters of the two-state model and MEMAR-C methylation probability. Notice that Eq. 12 only determines the ratio between 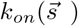 and 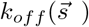; equivalent solutions are obtained multiplying 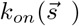 and 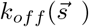 by the same factor. We choose 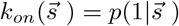and 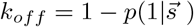.

#### AUPRC from the two-state model

We consider a testing dataset with an uniform distribution of 8-mer motifs (see Appendix A6 for generalization to the non-uniform case) and rank the possible *N*_*m*_ = 4^8^ 8-mers according to the MEMAR-C probability 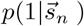, from 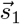 to 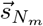. We assume as ground truth the methylation probability in the two-state model (Eq. 10). When ranking all the possible 8-mers according to MEMAR-C predictions we obtain at rank *n* the recall and precision defined by:

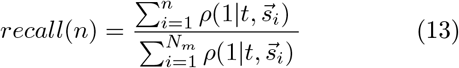

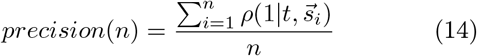

Moreover, the ratio of methylated adenosines, following the two-state model as ground truth is:

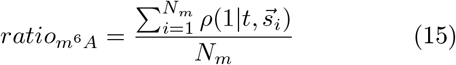

The precision-recall curve and its area, the AUPRC, as well as the methylation percentage can then be computed for each value of the time *t*.

#### Kinetic parameter variations in altered-methylation experiments

From the two-state dynamical model defined in Eq. 10 and its equilibrium solution (Eq. 11), we study how altering the methylation process affects *m*^6^*A* profiles. In our theoretical framework, under the assumptions that altered conditions change only the concentration of writers or erasers, we can model these alteration varying the writers/erasers binding constants by a multiplicative factor *α*_*on*_/*α*_*off*_ independently of the motif. We call this procedure an uniform rescaling of parameters. In particular in knock-down of methylation writer complex experiments, we consider 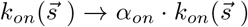with *α*_*on*_ *<* 1 leaving unchanged 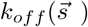, while for over-expression of eraser proteins 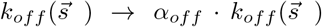with *α*_*off*_ *>* 1 and 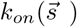 unchanged. Variations that do not target *directly m*^6^*A* writer or eraser concentrations, such as hypoxia or silencing of the reader YTHDF2, will be equivalently analysed as if modifications were solely impacting the methylation binding constants 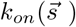. The multiplicative factors *α*_*on*_*/α*_*off*_ which best fit the data can be obtained by minimization of the L1-error given the probability 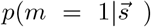 obtained by MEMAR-C re-inferred in each altered dataset (see Appendix A6 for details).

This uniform rescaling of kinetic constants is performed from MEMAR-C inferred on GLORI HEK293T training set (Eq. 12). After the rescaling, the equilibrium methylation probabilities of each motif are then compared to MEMAR-C probabilities re-inferred in each altered dataset. The validity of this uniform rescaling hypothesis, can be assessed comparing the methylation probabilities for each DRACH motif (see paragraph Marginalization), looking at the compatibility across three-independent datasets to compute the standard deviations. In particular L1-errors smaller than 0.005, 0.015 and 0.03, will be considered as giving respectively, strongly compatible, compatible or incompatible results. Any larger deviation is considered as strongly incompatible. We extend and validate the above comparison by computing the uniform rescaling starting from the kinetic constants of the two-state model obtained from MEMAR-C re-inferred on the control conditions (HEK293T or HeLa).

### Influenza A viral sequences

#### IAV Mapping and filtering

To generalize MEMAR performance over non-human sequences, a specific dataset was used for Influenza A Virus [58]. The methylations were identified combining meRIP-seq data and bioinformatic analysis after infection over A549 human cell line. Despite the access name (A/Goose/Sichuan/SC15/2015(H5N6)), the original viral sequence could not be found in genome databases. Since small sub-sequences of 13-mers centred around recognized *m*^6^*A* were provided by authors, these were put together for segment HA, in order to reconstruct as much as possible the full segment. BLAST [59] was run to find the sequences with the highest correspondence, and a closely-related strain with 95.83% identity was retained: A/Duck/Sichuan/NCXJ16/2014(H5N6). All segments of this strain were downloaded from NCBI and methylation were mapped accepting at most 1 nucleotide mutations around methylated adenosines (see for more details Tab. 5, Suppl. Tables S4-S7, and Suppl. Fig. S10).

#### IAV Prior and testing

Due to limited *m*^6^*A* experimental maps of Influenza A, we considered an independent dataset from the test set [58] for the *prior*. We grouped IAV segments into three classes, behind the hypothesis that they share similar methylation percentage: three polymerase segments (PB2, PB1, PA), two subtype ones (HA, NA) and the remaining (NP, M, NS). One positive strand segment for each class was chosen to form the *prior* determining set *I* = *{PA, NA, M}* weighted by the number of segments in the same class: 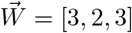. The constant IAV methylation *prior* was computed from the experimental imbalance of the chosen segments as follows:

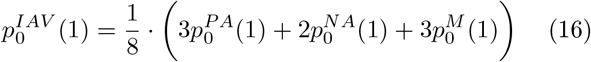

All positive and negative strand segments not belonging to the set *I* were taken as test set.

For the transcript-dependent *prior*, a different experimental setting was considered: PA-m6A-seq in HEK293T human cells infected with the viral sequence A/Puerto Rico/8/34/Mount Sinai (H1N1) [41]. As single-nucleotide cannot be resolved with this technique, the computation in Eq. 9 is below generalized. We considered the average methylation signal of segment *i* as the normalized area under the peak curve: 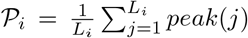 and the total average methylation signal over all segments 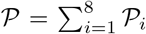. For each testing sequence, independently of the strand, the transcript-dependent *prior* was chosen as:

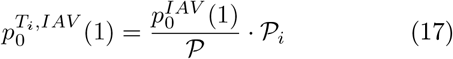

#### Threshold

In order to assign the methylated label (*m* = 1) and non-methylated label (*m* = 0) from MEMAR probability *p* in IAV, the threshold was chosen in the most general way to maximize the F1 score from MEMAR-XT in GLORI validation set in human HEK293T cells (Tab. 3). The corresponding value, 0.225, was used for both positive and negative strand predictions.

## Results

### MEMAR performance in GLORI data

We assess MEMAR performance comparing the four different classes (Eq. 4, 5, 6, and 7), progressively enriching the model with the *prior* knowledge of the distribution of methylation sites along the meta-transcript (see ***m***^**6**^***A*** position-dependence along the transcript) and the transcripts’ expected methylation level (see Transcript-dependent priors). We also compare MEMAR with a model that assigns probability 1 to all DRACH motifs, and 0 to the remaining 4-mers surrounding *A* (see Baseline DRACH model). We evaluate the performance on a test set made of GLORI HEK293T data [34], having an imbalance between the methylated and the non-methylated class of about 0.6% and 99.4% respectively, see Tab. 4. As shown in Fig. 1**A**, ROCs display similar results for all strategies with AUROCs above 0.96 values, exhibiting classification performance of high quality due to the insensitivity of ROCs to the imbalance in the test set, as discussed in the Performance indicators paragraph. Moreover, the baseline DRACH model, is already able to have a significant *T PR* at a very low *FPR*. The base-line DRACH model demonstrates a sufficient capacity to discriminate methylated from non-methylated motifs, even though it shows worse performance than MEMAR. MEMAR-C (Eq. 4) performs almost equivalently to the other model classes (Eq. 5, 6 and 7), indicating that the additional information provided (such as the *m*^6^*A* position or transcript-dependent imbalance) do not impact results in this setting and testing measure.

**Figure 1:**
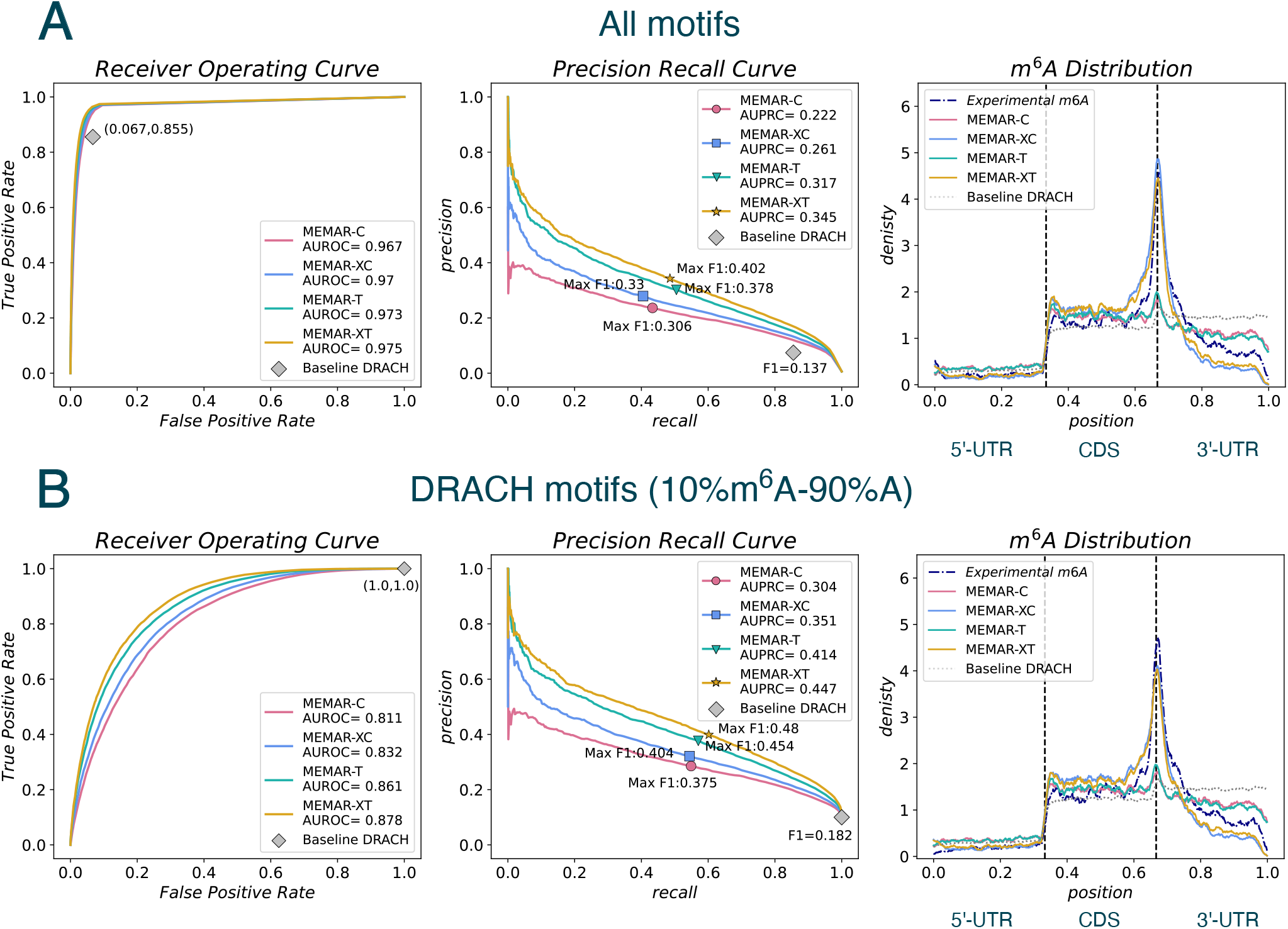
MEMAR tested on GLORI data of human transcripts from HEK293T cell line. The performance, on an in-sample test dataset (from [34]) of the four classes of MEMAR, are compared to the baseline DRACH model by plotting Receiver Operating and Precision Recall Curves, and reporting AUROC and AUPRC values along with the maximum F1 scores. The distribution of predicted *m*^6^*A* sites in transcripts at the threshold of the maximum F1 score is plotted to compare with the experimental one. **A)** Test dataset obtained without altering class imbalance between methylated and non-methylated adenosines. Due to the class imbalance, AUROCs are higher than AUPRCs. MEMARs always perform better than the baseline DRACH model. Including the *prior* and the transcript-dependent position distribution significantly increases the model score in AUPRCs compared to MEMAR-C. Only models MEMAR-XC and MEMAR-XT are able to reproduce the experimental localization of methylated sites with the enrichment near the codon STOP. **B)** Test dataset composed only of DRACH motifs, with a fixed imbalance of 10% *m*^6^*A* sites. AUROCs are lower compared to **A**, whilst richer MEMAR classes already impact classification quality in this testing metric. AUPRCs show a similar performance pattern to **A**. As in **A**, only MEMAR-XC and MEMAR-XT are able to reproduce the experimental localization of methylated sites, despite having slightly worse accuracy in the peak near codon STOP region.

The PRCs provide a complementary metric to asses classification quality compared to AUROCs in highly unbalanced data. In fact, *precision* depends on the amount of *m*^6^*A* and *A* sites, as it is defined by the ratio between number of true positives (*T P*) in the positive predictions. The latter changes with the value of the threshold separating the predicted *methylated* and *non-methylated* labels in the test set, ranked according to the methylation probabilities. The lowest *precision* is reached when all sites are predicted as positive, and corresponds to 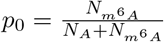. The higher the imbalance, the lower *p*_0_, and the lower bound on the *precision*. AUPRC values, being strongly affected by class imbalance, are therefore lower than AU-ROCs. Fig. 1**A** (central panel) shows that within the PRC metric, MEMAR performance increases with the complexity of the model class and the baseline DRACH model is underperforming with respect to MEMARs. The simplest model class MEMAR-C (Eq. 4) distinguishes motifs only based on their 8-mers statistics surrounding the central *A*. It therefore represents a generalization of the baseline model. At the maximum F1 threshold value, MEMAR-C predicts only 746 motifs to be methylated, which amounts of 1.1% of the possible 65536 8-mer motifs. This subset of 746 motifs encompasses 8 DRACH motifs and only one non-DRACH (Suppl. Fig. S3). The 10 remaining DRACH motifs are never predicted as potential methylation motifs by MEMAR-C, at the maximum F1-score threshold value. The narrower classification amongst the subset of the 8 DRACH motifs depending on the 8-mers surrounding the *A* increases the maximum F1 score from 0.18 in the DRACH baseline model to 0.38, demonstrating the importance of considering 8-mer motifs in modelling *m*^6^*A*. Richer model-classes, MEMAR-XC and MEMAR-T, taking as input the 8-mers motif and either the relative position of the motif in the transcripts or the transcript-dependent methylation rate (Eqs. 5, 6), reach better performance in the classification, with MEMAR-T being the best. Lastly the model-class MEMAR-XT, combining the position and the transcript-dependent methylation rate (Eq. 7), contributes to the highest score. In the richer model classes, motifs with the same statistics can obtain different methylation probability depending of either the position, the transcript, or both quantities; moreover, modelling longer motifs does not improve performance (Suppl. Fig. S2).

In Fig. 1**A** (right panel), the experimental position-dependent methylation probability density is compared to the predicted mean density on the *D*_*methylated*_ test set. Only MEMAR model-classes with the position-dependent *prior* (MEMAR-XC, MEMAR-XT) are able to replicate the known enrichment of *m*^6^*A* sites near the codon STOP [24], with only small discrepancy in CDS and 3’-UTR regions (respectively being slightly higher or lower than the experimental density). On the contrary, the difference of methylation levels between the 5’-UTR part and the rest of the transcript is already well captured by the simplest model, MEMAR-C, showing that this feature is well accounted by the sole sequence dependent enrichment/depletion of methylation motifs along the transcript.

In Fig. 1**B**, MEMAR performance is evaluated considering a test set composed of DRACH motifs only, as commonly done in the literature, and fixing the imbalance to 10% *m*^6^*A* and 90% *A* (see Testing paragraph for details). AUROC values show a substantial drop compared to the previous case. This can be explained by the harder classification task, which has shifted from classifying methylated *versus* non-methylated motifs in all possible 8-mers, to predicting methylations on the subset of 8-mers containing DRACH motifs, which are the preferred *m*^6^*A* sites. Differently from Fig. 1**A**, small performance variations amongst model classes can be noticed in ROCs. Although, similarly to PRCs, performance increases with the added complexity of the *prior*. This finding confirms that the added information are effective and consistent even when focusing on a smaller subset of 8-mers. The baseline DRACH model predicts all motifs as being methylated in this setting (since they are all DRACH) and therefore its performance corresponds to the point (1, 1). For the precision-recall, the baseline DRACH model is at the end point of PRCs, as all *m*^6^*A* motifs are retained (*recall* = 1) while making a proportion of errors corresponding to the imbalance in the data (*precision* = 0.1). For all MEMAR classes, AUPRC and maximum F1 scores are higher compared to the ones in Fig. 1**A**. Once again, only MEMAR-XC/XT model-classes, taking as input the distance *m*^6^*A* distribution, are able to replicate the observed higher density of methy-lated sites around the STOP codon.

### Performance comparison over human cell lines and altered methylation conditions

In this section we investigate the generalization of MEMAR-XT performance (Eq. 7) and to what extent biological features impact predictions, comparing nine GLORI experiments (obtained with higher sequencing depths), in HEK293T and HeLa cells [33]. These experiments include control conditions (with unaltered methylation level), and altered conditions with either regulation of the expression of specific genes (increased expression of the methylation eraser FTO, or reduced activity with knock-down of the writers METTL3, METTL14 and the reader YTHDF2, through silencing RNA [60]), and one hypoxic condition (induced stress from oxygen deprivation). The transcript-dependent *prior* (Eq. 9) is kept constant over cell lines and experimental conditions and set to the one previously used (see Transcript-dependent priors paragraph). As shown in Fig. 2**A**, AUPRCs generalize well from HEK293T to HeLa cell line in control conditions, (0.40 against 0.37). We distinguish the remaining predictions in two groups: one unifying all silencing RNA (siMETTL3, siMETTL14 and siYTHDF2) and hypoxia condition in HeLa cell line (with AUPRC values around 0.3) and another group with lower AUPRCs (around 0.2) including the inhibition of METTL3 activity by STM2457 in both cell lines, and the overexpression of FTO in HEK293T. The observed model deterioration is expected, as altered methylation datasets have lower *m*^6^*A* abundance (see Tab. 4). In fact, the AUPRC is sensitive to the imbalance between the methylated and non-methylated classes, leading, as shown in Fig. 2**B**, to a global decrease with the overall measured percentage of *m*^6^*A* (with a Pearson correlation coefficient of 0.93). The scaling with respect to the methylation percentage is covered in more detail in section A two-state dynamical model provides interpretation of performance scaling (Suppl. Fig. S11).

**Figure 2:**
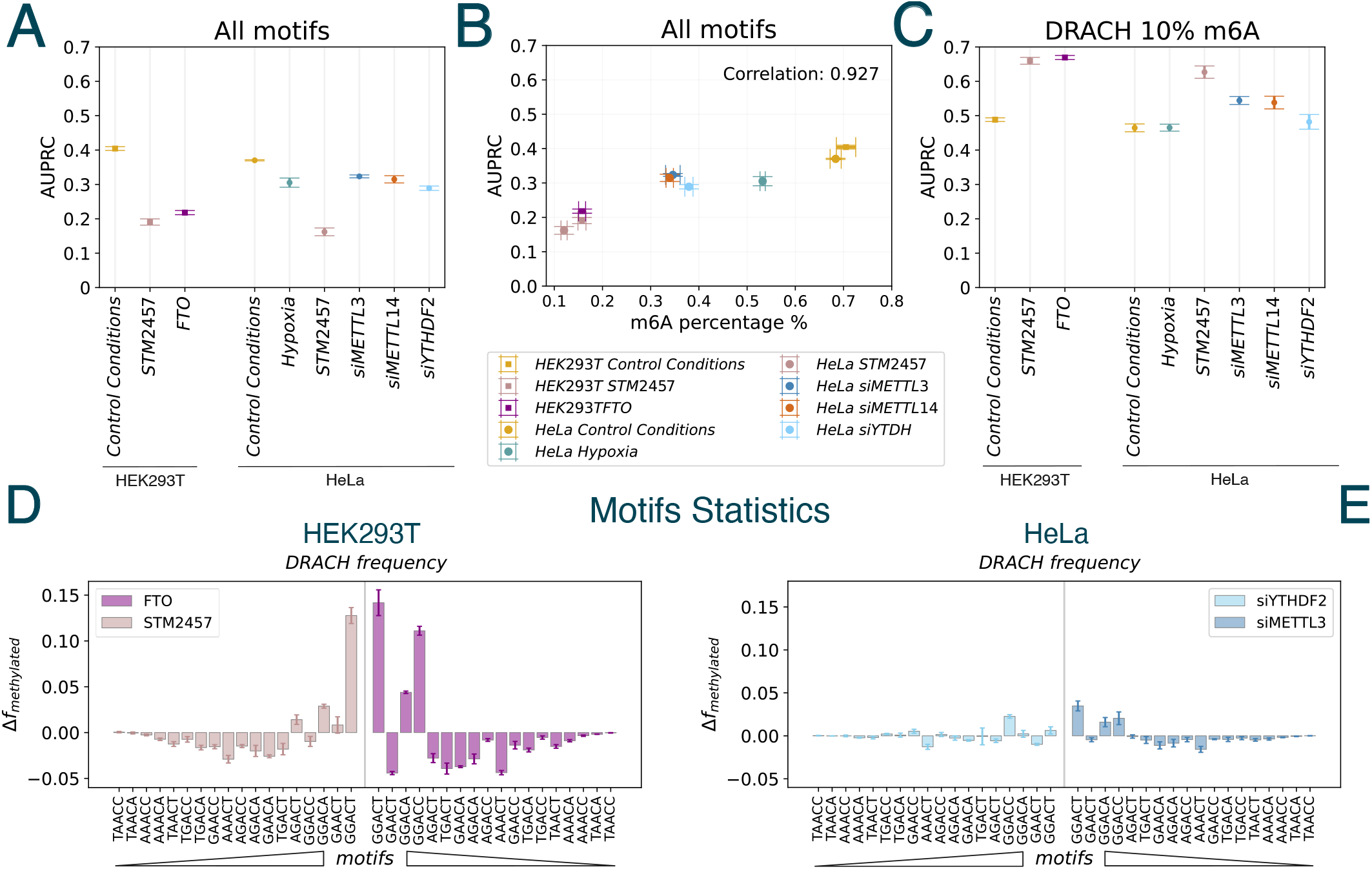
MEMAR performance over GLORI data in out-of-sample test sets obtained in HEK293T and HeLa cell lines in different experimental conditions. **A**) AUPRCs show three groups of performance for methylation predictions for all 8-mers surrounding an *A*, obtained by GLORI [33], in different conditions: control conditions for both HeLa and HEK293T (giving the best MEMAR performance), knock-down with silencing RNA methods and hypoxia in HeLa (with intermediate MEMAR performance), and lastly with FTO overexpression and METTL3 inhibition in both cell lines (with lowest MEMAR performance). **B**) Scatter plot of AUPRC scores versus *m*^6^*A* percentage imbalance, for all test datasets, showing how MEMAR performance increases with the percentage of detected *m*^6^*A* sites (0.927 Pearson correlation coefficient). **C**) AUPRCs on test sets of DRACH motifs at fixed 10% *m*^6^*A* imbalance. Performance pattern is inverted compared to panel **A.** Hypoxia, silencing of YTHDF2 and control conditions have the lowest performance, followed by silencing of METTL3 and METTL14, and lastly best performance are reached by FTO and inhibitor of METTL3 in both cell lines. **D**) Difference in the experimental methylation frequencies of the DRACH motifs, between FTO/STM2457 and control conditions, ranked by MEMAR predicted probabilities (from lowest to highest and vice-versa) showing higher proportion of top-scored motif. **E**) Same as **D** for siYTHDF2/siMETTL3 showing a much lower motif-dependent methylation difference compared to HEK293T case.

We further evaluate model generalization by controlling the methylation imbalance on the test dataset. We therefore filtered, as done in Fig. 1**B**, the experimental data over 8-mers containing DRACH motifs and fixed the class imbalance at 10% *m*^6^*A* and 90% *A* sites. A strong change of performance is observed in Fig. 2**C**, with STM2457-mediated inhibition of METTL3 in both cell lines and FTO over-expression reaching the highest AUPRC (around 0.65), followed by the knock-down conditions of the writer complex METTL3 and METTL14 in HeLa cells with siRNA (around 0.55). Lastly the control conditions, along with the hypoxia and the knockdown of the reader YTHDF2, have the lowest performance (around 0.48). Surprisingly, in this setting, the model performance in altered contexts are as good or better than HEK293T control condition. This unexpected behaviour could be linked to variations of the *m*^6^*A* deposition. In the following, we verify this hypothesis considering MEMAR module 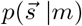.

We focus on four altered conditions, corresponding to FTO and STM2457 for HEK293T and silencing of METTL3 and YTHDF2 in HeLa. The remaining conditions are shown in Suppl. Fig. S4. In Fig. 2**D**, DRACH motifs were ordered according to the score given by 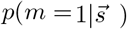 (see Marginalization paragraph and Suppl. Fig. S4). For each DRACH ranked motif 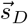, the methylation distribution *f*_*m*_ is computed as: 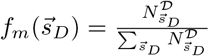 where 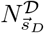 represents the total number of time the motif is present in a given methylated dataset *D*. The difference in the experimental methylation distribution Δ*f*_*m*_ between altered and control conditions is then reported. A strong variation is observed, in particular, at the predicted highest-ranked methylation motif *GGACT*, which is overrepresented in both FTO and STM2457 cases in HEK293T. Additional positive contributions for third and fourth-scored motif (*GGACA* and *GGACC*) are seen in FTO, with only much smaller contribution for STM2457. Since FTO and STM2457 altered conditions tend to restrict methylation to the highest scored motifs, MEMAR-XT methylated predictions on higher-scored sites have higher success rate when fixing the 10% *m*^6^*A* imbalance leading to higher AUPRCs. This effect is further explained in section A two-state dynamical model provides interpretation of performance scaling and predicted (Suppl. Fig. S15).

In HeLa cell line, the same analysis is presented for siYTHDF2 and siMETTL3 in 2**E**. The distribution difference of methylated motifs is much smaller compared to the results of panel **D**, with only first, third and fourth motifs showing a significant positive variation in siMETTL3 with an overall null-effect for YTHDF2 silencing. The equivalent accuracy of the model, at fixed 10% imbalance, between siYTHDF2 and control conditions can substantially be explained by negligible impact on motifs statistics, while the small gain in siMETTL3 can be due to slight variations on first motifs. Similar arguments explain model predictions, in the remaining conditions (Suppl. Fig. S4). To investigate the accuracy of the full MEMAR model, at fixed imbalance, the position and transcript-dependent modules are evaluated in Suppl. Figs. S5, S6, for altered conditions. No relevant variations in the *m*^6^*A* distribution throughout experimental conditions can be observed, consequently not impacting predictions.

### Performance comparison over GLORI and miCLIP data

In this section, we explore the performance dependency of MEMAR over data quality, training choice and experimental techniques in the HEK293T cell line. To this end, we trained models on GLORI [34] and miCLIP [26] data (for more details, see Training). We first focus on AUPRCs over all DRACH motifs with MEMAR-XT (Eq. 7). As shown in Fig. 3**A**, lower performance are retrieved over miCLIP tests independently of the data used to train the model. In particular, performance drop from GLORI to miCLIP test sets with GLORI train (from 0.507 to 0.079), and with miCLIP train (from 0.43 to 0.12). This is expected due to a much higher imbalance in miCLIP data due to lower detection power (see Tab. 4). Across rows, varying training models from GLORI to miCLIP, at fixed GLORI test, results in a small loss of −17% indicating that MEMAR retains good predictive performance even when trained on lower-quality experimental data. At fixed miCLIP test there is a variation of −34% from mi-CLIP train to GLORI. These results also confirm that in-sample performance are better than out-of sample ones.

**Figure 3:**
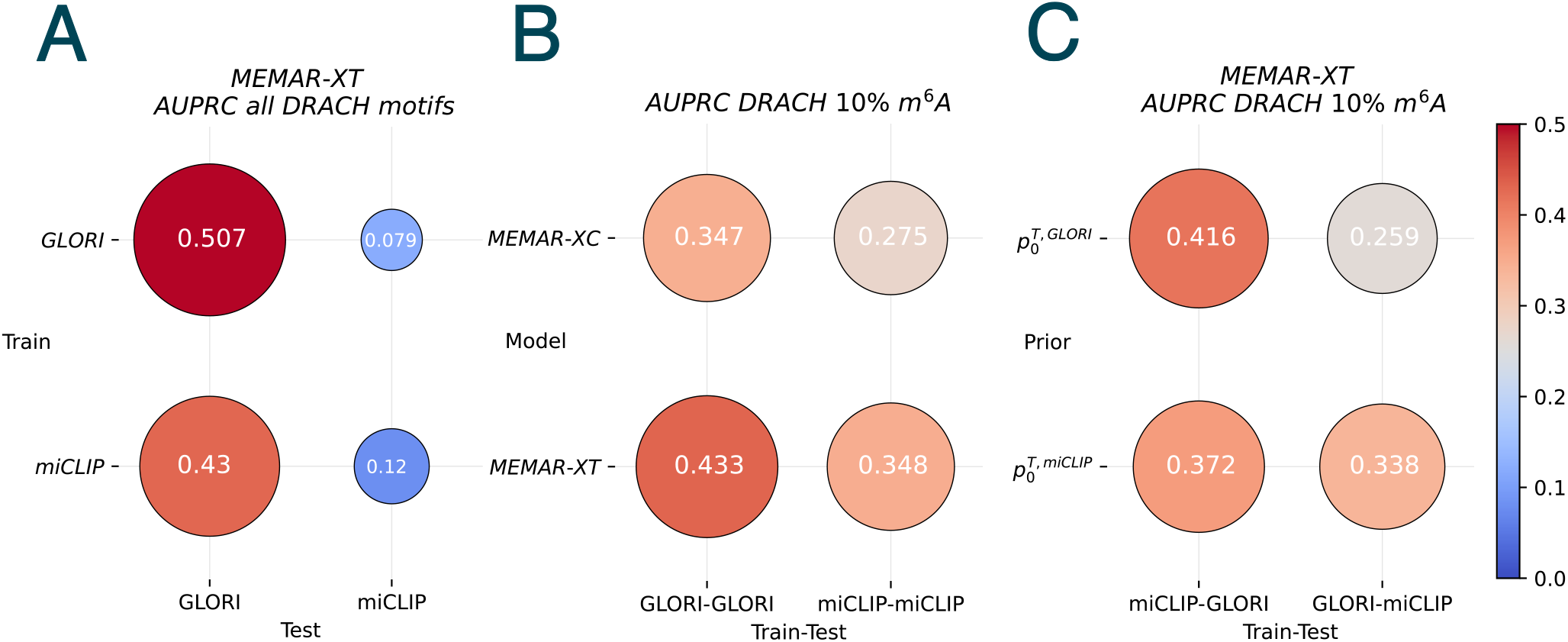
MEMAR performance comparing GLORI [34] and miCLIP [26] in HEK293T cell lines, as train or test datasets for *m*^6^*A* predictions. AUPRC scores on DRACH motifs are shown inside circles filled with the colour and the size reflecting model performance. **A**) MEMAR-XT performance as a function of the type (GLORI or miCLIP) of train/test datasets. A significant difference in performance arises only from the test set, with lower performance on miCLIP data. In-sample performance are higher than out-of-sample ones. **B**) Performance using either MEMAR-XC or MEMAR-XT in-sample test set, at fixed percentage of 10% *m*^6^*A* sites in test dataset. MEMAR-XC always underperforms compared to MEMAR-XT with lowest performance with train and test from miCLIP data. **C**) Performance for MEMAR-XT for out-of-class test sets, fixing imbalance to 10% *m*^6^*A*, as a function of the transcript-depend *prior*, either from GLORI or miCLIP data

Secondly, in Fig. 3**B**, we compared the performance between MEMAR-XC (Eq. 5) and MEMAR-XT (Eq. 7) training and testing on GLORI or miCLIP, with a *prior* following the given experimental technique. Evaluating the models using a fixed proportion of 10% DRACH *m*^6^*A* reduces as expected the differences in performance. In both cases, the transcript-dependent *m*^6^*A* rate improves the classification task, with increase of +24.8% for GLORI and +26.5% for miCLIP. These similar relative variations confirm, now in different techniques, the performance pattern already seen amongst MEMAR classes in section MEMAR performance in GLORI data. Smaller AUPRC values are retrieved over miCLIP tests for both model classes.

We lastly evaluated generalization performance of MEMAR-XT, training over miCLIP and testing on GLORI or *vice-versa* as a function of the *prior*. This is done at fixed methylation imbalance of 10% filtering over DRACH motifs. We first compare the performance at same train and test set from Fig. 3**B** (lower row), to the generalization of performance at fixed test *prior* in Fig. 3**C** (diagonal elements). Very small changes in performance are seen changing training data (from miCLIP to GLORI) and testing on miCLIP data when fixing the miCLIP *prior* (0.348 against the 0.338, accounting only of a relative variation of −2.89%). This shows that robust prediction can be achieved on lower quality data by simply adjusting the transcript-dependent *m*^6^*A* level to the test data. A similar variation is seen when changing the training data from GLORI to miCLIP, over GLORI test data at fixed GLORI *prior* (0.433 against 0.416, which is equivalent to −3.7% relative change), confirming how MEMAR is robust towards training set variations with good *prior* knowledge. We look at the most out-of-class generalization, with both complementary test data and *prior* (anti-diagonal elements). A marked difference between results can now be seen across rows of Fig. 3**C** when fixing the *prior* from the different experimental test set. A larger loss is seen for miCLIP test data (from 0.338 to 0.259), compared to GLORI test (from 0.416 to 0.372).

Our results confirm the importance of the knowledge of the transcript-dependent methylation level on the test set to achieve good predictions on data obtained with different techniques.

### Benchmark against a state-of-the-art predictor

In order to better understand how MEMAR results compare to state-of-the-art models, an analysis with DeepSRAMP [39], which is a popular predictor of mammalian *m*^6^*A* sites, is detailed in this section. Predictions are benchmarked against both GLORI and miCLIP experimental data in HEK293T cell line. As shown in Fig 4**A**, DeepSRAMP tested on miCLIP reached a AUPRC around 0.4. By comparison, MEMAR-XC trained on GLORI, reached 0.28 value while MEMAR-XT trained on miCLIP data with a transcript-dependent *prior* (Eq. 7), is slightly underperforming (AUPRC of 0.35) compared to DeepSRAMP. It is expected that a simple model such as MEMAR-XT, which considers only a position-distribution of methylation over the meta-transcript and the statistics of motifs, is not able to capture sophisticated features which are encoded in complex deep-learning architectures such as the one used by DeepSRAMP. If performance are tested on GLORI instead, DeepSRAMP has a little drop in AUPRC of approximately −17%, whilst MEMAR-XC and MEMAR-XT using respectively miCLIP training data, or GLORI data, have an increase of performance of +16% and +25% compared to test over miCLIP. Testing with high quality GLORI *m*^6^*A* data is therefore beneficial for MEMAR-XC as opposed to DeepSRAMP, despite being both trained on lower quality CLIP maps.

**Figure 4:**
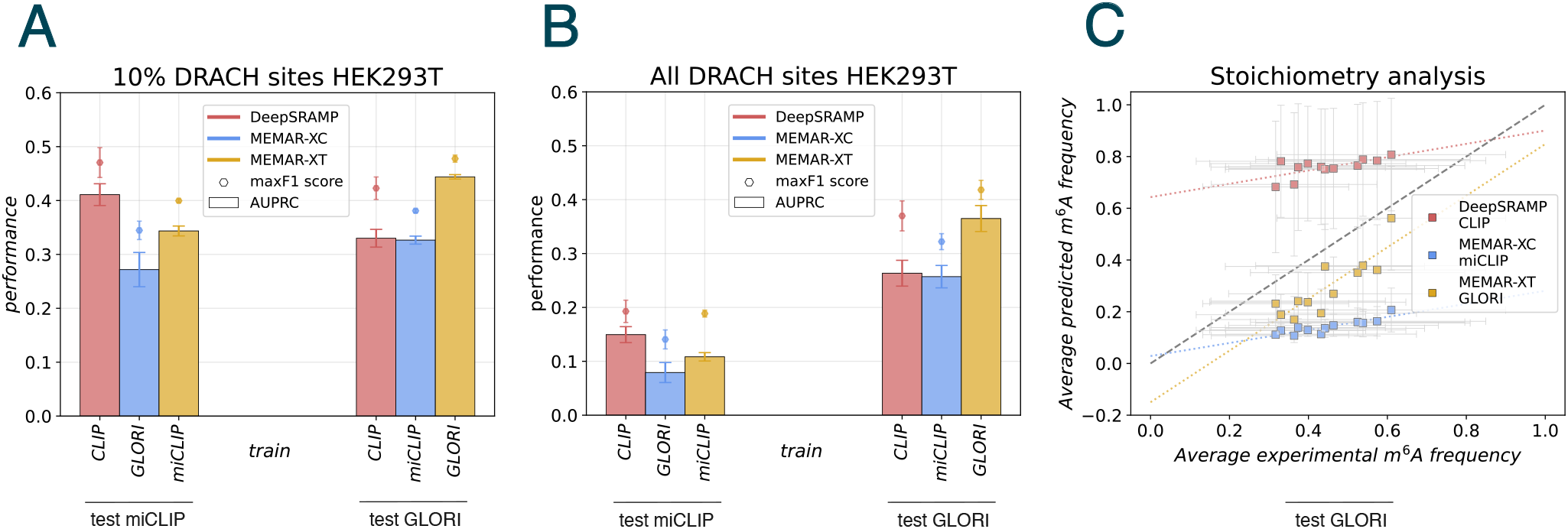
MEMAR performance compared to DeepSRAMP [39]. **A**) AUPRCs (bar-plots) and maximum F1 scores (dots) over miCLIP (from [26]) and GLORI (from [34]) test-sets in human HEK293T cell with fixed 10% methylation percentage. The performance of the mammalian *m*^6^*A* predictor DeepSRAMP are compared to MEMAR-XC and MEMAR-XT with constant or in-sample transcript-dependant *prior*. DeepSRAMP shows better performance compared to MEMAR-XC and MEMAR-XT when tested on miCLIP data. However, when tested on GLORI data, MEMAR-XC reaches equivalent performance to DeepSRAMP, and MEMAR-XT largely outperforms it. **B**) AUPRCs (bar-plots) and maximum F1 scores (dots) for DRACH motifs over miCLIP (from [26]) and GLORI (from [34]) test-sets in human HEK293T cell with true methylation imbalance. All models show a drop of performance, particularly noticeable on miCLIP test sets. **C**) Stoichiometry analysis of 12 DRACH motifs in the GLORI test set [34] from human HEK293T cell line. Average predicted *m*^6^*A* frequencies are plotted against average experimental stoichiometry. MEMAR-XT predictions are highly correlated to experimental data. MEMAR-XC trained on miCLIP data [26] and DeepSRAMP have comparable lower correlation to experimental stoichiometry with the highest and lowest offset respectively.

As shown in Fig 4**B**, when considering true data imbalance between methylated and non-methylated sites, all models tested on miCLIP experiments show a significant drop in performance due to the absolute low quantification of miCLIP data (see Table 4): −63% for DeepSRAMP, −72% for MEMAR-XC miCLIP and −69% for GLORI MEMAR-XT. The drop in performance is much less pronounced when the models are tested on GLORI data: −20% for DeepSRAMP, −21% for miCLIP

**Table 4:** Test Dataset specifications.

| Test | Genes | $m^6A$ | A | DR$m^6$ACH | DRACH | Accession |
| --- | --- | --- | --- | --- | --- | --- |
| GLORI HEK293T | 1 398 | 10 783 | 1 709 428 | 9 170 | 114 508 | GSM7438377 |
| GLORI HEK293T | 3 694 | 34 417 | 4 597 936 | 28 042 | 302 197 | GSM6432591 |
| GLORI HEK293T STM2457 | 3 694 | 7 451 | 4 624 902 | 5 174 | 325 065 | GSM6432635 |
| GLORI HEK293T FTO | 3 694 | 7 737 | 4 624 616 | 5 597 | 324 642 | GSM6432593 |
| GLORI HeLa | 2 500 | 21 267 | 3 089 235 | 17 711 | 203 493 | GSM6432595 |
| GLORI HeLa hypoxia | 2 500 | 16 556 | 3 093 946 | 13 395 | 207 809 | GSM6432598 |
| GLORI HeLa siMETTL3 | 2 500 | 10 765 | 3 099 737 | 9 335 | 211 869 | GSM6432607 |
| GLORI HeLa siMETTL14 | 2 500 | 10 560 | 3 099 942 | 9 140 | 212 064 | GSM6432611 |
| GLORI HeLa siYTHDF2 | 2 500 | 11 800 | 3 098 702 | 10 245 | 210 959 | GSM6432623 |
| GLORI HeLa STM2457 | 2 500 | 3 740 | 3 106 762 | 2 791 | 218 413 | GSM6432635 |
| GLORI HEK293T | 263 | 3 542 | 323 183 | 2 929 | 20 174 | GSM7438377 |
| miCLIP HEK293T | 263 | 642 | 32 3183 | 518 | 21 667 | GSM1556679 |
| GLORI HEK293T | 1 259 | 9 188 | 1 502 545 | 7 852 | 100 975 | GSM7438377 |
| miCLIP HEK293T | 301 | 690 | 347 574 | 554 | 23 955 | GSM1556679 |

**Table 5:** IAV Dataset specifications of methylated sites.

| IAV | PB1 | PB2 | PA | HA | NP | NA | M | NS |
| --- | --- | --- | --- | --- | --- | --- | --- | --- |
| +sstr | 10 | 5 | 4 | 16 | 23 | 19 | 5 | 12 |
| -sstr | 4 | 2 | 1 | 3 | 6 | 3 | 4 | 2 |
| Accession | AF389115 | AF389116 | AF389117 | AF389118 | AF389119 | AF389120 | AF389121 | AF389122 |

MEMAR-XC and −18% for GLORI MEMAR-XT. This confirms the benefit to evaluate models with GLORI experiments due to higher absolute quantification. Performance across GLORI display better generalisation over all candidate sites in the transcriptome, without having to artificially fix the imbalance in data. In this setting, relative performance pattern (panel **A**) is conserved, with MEMAR-XT trained on GLORI still outperforming the remaining models.

An interesting additional feature of GLORI data compared to miCLIP is the access to the stoichiometry of methylation at each site. This gives the opportunity to predict a true site methylation frequency, rather than a probability to binary classify sites in methylated or non-methylated. In Fig 4**C**, the average stoichiometries of twelve DRACH motifs are compared with their average predicted probability, assigned by models (see Stoichiometry analysis on DRACH motifs). Linear regression were plotted for the three models. As expected, MEMAR trained with GLORI data, and thus provided with information about the stoichiometry, is able to predict average methylation frequency better than all counterparts. The linear regression shows a strong agreement, due to the closer-to-unitary slope, although with a negative offset (*y* = −0.149 + 0.998·*x*). The model is therefore able to greatly reproduce the experimentally observed relative importance between motifs. DeepSRAMP, is, by design, not trained with stoichiometry information, and displays a fitted line with a very strong offset and an important deviation, as expected, to the unitary slope (*y* = 0.643 + 0.257 *·x*). The model therefore tends to assign scores which are generally higher than actual stoichiometry, showing only small variations between motifs. Surprisingly, MEMAR-XC trained with miCLIP, without any stoichiometry information, similar to DeepSRAMP, shows the smallest offset amongst all models and a comparable slope to DeepSRAMP (*y* = −0.028 + 0.252*· x*).

The simple parameters of MEMAR (Appendix A4) and the numerical implementation (paragraph Training and Appendices A2,A3) allow a fast training of 1 min 14 s on a M1 Apple Silicon chip using one core (on the whole GLORI HEK293T methylated and non-methylated datasets) and evaluation of the methylation probabilities over test sets. Over 200 randomly sampled human transcripts, MEMAR was able to score transcripts in less than 100 milliseconds, depending directly on the size of the transcript. By comparison, DeepSRAMP needs between 25 and 225 seconds to perform the same operation, therefore a 1000 fold factor (see Performance evaluation against DeepSRAMP paragraph and Suppl. Fig. S8). This capacity of MEMAR enables to test the whole human transcriptome rapidly.

### MEMAR generalization over influenza A virus

In this section we apply MEMAR to an influenza A virus strain to further evaluate its generalizability beyond human transcripts (see paragraph IAV Prior and testing for datasets construction details). To this end we will consider MEMAR-XT performance on five positive strands (+sstr segments). MEMAR-T is used over the eight genomic strands (-sstr segments, vRNA), since the negative sensed counterparts do not have a coding region to project positions along the meta-transcript as defined in ***m***^**6**^***A*** position-dependence along the transcript. In Fig. 5**A** the AUPRC over all DRACH motifs, in +sstr, reaches 0.79, and 0.49 on -sstr. This great difference relies once again on the data imbalance. Indeed, positive strands from experimental data [58] were more methylated than negative strands (28.4% against 13.6%). In both PRCs, the threshold set over the validation set in human genomes (see paragraphs Validation and Threshold) generalizes well with respect to the maximum F1 scores. Moreover, the results are compared to DeepSRAMP [39], which over the same settings, underperforms compared to both MEMAR classes, with AUPRC values of 0.54 and 0.33 over +sstr and vRNA respectively. DeepSRAMP lower performance on viral sequences are expected as the model has been designed and optimized for mammalian transcripts. In Fig. 5**B**, the resulting *m*^6^*A* predicted probabilities are shown, on the +ssRNA (upper graphs) or -ssRNA (lower ones) for five IAV segments (PB2, PB1, HA, NP, NS), with the remaining three (PA, NA and M) shown in Suppl. Fig. S9 (see also Suppl. Tab. S3 for segment-imbalance details). A first difference arises between the polymerase segments, having very few predicted methy-lated sites with low accuracy, and the remaining segments with much more predicted methylations and higher accuracy. MEMAR is not able to reliably predict methylated sites on the polymerases segments. Additionally to the known performance-degradation due to the lower percentage of methylated sites, poor accuracy can be also heightened by the *prior* (see Suppl. Tab. S2) leading to lower predicted methylation probabilities on these segments. The *prior* was, indeed, determined on an independent experimental set, at low resolution [41] (see IAV Prior and testing), with very low detected methylation signal for polymerase sequences. On the contrary, the remaining segments have a much higher methylation signal, allowing to include more motifs in the methylated class and to greatly improve model’s prediction, with NS reaching the highest performance.

**Figure 5:**
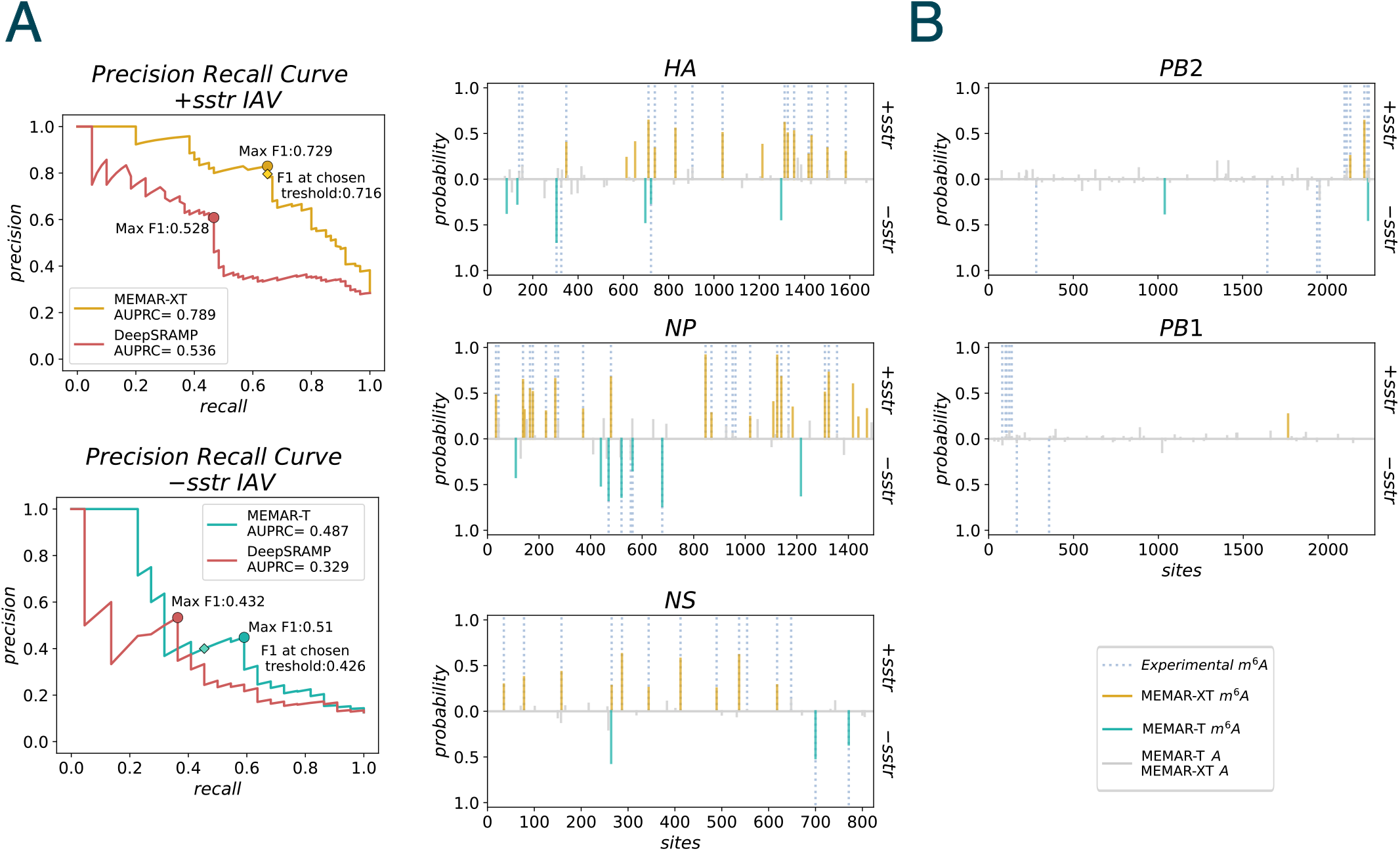
MEMAR predictions on out-of sample influenza A viral segments. Performance of prediction of *m*^6^*A* on DRACH motifs comparing MEMAR-XT (trained on GLORI on human HEK293T cell line [34]), with transcript-dependent IAV prior, and DeepSRAMP. **A top)** MEMAR-XT Precision-Recall Curve over positive IAV strands shows a high AUPRC. The optimal F1 score and the one obtained from the threshold in human validation dataset, are very close and outperform DeepSRAMP [39] predictions. **A bottom)** MEMAR-T performance on negative strands, with smaller *m*^6^*A* ratio and without the position-dependent *prior*, shows worse performance. AUPRC and Maximal F1 (obtained at a threshold different from the validation one in human dataset) are above DeepSRAMP values. **B)** Landscape of predicted *m*^6^*A* sites over two polymerase segments (PB2 and PB1), one subtype segment (HA) and NP and NS against experimental *m*^6^*A*s [58]. Positive strand results are shown on upper parts while negative on lower ones. Polymerase segment predictions show MEMAR-XT (+sstr) and MEMAR-T (-sstr) weakest performance, while accurate predictions are obtained for the remaining segments, amongst which NS yields best predictions.

In conclusion, MEMAR trained with human transcripts *m*^6^*A* sites has a good generalization potential on IAV strains infecting human cells. Adjusting the *prior* values is enough to reach good performance, with a strong agreement between the experimentally characterized *m*^6^*A* sites in the viral segments and the sites with the highest scores in non-polymerase segments.

### A two-state dynamical model provides interpretation of performance scaling

In this section we discuss the model generalization in human and viral data considered above by providing additional supporting arguments with a biophysical two-state model. To this end we consider MEMAR-XT applied to test subsets from GLORI data in HEK293T [34], chosen at different *m*^6^*A* percentages from 1% to the full balanced 50% case (for the other model classes see Suppl. Fig. S11). For each *m*^6^*A* imbalance value, 30 random subsets are taken to compute average AUPRCs with their standard deviations. The resulting curve depicts the performance at increasing methylated percentage. In Fig. 6**A** **top**, the curve shows a monotonic increasing behaviour, while reaching a saturation plateau of about 0.85 AUPRC around 35%. The AUPRC values for the different test sets that were considered in previous sections are shown against this curve. Unaltered conditions in human and viral sequences and altered ones which are known to not *directly* target methylation editing are shown with triangles and include HEK293T and HeLa in control conditions, siYTHDF2, and hypoxia, as well as HEK293T with miCLIP and IAV viral sequences infecting A549 cell line. Despite coming from very different settings compared to GLORI HEK293T in control condition, all of these AUPRCs fall on the numerical curve that depends only on the methylation ratio in the test set, with a small deviation for miCLIP. As already discussed, this might be mainly linked to the the antibody dependence of this experimental technique. Surprisingly, IAV sequences are also well explained and generalized, despite the different cell line, transcripts and experimental method (still based on antibodies). The scaling of the model with *m*^6^*A* percentage is therefore sufficient to explain the performance gain/loss in these cases. Such scaling is an *universal* property that can be explained by a two-state model, describing the methylation probability as the result of the dynamical binding and unbinding of writers and erasers in time. Based on this description we define the two sets of motif-dependent kinetic constants: 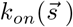 and 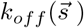 whose values can be obtained from the simplest MEMAR-C model (see paragraph Two-state dynamical model). Using this two-state model, the AUPRC and the methylation ratio can be expressed as a function of a uniform rescaling of the de-methylation rate 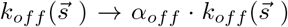; by decreasing *α*_*off*_, the percentage of methylated sites increases. We can also vary the time parameter in the two state model; at short time scale, the methylation is indeed described by an out-of-equilibrium process (see paragraph AUPRC from the two-state model and Appendix A6).

**Figure 6:**
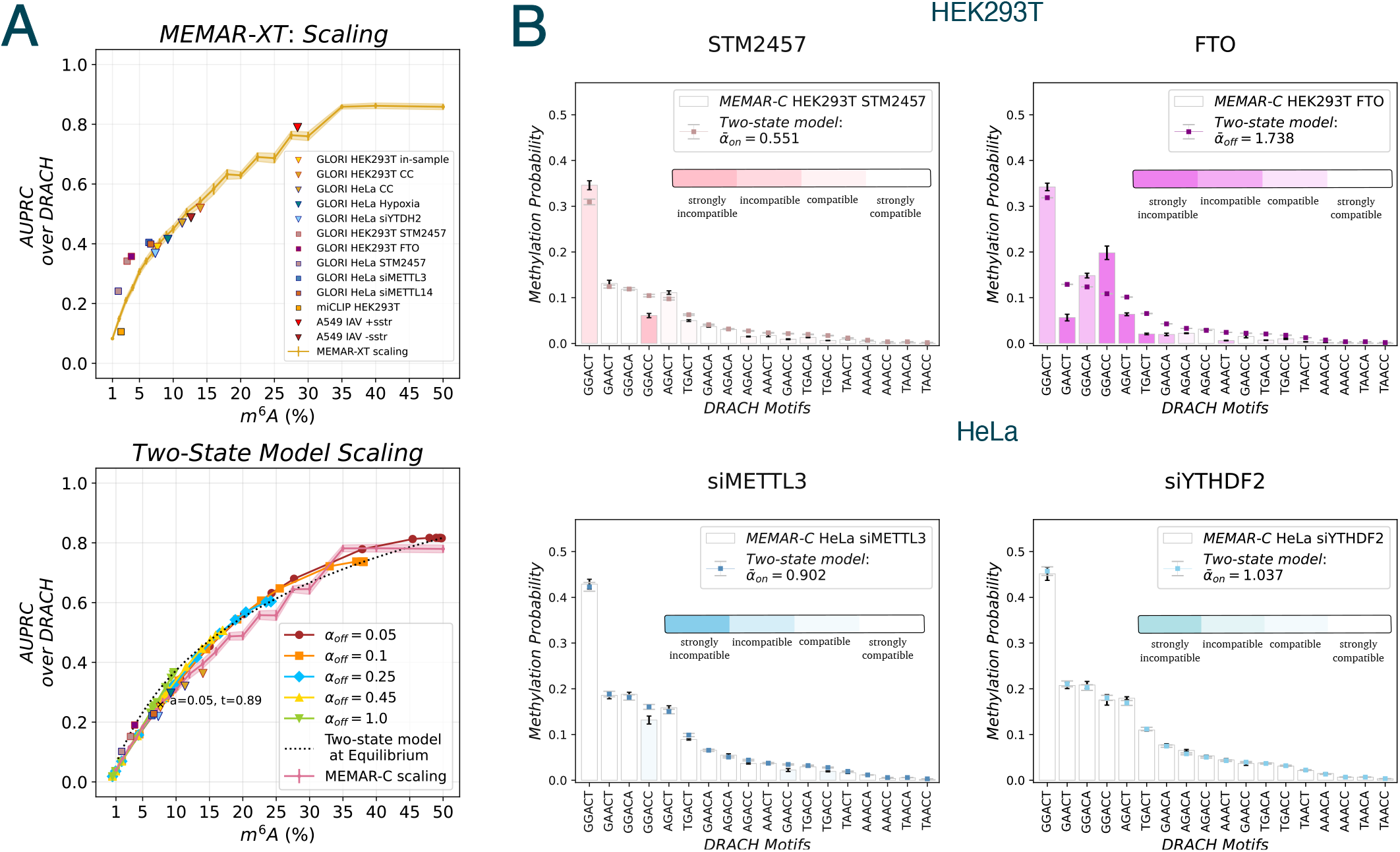
Numerical and theoretical scaling of MEMAR performance with respect to data imbalance and variations in altered methylation conditions. **A top**) MEMAR-XT AUPRC performance on test-sets on DRACH motifs in HEK293T at different *m*^6^*A* percentages. AUPRC of datasets, at their unaltered imbalance values, are shown with triangle if the corresponding experiments do *not directly* target *m*^6^*A* writers or erasers expressions. All scattered squares points represent datasets where *m*^6^*A* editing is *directly* targeted, either by writers knock-down or over-expression of erasers. **A bottom**) Theoretical scaling of MEMAR-C from the two-state dynamical model. The curves display different AUPRC and *m*^6^*A* percentage as a function of time and at different de-methylation rates overall collapsing into a single curve representing equilibrium points. The parameter *α*_*off*_ represents a multiplication factor of the de-methylation binding constant to modify the percentage of methylated sites at equilibrium. All MEMAR-C predictions on GLORI dataset are shown against the curves. **B**) Comparison between MEMAR-C methylation probability inferred for a given *m*^6^*A*-altered dataset and the two-state model obtained from MEMAR-C and rescaled uniformly either increasing the eraser (*k*_*off*_) or decreasing the writer (*k*_*on*_) binding constants. Single-motif methylation probabilities obtained by the rescaled two-state model are coloured by the degree of compatibility with MEMAR-C directly inferred on the altered condition data. We refer to strongly compatible, compatible, incompatible, or strongly incompatible models, if the absolute difference between the two models prediction *σ*, is respectively: *σ* ≤ 0.005, 0.005 *< σ* ≤ 0.015, 0.015 *< σ* ≤ 0.03, *σ >* 0.03.

The variations of both time, until the equilibrium condition, and of kinetic rates give the theoretical AUPRC curves, displayed in Fig. 6**A** **bottom**. These curves tend to overall collapse into a single one, made by all equilibrium points. This collapse allows a dual description of the methylation process. Indeed, considering the standard GLORI dataset for inference [34], its percentage of DRACH methylated sites, which is equal to 7.66%, *m*^6^*A*s can be described either at equilibrium (*t* → ∞) or, in a an out-of equilibrium description at a finite time *t* (for instance, choosing *α*_*off*_ = 0.05 and computing the time which guarantee the same percentage of methylated sites, *t* = 0.885 in the case considered). Since this two-state model neglects any position/transcript-dependence, its scaling can be compared to the one obtained from MEMAR-C on test subsets at different *m*^6^*A* imbalance percentages, showing a high agreement. All MEMAR-C predictions on GLORI data amongst the same set of transcripts, are shown in the plot and overall are well described.

Hereafter, we exploit the two-state model at equilibrium to reproduce the methylation probability in altered conditions. We focus therefore on the squared points corresponding to altered conditions by FTO and STM2457 in HEK293T as well as STM2457, siMETTL3 and siMETTL14 in HeLa. The simplest way of describing inhibition or silencing of methylation writer is by an uniform decrease *α*_*on*_ of the methylation kinetic constant 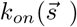or by a constant increase *α*_*off*_ of the de-methylation kinetic constant 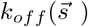 for FTO from the two-state model. After focusing on DRACH motifs only (see Marginalization), we obtained the average uniform parameter 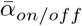which minimizes the difference between the two-state model and MEMAR-C methylation probability re-inferred from each given dataset (see paragraph Kinetic parameter variations in altered-methylation experiments and Appendix A6).

Fig. 6**B** displays motif-dependent methylation probabilities, obtained from the kinetic model after the uniform rescaling, compared to the ones obtained by MEMAR-C directly inferred from the altered condition dataset. We found that the uniform rescaling consistently predict methylation probabilities in altered conditions apart from few motifs in specific conditions. In particular for STM2457 in HEK293T, and to a lesser extent in HeLa (see suppl. Fig. S12), the motif *GGACT* appears to be less impacted by methylation inhibition than predicted with the uniform rescaling; while the motif *GGACC* appears to be more affected in STM2457 and less in FTO. In FTO conditions, the uniform rescaling is also non satisfied for other motifs in particular *GAACT*, AGACT and *TGACT*, which are more affected by de-methylation than what predicted by the uniform hypothesis.

Similar motif-dependent anomalies are obtained for FTO and STM2457 when applying an uniform rescaling from MEMAR-C directly inferred on control-condition, deep sequencing data in HEK293T (and equivalently in HeLa) [33], as shown in Suppl. Fig. S13 and S14. The anomalies of methylation probabilities in the two-state binding constants allow us to correctly predict the increase of the proportion of first-ranked motifs amongst methylated sites, as observed in Fig. 2**D**, see Suppl. Fig. S15.

For the case of silencing of METTL3 (Fig. 6**B**) and METTL14 (Suppl. Fig. S12) the uniform decrease of the binding constants (*α*_*on*_ *<* 1) well describes the altered condition (see also Suppl. Fig. S14). Despite the validity of the uniform rescaling, the non linearity of the two-state model explains, since *α*_*on*_ *<* 1, a small higher proportion of the first-ranked motif with respect to the control condition (Suppl. Fig. S15).

We lastly consider altered conditions in which the readers’ expressions are silenced. In this case, we expect the statistics or the recruitment of the writers to be unaltered. Since the model actually neglects parameters linked to the reader process contributions, we will uniformly lower 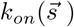 over all the motifs to take into account the lowering of the methylation amount. Fig 6**B** shows an overall strong agreement to the uniform rescaling of the binding constants. Similar plots are shown for all remaining conditions in Suppl. Fig. S12, providing strong agreement to the uniform hypothesis. This is further confirmed studying variations of the two-state model from control conditions with deep-sequencing (Suppl. Fig. S14). Moreover, the small uniform decreases of 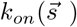 do not alter the proportion of the first-ranked motif in siYTHDF2 (Suppl. Fig. S15).

## Discussion

MEMAR is a maximum entropy model built to reproduce the pattern of motifs which are methylated in a transcript, resulting from the interplay between *m*^6^*A* writers and erasers. This simple model can be quickly inferred on one core of a M1 Apple Silicon processor (3.20 GHz) in about one minute, and used to test sequences in a time scale of milliseconds. MEMAR, trained on GLORI data [34], one of the most powerful experimental method for *m*^6^*A* mapping which also includes stoichiometry information of methylated sites, provides reliable frequency of methylation. Moreover the model has good generalization properties by simply adjusting the *prior* knowledge of *m*^6^*A* levels per transcript, for experiments with partial or lower quality data, as reflected by the agreement with the cross-prediction performance over miCLIP [26] data or IAV methylation maps [58].

### MEMAR modules

MEMAR architecture is focused on three aspects of methylation: unveiling the 8-mer motifs surrounding the adenosine which may lead to methylations, the average distribution of methylation along the meta-transcript consisting of the 5’UTR-CDS-3’UTR, and the different level of methylation amongst transcripts. MEMAR-C, our simplest model class based only on motif statistics, extending predictions beyond the -2/+2 position surrounding an adenosine up to -4/+4 nucleotides, is already able to increase performance compared to the baseline DRACH model. This result is in agreement with the finding of [61], showing that mutations in the -4/+4 nucleotide range directly affect the methylation of the central adenosine in *Saccharomyces* species, particularly at the -4, -2, -1, + 1, and + 4 positions. This supports the importance of choosing a larger window of nucleotides.

The addition of a non-constant *prior* between transcripts allows to further increase performance between MEMAR-C/XC and MEMAR-T/XT. It was recently experimentally proven that the uneven distribution of *m*^6^*A* between transcripts could be the mechanical consequence of different dwelling times of transcripts inside the nucleus [10]. If the methylation happens post-transcriptionally in the nucleus, then transcripts spending more time in the nucleus should have higher methylation levels. Therefore, the *prior* knowledge of *m*^6^*A* levels per transcript could indirectly incorporate this unknown mechanism, distinguishing genes with different methylation levels. As a future research direction, the parameter corresponding to the time of two-state model could be directly fitted from a deeper knowledge of the transcript-dependent dwelling times.

MEMAR-XC and MEMAR-XT predicted position distributions of methylation sites along transcripts agree to the well-defined experimental one [24] with *m*^6^*A* virtually absent in the 5’UTR part of transcripts and enriched around the STOP codon. The lack of methylation in the 5’UTR part (predicted by all MEMAR classes) directly arises from the low number of DRACH motifs in this region, apparently counter-selected by evolution. Moreover, recent works have shown how codon Exon-Junction Complexes, which are bound to transcripts in coding regions, prevent methyl-transferase to methylate sites in their proximity [62, 63], impacting *m*^6^*A* density in CDS. Overall, MEMAR classes provide predictions tailored on the knowledge of the testing sequence. For instance, if only the nucleotide sequence is known, for a given transcript, MEMAR-C should be used. If the coding region is additionally known, then MEMAR-XC is suggested for higher accuracy. Lastly, if any prior information regarding the transcript-dependent methylation imbalance can be obtained, MEMAR-T and MEMAR-XT are advised.

### Features not taken into account by MEMAR

The simple architecture of the model does not take into account any structural information. As a matter of fact, it has been suggested that the RNA secondary and tertiary structures could have an influence on the methylation level, based on the intuition that sites in single strands are more accessible to the writer complex than sites in double-strand RNA. Yet this structural effect on methylation is not experimentally proven. Some authors have claimed that about 30% of the differentially methylated sites are not due to mutations directly in the DRACH motif, but rather in neighbouring residues that indeed favour an open conformation of RNA and a higher methylation rate based on structural simulations [61]. Another study has argued that inhibition of methylation induces the closing of RNA loops according to icSHAPE experiments, which shows that the open-conformation is not a pre-requisite for methylation [64]. In that case, *m*^6^*A* could destabilize RNA secondary structures to expose binding motifs to proteins, hence the positive correlation between *m*^6^*A* and open structures [65]. It would be of particular interest, in future studies, to better understand the role of the structure and its connection to the *m*^6^*A* deposition especially in the specific case of viruses. This could potentially increase model performance by adding possible filters linked to inaccessible regions, or additional analysis to evaluate destabilization/stabilization effect of *m*^6^*A* on viral sequences. At last, MEMAR neglects any information relative to the position of splicing sites, the proximity effect of DRACH methylated motifs [33], or additional broader contextual information [66].

### Experimental techniques differences

MEMAR uses high quality experimental methylation profiles in the training data from GLORI experiments. Despite some limitations regarding adenosine resisting the chemical treatment and being detected as *m*^6^*A* [34], it is one of the most trustful and powerful method existing at the moment [67]. Yet, only 58% of miCLIP detected sites are retrieved by GLORI on the same cell lines [33]. This is mainly due to the known issues of antibody-reliant technique, the low reproducibility between replicates [68], and a high rate of falsely detected sites by miCLIP. This makes it crucial to benchmark predictors on data obtained by unbiased experiments and to avoid mixed datasets from multiple experiments in test sets. As we have shown, higher quality data are useful to understand model generalization from or to limited or low quality data. Third-generation sequencing, which rely neither on antibody/enzymes nor chemical treatments, may provide useful data in the future to assess the power of existing classification algorithms.

### Comparison with DeepSRAMP

DeepSRAMP [39] is a deep-network model trained on CLIP data. It uses a higher number of parameters than MEMAR and reaches AUPRC between 0.379 to 0.869 in human transcripts depending on the tissue type. This complex architecture remains partly interpretable, as authors have shown how to extract the impact of genomic features (such as the distance to splicing sites or the the 5’ and 3’ UTR of transcripts) from model parameters, although it requires a lot of training data and it lacks adaptable parameter to easily generalize predictions. Moreover, DeepSRAMP testing time is of a factor 1000 higher than MEMAR (seconds against milliseconds) which makes it difficult to apply for a vast analysis of sequences, such as required for the study of methylation maps in viral sequences to compare different subtypes and clades.

DeepSRAMP tends to assign overall-high probabilities to real methylated sites (0.77 on average). While the predictor is not expected to well capture stoichiometry, due to CLIP training, the architecture strong predictions seem to lack the methylation variability. In fact, the absolute quantification of methylation stoichiometry in HEK293T, enabled by GLORI data, gives a median of 0.4 frequency of methylation for the methylated sites with only 38% of sites above 0.5 [33]. Nonetheless, MEMAR trained on miCLIP data, without any stoichiometry information likewise, predicts average methylation frequency of 0.18. This avoids polarization into high-scored methylations which, despite an overall under-estimation, better reflects their stochastic nature. Moreover, we have shown how DeepSRAMP does not benefit from high quality GLORI test data, as opposed to MEMAR-XC. This likely indicates how the model complexity tends to over-fit to some artefactual features linked to the experimental technique.

On HEK293T miCLIP test data, DeepSRAMP’s authors have reported an higher AUPRC of 0.6 against 0.4 in our own evaluation. This variation is due to a few differences with authors original test sets. In our case, we have restricted predictions over a set of 301 test genes, to exclude transcripts overlapping with MEMAR training. We also chose one single experiment with mapped methy-lated sites over only the longest isoforms, while randomly selecting *m*^6^*A* sites at fixed 10% imbalance without any bias in the position.

Lastly, as expected from the lower amount of features inferred from human transcripts and thanks to the *prior* flexibility, MEMAR is better transferable to viral datasets than DeepSRAMP.

### MEMAR generalization on different test data and altered *m*^6^*A* conditions

Model performance were shown to be impacted by several factors, due to experimental biases or biological features which modify *m*^6^*A* landscapes, changing overall methylation percentage. In order to overcome the *m*^6^*A* imbalance, and to compare different datasets in a fairer way, we have in several cases fixed the imbalance between methylated and non-methylated sites to 10 − 90% in the test dataset. In altered conditions, we noticed performance improvements as a consequence of different *m*^6^*A* profiles, in particular due to the higher proportion of MEMAR highest ranked motif *GGACT*.

Subsequently, in the last section, we have compared out-of-sample test datasets with their own methylation percentage, towards the expected numerical scaling of MEMAR performance when rebalancing the in-sample test data. The universality of the numerical scaling curve is explained thanks to a two-state biophysical model between *m*^6^*A* writer and erasers proteins. We proved that a uniform rescaling of the binding constants of *m*^6^*A* writers, and therefore a simple variations in protein concentrations, was able to describe methylation profiles for silencing of METTL3 and METTL14 writers. Although, non-uniform effects over the higher-ranked motifs were observed for FTO and STM2457 conditions. These results ultimately suggest that these conditions may not solely impact the writer/eraser abundance, but also affect the deposition of methylation at specific motifs.

Some experimental conditions, not targeting directly methylation writer or erasers, can trigger complex signalling pathways which could alter differentially *m*^6^*A* [8]. For instance, it has been experimentally observed that 5% to 11% of the sites are differentially up or down methy-lated in conditions such as hypoxia or heat shock [33]. In this article, we have shown, in particular, how the dynamical alteration induced by hypoxic and siYTHDF2 conditions does not impact methylation profiles over the motifs, but solely abundance and/or position, as an overall uniform rescaling of the writer binding constants is sufficient to explain the statistics of methylated motifs.

### Prediction of methylation in viral sequences

The genome of influenza A is known to be methylated since the work of Krug, Morgan and Shatkin in 1976 [69]. As viruses such as IAV hijack the host’s machinery during their replication cycle, they exploit the METTL3/14 *m*^6^*A* writer complex. We therefore expect that the methylation pattern in viruses in infected human cells should be similar to the human methylation profile. It is known that *m*^6^*A* methylation in IAV is involved in gene expression regulation, from stability, translation efficiency or secondary structure shifts, with important effect on viral replication and potential antiviral innate immune response [70]. The A385 *m*^6^*A* site of the NS segment, is conserved across human and avian strains, and is involved in the splicing equilibrium between the NS1 and NS2 proteins through the recruitment of the reader YTHDC1 [71]. Other reader proteins are also able to bind to other *m*^6^*A* sites on the mRNA and vRNA of the HA, NP, NA and M segments, and the overexpression of YTHDF2 enhanced the viral replication [41, 72]. The inhibition of methylation in the human lung epithelial cell line A549 has a dramatic effect on the expression of the NS1, NP and M2 proteins, and a virus containing mutated HA segments with a lower amount of methylated sites has a lower amount of expressed HA protein and a lower pathogenicity compared to the wild type strain [41]. We demonstrated that MEMAR can be used to predict with high accuracy the methylation sites of influenza A virus over the positive strands of non-polymerase segments. Generalization over negative strands shows lower performance whilst being perfectly explained by MEMAR scaling of performance due to the low amount of retrieved *m*^6^*A* compared to non-methylated adenosine. These results have been achieved with an adapted methylation *prior*, a strategy that we had shown to be efficient to improve performance over miCLIP data. The overall *prior* is fixed using as few data as three influenza sequences (PA, M and NA segments).

It has been reported that the amount of methylation per segment is highly variable in IAV. The HA and NA segment bear about 7 to 8 methylation sites [73], while other segments such as PB1, PA or M have a much lower amount of *m*^6^*A* sites, below 1 or 2 [73, 41]. Interestingly, MEMAR results are in agreement with these experimental evidence, predicting 6 sites for HA, or 0 site for PB1. We also predict much fewer sites on the negative genomic strands than on the positive, mRNA transcripts, in accordance with [41]. If this observation is specific to IAV or a more general feature of negative RNA virus is of particular interest. We can therefore expect that the amount of *m*^6^*A* on vRNA and mRNA on the different segments of Influenza A virus are “hard-coded” within the sequence itself, and could evolve accordingly to maximize the replication and transmission of the virus. Further assessments with higher-resolution, antibody-independent maps are still required to confirm model generalization. In the future, MEMAR’s performance, adaptability and time-efficiency in predicting methylation sites could enable comparative studies of methylation evolution and control across thousands of already sequenced strains, helping the identification of sites that are crucial for viral pathogenicity.

## Data availability

The source code to replicate article results, as well as to apply inferred model to users personal purposes is publicly available at GITHUB: *https://github.com/IvanLec/MEMAR*.

## Supplementary Data Statement

Three Supplementary files are available: Appendices (containing MEMAR computation and the two-state biophysics model), Supplementary Figures and Supplementary Tables.

## Author Contributions Statement

**Ivan Lecce**: *Conceptualization, Methodology, Software, Validation, Formal analysis, Investigation, Resources, Data Curation,Writing-Original Draft, Writing-Review & Editing, Visualization* **Pierre-Guillaume Brun** : *Conceptualization, Investigation, Resources, Data Curation, Writing-Original Draft, Writing-Review & Editing, Visualization*. **Simon Cocco**: *Conceptualization, Methodology, Investigation,Writing-Original Draft, Writing-Review & Editing, Supervision, Project administration, Funding acquisition*.

## References

[1] Jeremy Monroe, Daniel E. Eyler, Lili Mitchell, Indrajit Deb, Abigail Bojanowski, Pooja Srinivas, Christine M. Dunham, Bijoyita Roy, Aaron T. Frank, and Kristin S. Koutmou. N1-methylpseudouridine and pseudouridine modifications modulate mrna decoding during translation. Nature Communications, 15(1):8119, Sep 2024.

[2] Brenda L. Bass and Harold Weintraub. An unwinding activity that covalently modifies its doublestranded rna substrate. Cell, 55(6):1089–1098, Dec 1988.

[3] Meng Liu, Gangqiang Guo, Pengge Qian, Jianbing Mu, Binbin Lu, Xiaoqin He, Yanting Fan, Xiaomin Shang, Guang Yang, Shijun Shen, Wenju Liu, Liping Wang, Liang Gu, Quankai Mu, Xinyu Yu, Yuemeng Zhao, Richard Culleton, Jun Cao, Lubin Jiang, Thomas E. Wellems, Jing Yuan, Cizhong Jiang, and Qingfeng Zhang. 5-methylcytosine modification by plasmodium nsun2 stabilizes mrna and mediates the development of gametocytes. Proceedings of the National Academy of Sciences, 119(9):e2110713119, 2022.

[4] Gangqiang Guo, Kan Pan, Su Fang, Lele Ye, Xinya Tong, Zhibin Wang, Xiangyang Xue, and Huidi Zhang. Advances in mrna 5-methylcytosine modifications: Detection, effectors, biological functions,and clinical relevance. Molecular Therapy -Nucleic Acids, 26:575–593, 2021.

[5] Yasuhiro Furuichi. Discovery of m7g-cap in eukaryotic mrnas. Proceedings of the Japan Academy, Series B, 91(8):394–409, 2015.

[6] Juyeong Hong, Kexin Xu, and Ji Hoon Lee. Biological roles of the RNA m6A modification and its implications in cancer. Experimental & Molecular Medicine, 54(11):1822–1832, November 2022.

[7] Jiangle Liu, Xueqing Yan, Hao Wu, Ziqin Ji, Ye Shan, Xinyan Wang, Yunfan Ran, Yichen Ma, Caitao Li, Yuchao Zhu, Ruichu Gu, Han Wen, Chengqi Yi, and Peng R. Chen. RNA codon expansion via programmable pseudouridine editing and decoding. Nature, 643(8074):1410–1420, July 2025.

[8] P. Cody He and Chuan He. m6a rna methylation: from mechanisms to therapeutic potential. The EMBO Journal, 40(3):EMBJ2020105977, Jan 2021.

[9] Shichen Su, Shanshan Li, Ting Deng, Minsong Gao, Yue Yin, Baixing Wu, Chao Peng, Jianzhao Liu, Jinbiao Ma, and Kaiming Zhang. Cryo-EM structures of human m6A writer complexes. Cell Research, 32(11):982–994, September 2022.

[10] Peng Tang, Jiayi Yang, Zonggui Chen, Chen Du, Yang Yang, Haiping Zhao, Li Huang, Guangnan Li, Feiyan Liu, Bei Dong, Ting Shan, Xi-chen Bao, and Yu Zhou. Nuclear retention coupled with sequential polyadenylation dictates post-transcriptional m6A modification in the nucleus. Molecular Cell, 84(19):3758–3774.e10, October 2024.

[11] Jinbo Huang, Xu Dong, Zhou Gong, Ling-Yun Qin, Shuai Yang, Yue-Ling Zhu, Xiang Wang, Delin Zhang, Tingting Zou, Ping Yin, and Chun Tang. Solution structure of the rna recognition domain of mettl3-mettl14 n6-methyladenosine methyltransferase. Protein & Cell, 10(4):272–284, 03 2018.

[12] Eva Schöller, Franziska Weichmann, Thomas Treiber, Sam Ringle, Nora Treiber, Andrew Flatley, Regina Feederle, Astrid Bruckmann, and Gunter Meister. Interactions, localization, and phosphorylation of the m6 A generating METTL3–METTL14– WTAP complex. RNA, 24(4):499–512, April 2018.

[13] Huaqing Yan, Liqi Zhang, Xiaobo Cui, Sinian Zheng, and Rubing Li. Roles and mechanisms of the m6A reader YTHDC1 in biological processes and diseases. Cell Death Discovery, 8(1):237, May 2022.

[14] Mathieu N Flamand and Kate D Meyer. m6A and YTHDF proteins contribute to the localization of select neuronal mRNAs. Nucleic Acids Research, 50(8):4464–4483, May 2022.

[15] Deepak P. Patil, Chun-Kan Chen, Brian F. Pickering, Amy Chow, Constanza Jackson, Mitchell Guttman, and Samie R. Jaffrey. m6A RNA methylation promotes XIST-mediated transcriptional repression. Nature, 537(7620):369–373, September 2016.

[16] Loic Broix, Rohini Roy, Belal Shohayeb, Ikumi Oomoto, Hiroki Umeshima, and Dan Ohtan Wang. m6A RNA methylation-mediated control of global APC expression is required for local translation of β-actin and axon development. Cell Reports, 44(6):115727, June 2025.

[17] Xiulin Jiang, Baiyang Liu, Zhi Nie, Lincan Duan, Qiuxia Xiong, Zhixian Jin, Cuiping Yang, and Yongbin Chen. The role of m6A modification in the biological functions and diseases. Signal Transduction and Targeted Therapy, 6(1):74, February 2021.

[18] Dermot Harnett, Mateusz C. Ambrozkiewicz, Ulrike Zinnall, Alexandra Rusanova, Ekaterina Borisova, Amelie N. Drescher, Marta Couce-Iglesias, Gabriel Villamil, Rike Dannenberg, Koshi Imami, Agnieszka Münster-Wandowski, Beatrix Fauler, Thorsten Mielke, Matthias Selbach, Markus Landthaler, Christian M. T. Spahn, Victor Tarabykin, Uwe Ohler, and Matthew L. Kraushar. A critical period of translational control during brain development at codon resolution. Nature Structural & Molecular Biology, 29(12):1277–1290, December 2022.

[19] Frederic Truffinet, Alejandro Arco-Hierves, Hosnia Shalabi, Juliette Pascaud, Paul Mazet, Elodie Rivière, Sacha E. Silva-Saffar, Lucilla Fabbri, Sophie Leboucher, Laetitia Besse, Cedric Messaoudi, Aurore Attina, Alexandre David, Stephan Vagner, Gaetane Nocturne, Xavier Mariette, and Rami Bechara. m6A RNA methylation controls salivary gland epithelial cell function and has a protective role in Sjögren’s disease. Annals of the Rheumatic Diseases, 84(5):746–759, May 2025.

[20] Andrew A. Guirguis, Yaara Ofir-Rosenfeld, Kathy Knezevic, Wesley Blackaby, David Hardick, Yih-Chih Chan, Ali Motazedian, Andrea Gillespie, Dane Vassiliadis, Enid Y.N. Lam, Kevin Tran, Byron Andrews, Michael E. Harbour, Lina Vasiliauskaite, Claire J. Saunders, Georgia Tsagkogeorga, Aleksandra Azevedo, Joanna Obacz, Ewa S. Pilka, Marie Carkill, Laura MacPherson, Elanor N. Wainwright, Brian Liddicoat, Benjamin J. Blyth, Mark R. Albertella, Oliver Rausch, and Mark A. Dawson. Inhibition of METTL3 Results in a Cell-Intrinsic Interferon Response That Enhances Antitumor Immunity.Cancer Discovery, 13(10):2228–2247, October 2023.

[21] Akiko Ogawa, Chisae Nagiri, Wataru Shihoya, Asuka Inoue, Kouki Kawakami, Suzune Hiratsuka, Junken Aoki, Yasuhiro Ito, Takeo Suzuki, Tsutomu Suzuki, Toshihiro Inoue, Osamu Nureki, Hidenobu Tanihara, Kazuhito Tomizawa, and Fan-Yan Wei. N6methyladenosine (m6A) is an endogenous A3 adenosine receptor ligand. Molecular Cell, 81(4):659– 674.e7, February 2021.

[22] Akiko Ogawa, Satoshi Watanabe, Iuliia Ozerova, Allen Yi-Lun Tsai, Yoshihiko Kuchitsu, Harrison Byron Chong, Tomoyoshi Kawakami, Jirio Fuse, Wei Han, Ryuhei Kudo, Tomoki Naito, Kota Sato, Toru Nakazawa, Yasunori Saheki, Akiyoshi Hirayama, Peter F Stadler, Mieko Arisawa, Kimi Araki, Liron BarPeled, Tomohiko Taguchi, Shinichiro Sawa, Kenji Inaba, and Fan-Yan Wei. Adenosine kinase and ADAL coordinate detoxification of modified adenosines to safeguard metabolism. Cell, 188(22):6151–6169.e24, October 2025.

[23] Michael Kahl, Zhaofa Xu, Saravanan Arumugam, Brittany M Edens, Mariafausta Fischietti, Allen C Zhu, Leonidas C Platanias, Chuan He, Xiaoxi Zhuang, and Yongchao C Ma. m6A RNA methylation regulates mitochondrial function. Human Molecular Genetics, 33(11):969–980, May 2024.

[24] Kate D. Meyer, Yogesh Saletore, Paul Zumbo, Olivier Elemento, Christopher E. Mason, and Samie R. Jaffrey. Comprehensive analysis of mrna methylation reveals enrichment in 3’utrs and near stop codons. Cell, 149(7):1635–1646, 2012.

[25] Kai Chen, Zhike Lu, Xiao Wang, Ye Fu, Guan-Zheng Luo, Nian Liu, Dali Han, Dan Dominissini, Qing Dai, Tao Pan, and Chuan He. High-resolution n6-methyladenosine (m6a) map using photo-crosslinking-assisted m6a sequencing. Angewandte Chemie International Edition, 54(5):1587–1590, 2015.

[26] Bastian Linder, Anya V. Grozhik, Anthony O. Olarerin-George, Cem Meydan, Christopher E. Mason, and Samie R. Jaffrey. Single-nucleotide-resolution mapping of m6a and m6am throughout the transcriptome. Nature Methods, 12(8):767–772, 2015.

[27] Anya V. Grozhik, Bastian Linder, Anthony O. Olarerin-George, and Samie R. Jaffrey. Mapping m6A at Individual-Nucleotide Resolution Using Crosslinking and Immunoprecipitation (miCLIP), pages 55–78. Springer New York, New York, NY,2017.

[28] Christopher Hendra, Ploy N. Pratanwanich, Yuk Kei Wan, W. S. Sho Goh, Alexandre Thiery, and Jonathan Göke. Detection of m6a from direct rna sequencing using a multiple instance learning framework. Nature Methods, 19(12):1590–1598, Dec 2022.

[29] Kate D. Meyer. Dart-seq: an antibody-free method for global m6a detection. Nature Methods, 16(12):1275–1280, 2019.

[30] Yu Xiao, Ye Wang, Qian Tang, Lianhuan Wei, Xiao Zhang, and Guifang Jia. An elongation- and ligation-based qpcr amplification method for the radiolabeling-free detection of locus-specific n6-methyladenosine modification. Angewandte Chemie International Edition, 57(49):15995–16000, 2018.

[31] Ye Wang, Yu Xiao, Shunqing Dong, Qiong Yu, and Guifang Jia. Antibody-free enzyme-assisted chemical approach for detection of n6-methyladenosine. Nature Chemical Biology, 16(8):896–903, 2020.

[32] Charles J Sheehan, Bahjat Fadi Marayati, Janvi Bhatia, and Kate D Meyer. In situ visualization of m6a sites in cellular mrnas. Nucleic Acids Research, 51(20):e101–e101, 10 2023.

[33] Cong Liu, Hanxiao Sun, Yunpeng Yi, Weiguo Shen, Kai Li, Ye Xiao, Fei Li, Yuchen Li, Yongkang Hou, Bo Lu, Wenqing Liu, Haowei Meng, Jinying Peng, Chengqi Yi, and Jing Wang. Absolute quantification of single-base m6A methylation in the mammalian transcriptome using GLORI. Nature Biotechnology, 41(3):355–366, March 2023.

[34] Weiguo Shen, Hanxiao Sun, Cong Liu, Yunpeng Yi, Yongkang Hou, Ye Xiao, Yufei Hu, Bo Lu, Jinying Peng, Jing Wang, and Chengqi Yi. GLORI for absolute quantification of transcriptome-wide m6A at single-base resolution. Nature Protocols, 19(4):1252–1287, April 2024.

[35] Wei Chen, Pengmian Feng, Hui Ding, Hao Lin, and Kuo-Chen Chou. iRNA-Methyl: Identifying N6methyladenosine sites using pseudo nucleotide composition. Analytical Biochemistry, 490:26–33, December 2015.

[36] Kunqi Chen, Zhen Wei, Qing Zhang, Xiangyu Wu, Rong Rong, Zhiliang Lu, Jionglong Su, João Pedro de Magalhães, Daniel J Rigden, and Jia Meng. Whistle: a high-accuracy map of the human n6-methyladenosine (m6a) epitranscriptome predicted using a machine learning approach. Nucleic Acids Research, 47(7):e41–e41, 02 2019.

[37] Zhen Chen, Pei Zhao, Fuyi Li, Yanan Wang, A Ian Smith, Geoffrey I Webb, Tatsuya Akutsu, Abdelkader Baggag, Halima Bensmail, and Jiangning Song. Comprehensive review and assessment of computational methods for predicting rna post-transcriptional modification sites from rna sequences. Briefings in Bioinformatics, 21(5):1676–1696, 11 2019.

[38] Yuanpeng Xiong, Xuan He, Dan Zhao, Tingzhong Tian, Lixiang Hong, Tao Jiang, and Jianyang Zeng. Modeling multi-species rna modification through multi-task curriculum learning. Nucleic Acids Research, 49(7):3719–3734, 03 2021.

[39] Rui Fan, Chunmei Cui, Boming Kang, Zecheng Chang, Guoqing Wang, and Qinghua Cui. A combined deep learning framework for mammalian m6A site prediction. Cell Genomics, 4(12):100697, December 2024.

[40] Oliver Manners, Belinda Baquero-Perez, and Adrian Whitehouse. m6A: Widespread regulatory control in virus replication. Biochimica et Biophysica Acta(BBA) -Gene Regulatory Mechanisms, 1862(3):370–381, March 2019.

[41] David G. Courtney, Edward M. Kennedy, Rebekah E. Dumm, Hal P. Bogerd, Kevin Tsai, Nicholas S. Heaton, and Bryan R. Cullen. Epitranscriptomic Enhancement of Influenza A Virus GeneExpression and Replication. Cell Host & Microbe, 22(3):377–386.e5, September 2017.

[42] Jun’e Liu, Yan-Peng Xu, Kai Li, Qing Ye, Hang-Yu Zhou, Hanxiao Sun, Xiaoyu Li, Liu Yu, Yong-Qiang Deng, Rui-Ting Li, Meng-Li Cheng, Bo He, Jia Zhou, Xiao-Feng Li, Aiping Wu, Chengqi Yi, and Cheng-Feng Qin. The m6A methylome of SARS-CoV-2 in host cells. Cell Research, 31(4):404–414, April 2021.

[43] Yiyang Ge, Shi Tang, Tian Xia, and Chunwei Shi. Research progress on the role of RNA N6methyladenosine methylation in HCV infection. Virology, 582:35–42, May 2023.

[44] Graham D. Williams, Nandan S. Gokhale, and Stacy M. Horner. Regulation of Viral Infection by the RNA Modification N6 -Methyladenosine. Annual Review of Virology, 6(1):235–253, September 2019.

[45] Stacy M Horner and Jordan V Reaves. Recent insights into N6-methyladenosine during viral infection. Current Opinion in Genetics & Development, 87:102213, August 2024.

[46] Stacy M. Horner and Matthew G. Thompson. Challenges to mapping and defining m6 A function in viral RNA. RNA, 30(5):482–490, May 2024.

[47] E. T. Jaynes. Information theory and statistical mechanics. Phys. Rev., 106:620–630, May 1957.

[48] E. T. Jaynes. Information theory and statistical mechanics. ii. Phys. Rev., 108:171–190, Oct 1957.

[49] Benjamin D. Greenbaum, Simona Cocco, Arnold J. Levine, and Rémi Monasson. Quantitative theory of entropic forces acting on constrained nucleotide sequences applied to viruses. Proceedings of the National Academy of Sciences, 111(13):5054–5059, April 2014.

[50] Elad Schneidman, Michael J. Berry, Ronen Segev, and William Bialek. Weak pairwise correlations imply strongly correlated network states in a neural population. Nature, 440(7087):1007–1012, Apr 2006.

[51] Simona Cocco, Rémi Monasson, Lorenzo Posani, and Gaia Tavoni. Functional networks from inverse modeling of neural population activity. Current Opinion in Systems Biology, 3:103–110, 2017.

[52] Simona Cocco, Christoph Feinauer, Matteo Figliuzzi, Rémi Monasson, and Martin Weigt. Inverse statistical physics of protein sequences: a key issues review. Reports on Progress in Physics, 81(3):032601, 2018.

[53] Fengyun Yu, Yu Zhang, Chao Cheng, Wenqing Wang, Zisong Zhou, Wenliang Rang, Han Yu, Yaxun Wei, Qijia Wu, and Yi Zhang. Poly(a)-seq: A method for direct sequencing and analysis of the transcriptomic poly(a)-tails. PLOS ONE, 15(6):1–21, 06 2020.

[54] Andrea Di Gioacchino, Ivan Lecce, Benjamin D Greenbaum, Rémi Monasson, and Simona Cocco. Deciphering the code of viral-host adaptation through maximum-entropy nucleotide bias models. Molecular Biology and Evolution, 42(6):msaf127, 06 2025.

[55] Takaya Saito and Marc Rehmsmeier. The precision-recall plot is more informative than the roc plot when evaluating binary classifiers on imbalanced datasets. PLOS ONE, 10(3):1–21, 03 2015.

[56] Ning Qian. On the momentum term in gradient descent learning algorithms.Neural Networks, 12(1):145–151, 1999.

[57] N.G. VAN KAMPEN. Chapter iv markov processes. In Stochastic Processes in Physics and Chemistry (Third Edition), North-Holland Personal Library, pages 73–95. Elsevier, Amsterdam, thirdedition edition, 2007.

[58] Qian Wang, Shuai Xu, Wentao Shen, Yanli Wei, Lu Han, Zhengxiang Wang, Yingying Yu, Minxuan Liu, Junwen Liu, Guohua Deng, Hualan Chen,and Qiyun Zhu. N 6-methyladnosine of vRNA facilitates influenza A virus replication by promoting the interaction of vRNA with polymerase proteins. Proceedings of the National Academy of Sciences, 122(11):e2411554122, March 2025.

[59] Stephen F. Altschul, Warren Gish, Webb Miller, Eugene W. Myers, and David J. Lipman. Basic local alignment search tool. Journal of Molecular Biology, 215(3):403–410, 1990.

[60] Neema Agrawal, P. V. N. Dasaradhi, Asif Mohmmed, Pawan Malhotra, Raj K. Bhatnagar, and Sunil K. Mukherjee. Rna interference: Biology, mechanism, and applications. Microbiology and Molecular Biology Reviews, 67(4):657–685, 2003.

[61] Ran Shachar, David Dierks, Miguel Angel Garcia-Campos, Anna Uzonyi, Ursula Toth, Walter Rossmanith, and Schraga Schwartz. Dissecting the sequence and structural determinants guiding m6A deposition and evolution via inter-and intra-species hybrids. Genome Biology, 25(1):48, February 2024.

[62] Xin Yang, Robinson Triboulet, Qi Liu, Erdem Sendinc, and Richard I. Gregory. Exon junction complex shapes the m6a epitranscriptome. Nature Communications, 13(1):7904, Dec 2022.

[63] Miona Ćorović, Peter Hoch-Kraft, You Zhou, Sarah Hallstein, Julian König, and Kathi Zarnack. m6A in the coding sequence: linking deposition, translation, and decay. Trends in Genetics, 41(11):963–973, Nov 2025.

[64] Robert C. Spitale, Ryan A. Flynn, Qiangfeng Cliff Zhang, Pete Crisalli, Byron Lee, Jong-Wha Jung, Hannes Y. Kuchelmeister, Pedro J. Batista, Eduardo A. Torre, Eric T. Kool, and Howard Y. Chang. Structural imprints in vivo decode RNA regulatory mechanisms. Nature, 519(7544):486–490, March 2015.

[65] Nian Liu, Qing Dai, Guanqun Zheng, Chuan He, Marc Parisien, and Tao Pan. N6-methyladenosine-dependent RNA structural switches regulate RNA– protein interactions. Nature, 518(7540):560–564, February 2015.

[66] Zhengtao Luo, Liyi Yu, Zhaochun Xu, Kening Liu, and Lichuan Gu. Comprehensive Review and Assessment of Computational Methods for Prediction of N6-Methyladenosine Sites. Biology, 13(10):777, September 2024.

[67] Yuening Yang, Yanming Lu, Yan Wang, Xianghui Wen, Changhai Qi, Weilan Piao, and Hua Jin. Current progress in strategies to profile transcriptomic m6A modifications. Frontiers in Cell and Developmental Biology, 12:1392159, July 2024.

[68] Alexa B. R. McIntyre, Nandan S. Gokhale, Leandro Cerchietti, Samie R. Jaffrey, Stacy M. Horner, and Christopher E. Mason. Limits in the detection of m6a changes using merip/m6a-seq. Scientific Reports, 10(1):6590, 2020.

[69] R M Krug, M A Morgan, and A J Shatkin. Influenza viral mRNA contains internal N6-methyladenosine and 5’-terminal 7-methylguanosine in cap structures. Journal of Virology, 20(1):45–53, October 1976.

[70] Xueer Liu, Weiqiang Chen, Kangsheng Li, and Jiangtao Sheng. RNA N6-methyladenosine methylation in influenza A virus infection. Frontiers in Microbiology, 15:1401997, June 2024.

[71] Yingyin Liao, Jiayan Liu, Pui Wang, Bobo Wing-Yee Mok, and Honglin Chen. Influenza NS1 drives N6 -methyladenosine (m6 A)-mediated autoregulation of viral mRNA splicing. Emerging Microbes &Infections, 14(1):2572761, December 2025.

[72] Mahmoud Bayoumi and Muhammad Munir. Evolutionary conservation of the DRACH signatures of potential N6-methyladenosine (m6A) sites among influenza A viruses. Scientific Reports, 11(1):4548, February 2021.

[73] P Narayan, D F Ayers, F M Rottman, P A Maroney, and T W Nilsen. Unequal distribution of N6-methyladenosine in influenza virus mRNAs. Molecular and Cellular Biology, 7(4):1572–1575, April 1987.

