## Appendices for "A maximum entropy model for *m*^6^*A* prediction: from human transcripts to influenza A viral segments"

#### A1: Principle of Maximum Entropy

In this section, we look for a probability distribution $p$ which is able to describe, based on a set of chosen features, the datasets $\mathcal{D}_{methylated}$ and $\mathcal{D}_{non-methylated}$ presented in the the main article. By recalling the definition of the DNA alphabet $\mathcal{A} = \{A, C, G, T\}$ and of motif $\vec{s}$, as a vector of letters: $s_i \in \mathcal{A}/i = 1, ..., \mathcal{N} = 8$, the discreet probability distribution is a non-negative function which is normalized to one over the set of all possible motifs of length $\mathcal{N}$. By indicating the space of real numbers restricted between zero and one as $\mathcal{R}_{[0,1]}$, the probability is denoted as: $p : \mathcal{A}^{\mathcal{N}} \to \mathcal{R}_{[0,1]} \wedge \sum_{\vec{s} \in \mathcal{A}^{\mathcal{N}}} p(\vec{s}\ ) = 1$. Consequently the space of probability distribution corresponds to the ensemble $\mathcal{P}$ of all possible functions satisfying the previous conditions : $\mathcal{P} = \{p/p : \mathcal{A}^{\mathcal{N}} \to \mathcal{R}_{[0,1]} \wedge \sum_{\vec{s} \in \mathcal{A}^{\mathcal{N}}} p(\vec{s}\ ) = 1\}$. To choose a probability distribution amongst this infinite set, we will follow the Principle of Maximum Entropy, which corresponds to maximizing the information that we have about the data under a set of constraints. The information corresponds to the Shannon Entropy, which is a functional over probability distributions defined as below:

$$S : \mathcal{P} \to \mathcal{R} \ / \ S[p] = - \sum_{\vec{s} \in \mathcal{A}^{\mathcal{N}}} p(\vec{s}\ ) \cdot log(p(\vec{s}\ )) \tag{1}$$

Defining the generic observable $\mathcal{O}$ as a quantity which can be directly evaluated from a dataset $\mathcal{D}$, we then look for a probability distribution such that the average of the observable from $p$, $\langle \mathcal{O} \rangle_p$, matches the empirical average from the dataset $\mathcal{D}$, $\langle \mathcal{O} \rangle_{\mathcal{D}}$. These quantities are chosen to be the set of binary variables to describe the presence or absence of sub-motifs $\vec{S}$ of length $L = \{1, 2, 3\}$ from a given position $i$ of the motif $\vec{s}$. Formally they represent the following Kronecker Delta functions: $\delta^{\vec{S}}_{i,i+L-1}(\vec{s}\ ) = \prod_{j=i}^{i+L-1} \delta(S_{j-i+1}, s_j) = \delta(S_1, s_i) \cdot ... \cdot \delta(S_L, s_{i+L-1})$. The averages of the model will therefore be the expected values:

$$\langle \delta^{\vec{S}}_{i,i+L-1}(\vec{s}\ ) \rangle_p = \sum_{\vec{s}\ \in \mathcal{A}^{\mathcal{N}}} p(\vec{s}\ ) \cdot \delta^{\vec{S}}_{i,i+L-1}(\vec{s}\ ) = \sum_{\vec{s}\ \in \mathcal{A}^{\mathcal{N}}} p(\vec{s}\ ) \cdot \prod_{j=i}^{i+L-1} \delta(S_{j-i+1}, s_j) \tag{2}$$

Analogously, the same observables can be averaged over the dateset. Since we are interested in modeling the probability of a given sequence to be methylated taking into account real frequency signals, when the stoichiometry $\vec{w}$ of motifs is available, we will consider:

$$\langle \delta^{\vec{S}}_{i,i+L-1} \rangle_{\mathcal{D}} = \frac{\sum_{k=1}^{M} w_k \prod_{j=i}^{i+L-1} \delta(\mathcal{D}_{k,j}, S_{j-i+1})}{\sum_{k=1}^{M} w_k} = f^{\vec{S}}_{i,i+L-1} \tag{3}$$

where $M$ represents the amount of motifs contained in the given dataset $\mathcal{D}$. The latter exactly correspond to the weighted frequencies $f^{\vec{S}}_{i,i+L-1}$ of all possible mono-, di-, tri-nucleotides at each position. When the stoichiometry is unavailable, due to experimental limitations or for all non-methylated adenosine, standard frequency computation are considered (corresponding to setting all weights equal to one: $w_k$=1). We can therefore add to the Shannon Entropy at Eq. S1 the set of Lagrange multipliers $\{h^a_i\}, \{J^{ab}_{i,i+1}\}, \{K^{abc}_{i,i+1,i+2}\}$ to respect the set of constraints given by the equality:

$$\langle \delta^{\vec{S}}_{i,i+L-1}(\vec{s}\ ) \rangle_p = f^{\vec{S}}_{i,i+L-1} \tag{4}$$

The resulting constrained entropy $\tilde{S}$ will be:

$$\tilde{S}[p] = -\sum_{\vec{s}\ \in\mathcal{A}^{\mathcal{N}}} p(\vec{s}\ )\cdot log(p(\vec{s}\ )) + \sum_{a\in\mathcal{A}}\sum_{i=1}^{\mathcal{N}} h_i^a\cdot\Big(\langle\delta_i^a(\vec{s}\ )\rangle_p - \langle\delta_i^a\rangle_{\mathcal{D}}\Big)+$$

$$+\sum_{i=1}^{\mathcal{N}-1}\sum_{a,b\in\mathcal{A}} J_{i,i+1}^{ab}\cdot\Big(\langle\delta_{i,i+1}^{ab}(\vec{s}\ )\rangle_p - \langle\delta_{i,i+1}^{ab}\rangle_{\mathcal{D}}\Big)+\sum_{i=1}^{\mathcal{N}-2}\sum_{a,b,c\in\mathcal{A}} K_{i,i+1,i+2}^{abc}\cdot\Big(\langle\delta_{i,i+1,i+1}^{abc}(\vec{s}\ )\rangle_p - \langle\delta_{i,i+1,i+2}^{abc,}\rangle_{\mathcal{D}}\Big)=$$

$$= -\sum_{\vec{s}\ \in\mathcal{A}^{\mathcal{N}}} p(\vec{s}\ )\cdot log(p(\vec{s}\ )) + \sum_{a\in\mathcal{A}}\sum_{i=1}^{\mathcal{N}} h_i^a\cdot\Big(\langle\delta_i^a(\vec{s}\ )\rangle_p - f_i^a\Big)+$$

$$+\sum_{i=1}^{\mathcal{N}-1}\sum_{a,b\in\mathcal{A}} J_{i,i+1}^{ab}\cdot\Big(\langle\delta_{i,i+1}^{ab}(\vec{s}\ )\rangle_p - f_{i,i+1}^{ab}\Big)+\sum_{i=1}^{\mathcal{N}-2}\sum_{a,b,c\in\mathcal{A}} K_{i,i+1,i+2}^{abc}\cdot\Big(\langle\delta_{i,i+1,i+1}^{abc}(\vec{s}\ )\rangle_p - f_{i,i+1,i+2}^{abc,}\Big)$$

where in the previous equation we have made use of Eq. S3. By expanding the expected values of model distribution following Eq. S2, and taking the functional derivative knowing that $\frac{\delta p(\vec{s}\ )}{\delta p(\vec{r}\ )} = \prod_{i=1}^{\mathcal{N}}\delta(s_i, r_i)$, we obtain:

$$\frac{\delta\tilde{S}[p]}{\delta p} = -\sum_{\vec{s}\ \in\mathcal{A}^{\mathcal{N}}}\frac{\delta(p(\vec{s}\ )\cdot log(p(\vec{s}\ )))}{\delta p(\vec{r}\ )} + \sum_{i=1}^{\mathcal{N}}\sum_{a\in\mathcal{A}}\sum_{\vec{s}\ \in\mathcal{A}^{\mathcal{N}}} h_i^a\cdot\delta_i^a(\vec{s}\ )\cdot\frac{\delta p(\vec{s}\ )}{\delta p(\vec{r}\ )}+$$

$$+\sum_{i=1}^{\mathcal{N}-1}\sum_{a,b\in\mathcal{A}}\sum_{\vec{s}\ \in\mathcal{A}^{\mathcal{N}}} J_{i,i+1}^{ab}\cdot\delta_{i,i+1}^{ab}(\vec{s}\ )\cdot\frac{\delta p(\vec{s}\ )}{\delta p(\vec{r}\ )}+\sum_{i=1}^{\mathcal{N}-2}\sum_{a,b,c\in\mathcal{A}}\sum_{\vec{s}\ \in\mathcal{A}^{\mathcal{N}}} K_{i,i+1,i+2}^{abc}\cdot\delta_{i,i+1,i+2}^{abc}(\vec{s}\ )\cdot\frac{\delta p(\vec{s}\ )}{\delta p(\vec{r}\ )}=$$

$$= -\sum_{\vec{s}\ \in\mathcal{A}^{\mathcal{N}}}\frac{\delta(p(\vec{s}\ )\cdot log(p(\vec{s}\ )))}{\delta p(\vec{s}\ )}\cdot\frac{\delta p(\vec{s}\ )}{\delta p(\vec{r}\ )} + \sum_{i=1}^{\mathcal{N}}\sum_{a\in\mathcal{A}}\sum_{\vec{s}\ \in\mathcal{A}^{\mathcal{N}}} h_i^a\cdot\delta_i^a(\vec{s}\ )\cdot\prod_{j=1}^{\mathcal{N}}\delta(s_j, r_j)+$$

$$+\sum_{i=1}^{\mathcal{N}-1}\sum_{a,b\in\mathcal{A}}\sum_{\vec{s}\ \in\mathcal{A}^{\mathcal{N}}} J_{i,i+1}^{ab}\cdot\delta_{i,i+1}^{ab}(\vec{s}\ )\cdot\prod_{j=1}^{\mathcal{N}}\delta(s_j, r_j)+\sum_{i=1}^{\mathcal{N}-2}\sum_{a,b,c\in\mathcal{A}}\sum_{\vec{s}\ \in\mathcal{A}^{\mathcal{N}}} K_{i,i+1,i+2}^{abc}\cdot\delta_{i,i+1,i+2}^{abc}(\vec{s}\ )\cdot\prod_{j=1}^{\mathcal{N}}\delta(s_j, r_j)=$$

$$= -\sum_{\vec{s}\ \in\mathcal{A}^{\mathcal{N}}}\frac{\delta(p(\vec{s}\ )\cdot log(p(\vec{s}\ )))}{\delta p(\vec{s}\ )}\cdot\prod_{i=1}^{\mathcal{N}}\delta(s_i, r_i) + \sum_{i=1}^{\mathcal{N}}\sum_{a\in\mathcal{A}} h_i^a\cdot\delta_i^a(\vec{r}\ )+$$

$$+\sum_{i=1}^{\mathcal{N}-1}\sum_{a,b\in\mathcal{A}} J_{i,i+1}^{ab}\cdot\delta_{i,i+1}^{ab}(\vec{r}\ )+\sum_{i=1}^{\mathcal{N}-2}\sum_{a,b,c\in\mathcal{A}} K_{i,i+1,i+2}^{abc}\cdot\delta_{i,i+1,i+2}^{abc}(\vec{r}\ )=$$

$$= -\sum_{\vec{s}\ \in\mathcal{A}^{\mathcal{N}}}\Big[log(p(\vec{s}\ ))+1\Big]\cdot\prod_{i=1}^{\mathcal{N}}\delta(s_i, r_i) + \sum_{i=1}^{\mathcal{N}}\sum_{a\in\mathcal{A}} h_i^a\cdot\delta_i^a(\vec{r}\ )+$$

$$+\sum_{i=1}^{\mathcal{N}-1}\sum_{a,b\in\mathcal{A}} J_{i,i+1}^{ab}\cdot\delta_{i,i+1}^{ab}(\vec{r}\ )+\sum_{i=1}^{\mathcal{N}-2}\sum_{a,b,c\in\mathcal{A}} K_{i,i+1,i+2}^{abc}\cdot\delta_{i,i+1,i+2}^{abc}(\vec{r}\ )=$$

$$= -log(p(\vec{r}\ )) - 1 + \sum_{i=1}^{\mathcal{N}}\sum_{a\in\mathcal{A}} h_i^a\cdot\delta_i^a(\vec{r}\ ) + \sum_{i=1}^{\mathcal{N}-1}\sum_{a,b\in\mathcal{A}} J_{i,i+1}^{ab}\cdot\delta_{i,i+1}^{ab}(\vec{r}\ )+\sum_{i=1}^{\mathcal{N}-2}\sum_{a,b,c\in\mathcal{A}} K_{i,i+1,i+2}^{abc}\cdot\delta_{i,i+1,i+2}^{abc}(\vec{r}\ )$$

Since we aim at maximizing the entropy, we impose the functional derivative to be equal to zero to look for the probability distribution which extremise $\tilde{S}$.

$$\frac{\delta\tilde{S}[p]}{\delta p} = 0 \rightarrow log(p(\vec{r}\ )) = \sum_{i=1}^{\mathcal{N}}\sum_{a\in\mathcal{A}} h_i^a\cdot\delta_i^a(\vec{r}\ ) + \sum_{i=1}^{\mathcal{N}-1}\sum_{a,b\in\mathcal{A}} J_{i,i+1}^{ab}\cdot\delta_{i,i+1}^{ab}(\vec{r}\ )+\sum_{i=1}^{\mathcal{N}-2}\sum_{a,b,c\in\mathcal{A}} K_{i,i+1,i+2}^{abc}\cdot\delta_{i,i+1,i+2}^{abc}(\vec{r}\ ) - 1$$

The probability distribution is fully attributed to a given dataset, once the complete set of forces $\vec{F} = \{h_i^a, J_{i,i+1}^{ab}, K_{i,i+1,i+2}^{abc}\}$ is known, respecting the given constraints of Eq. S4. Going back to the variable $\vec{s}$ and taking the exponential to both sides, leads to the form:

$$p(\vec{s}\ |\vec{F}) \propto e^{\sum_{i=1}^{\mathcal{N}}\sum_{a\in\mathcal{A}} h_i^a\cdot\delta_i^a(\vec{s}\ ) + \sum_{i=1}^{\mathcal{N}-1}\sum_{a,b\in\mathcal{A}} J_{i,i+1}^{ab}\cdot\delta_{i,i+1}^{ab}(\vec{s}\ )+\sum_{i=1}^{\mathcal{N}-2}\sum_{a,b,c\in\mathcal{A}} K_{i,i+1,i+2}^{abc}\cdot\delta_{i,i+1,i+2}^{abc}(\vec{s}\ )} \tag{5}$$

where, the proportionality allows to neglect the explicit constant multiplication term $e^{-1}$. By imposing the correct normalization for the probability, we can introduce the partition function $\mathcal{Z}$:

$$\mathcal{Z}(\vec{F}) = \sum_{\vec{s}\,\in\mathcal{A}^{\mathcal{N}}} e^{\sum_{i=1}^{\mathcal{N}}\sum_{a\in\mathcal{A}} h_i^a \cdot \delta_i^a(\vec{s}\,) + \sum_{i=1}^{\mathcal{N}-1}\sum_{a,b\in\mathcal{A}} J_{i,i+1}^{ab} \cdot \delta_{i,i+1}^{ab}(\vec{s}\,) + \sum_{i=1}^{\mathcal{N}-2}\sum_{a,b,c\in\mathcal{A}} K_{i,i+1,i+2}^{abc} \cdot \delta_{i,i+1,i+2}^{abc}(\vec{s}\,)} \tag{6}$$

The complete resulting probability is:

$$p(\vec{s}\,|\vec{F}) = \frac{1}{\mathcal{Z}(\vec{F})} \cdot e^{\sum_{i=1}^{\mathcal{N}}\sum_{a\in\mathcal{A}} h_i^a \cdot \delta_i^a(\vec{s}\,) + \sum_{i=1}^{\mathcal{N}-1}\sum_{a,b\in\mathcal{A}} J_{i,i+1}^{ab} \cdot \delta_{i,i+1}^{ab}(\vec{s}\,) + \sum_{i=1}^{\mathcal{N}-2}\sum_{a,b,c\in\mathcal{A}} K_{i,i+1,i+2}^{abc} \cdot \delta_{i,i+1,i+2}^{abc}(\vec{s}\,)} \tag{7}$$

which represents a Boltzmann distribution. By introducing the energy of the configuration of the system $E(\vec{s}\,|\vec{F})$, it is possible to rewrite $p$ in its more general way:

$$p(\vec{s}\,|\vec{F}) = \frac{e^{-\beta\cdot E(\vec{s}\,|\vec{F})}}{\mathcal{Z}(\vec{F})} \tag{8}$$

Imposing the equality between Eq. S7 and S8, the energy will be identical to:

$$E(\vec{s}\,|\vec{F}) = -\frac{1}{\beta}\sum_{i=1}^{\mathcal{N}}\sum_{a\in\mathcal{A}} h_i^a \cdot \delta_i^a(\vec{s}\,) - \frac{1}{\beta}\sum_{i=1}^{\mathcal{N}-1}\sum_{a,b\in\mathcal{A}} J_{i,i+1}^{ab} \cdot \delta_{i,i+1}^{ab}(\vec{s}\,) - \frac{1}{\beta}\sum_{i=1}^{\mathcal{N}-2}\sum_{a,b,c\in\mathcal{A}} K_{i,i+1,i+2}^{abc} \cdot \delta_{i,i+1,i+2}^{abc}(\vec{s}\,) \tag{9}$$

### A2: Inference

To proceed in finding the set of forces $\vec{F}$ which satisfy the constraints imposed through the Lagrange multipliers at Eq. S4, we will follow a Bayesian inference method. We hence define the following loss function $\mathcal{L}$:

$$\mathcal{L}(\vec{F}|\{\vec{\mathcal{D}}_i\}_{i=1}^M) = -log\Big(\prod_{i=1}^{M} p^{w_i}(\vec{\mathcal{D}}_i|\vec{F})\Big) = -log\Big(\prod_{i=1}^{M}\frac{1}{\mathcal{Z}^{w_i}(\vec{F})}\cdot e^{-\beta\,\cdot\, w_i\,\cdot\, E(\vec{\mathcal{D}}_i|\vec{F})}\Big) =$$

$$= \sum_{i=1}^{M} w_i \cdot log(\mathcal{Z}(\vec{F})) + \beta\cdot\sum_{i=1}^{M} w_i \cdot E(\vec{\mathcal{D}}_i|\vec{F})$$

where $\vec{\mathcal{D}}_i$ correspond to the sequence of the dataset $\mathcal{D}$ at row $i$ and $\{\vec{\mathcal{D}}_i\}_{i=1}^M$ represents the collection of all $M$ sequences. When all the weights, which corresponds to the *stoichiometry*, are equivalent to one (for all non-methylated adenosines or due to inaccessibility in methylated sites due to experimental limitations) the previous quantity exactly corresponds (up to a sign) to the log-likelihood function. We now take the partial derivative relatively to a force parameter. Without any loss of generality we show it for the generic mono-mer force $h_j^b$:

$$\frac{\partial\mathcal{L}(\vec{F}|\{\vec{\mathcal{D}}_i\}_{i=1}^M)}{\delta h_j^b} = \sum_{i=1}^{M} w_i\cdot\frac{\partial log(\mathcal{Z}(\vec{F}))}{\partial h_j^b} + \beta\cdot\sum_{i=1}^{M} w_i\cdot\frac{\partial E(\vec{\mathcal{D}}_i|\vec{F})}{\partial h_j^b} \tag{10}$$

We focus on the first term of the previous equation:

$$\frac{\partial log(\mathcal{Z}(\vec{F}))}{\partial h_j^b} = \frac{1}{\mathcal{Z}(\vec{F})}\frac{\partial\mathcal{Z}(\vec{F})}{\partial h_j^b} =$$

$$= \frac{1}{\mathcal{Z}(\vec{F})}\frac{\partial}{\partial h_j^b}\Bigg(\sum_{\vec{s}\,\in\mathcal{A}^{\mathcal{N}}} e^{\sum_{i=1}^{\mathcal{N}}\sum_{a\in\mathcal{A}} h_i^a \cdot \delta_i^a(\vec{s}\,) + \sum_{i=1}^{\mathcal{N}-1}\sum_{a,b\in\mathcal{A}} J_{i,i+1}^{ab} \cdot \delta_{i,i+1}^{ab}(\vec{s}\,) + \sum_{i=1}^{\mathcal{N}-2}\sum_{a,b,c\in\mathcal{A}} K_{i,i+1,i+2}^{abc} \cdot \delta_{i,i+1,i+2}^{abc}(\vec{s}\,)}\Bigg) =$$

$$= \frac{1}{\mathcal{Z}(\vec{F})}\sum_{\vec{s}\,\in\mathcal{A}^{\mathcal{N}}}\delta_j^b(\vec{s}\,)\cdot e^{\sum_{i=1}^{N}\sum_{a\in\mathcal{A}} h_i^a \cdot \delta_i^a(\vec{s}\,) + \sum_{i=1}^{N-1}\sum_{a,b\in\mathcal{A}} J_{i,i+1}^{ab} \cdot \delta_{i,i+1}^{ab}(\vec{s}\,) + \sum_{i=1}^{N-2}\sum_{a,b,c\in\mathcal{A}} K_{i,i+1,i+2}^{abc} \cdot \delta_{i,i+1,i+2}^{abc}(\vec{s}\,)} = \langle\delta_j^b(\vec{s}\,)\rangle_p$$

where we have exploited the relation: $\frac{\partial h_i^a}{\partial h_j^b} = \delta(i,j) \cdot \delta(a,b)$. The same can be obtained for each force parameter, following the same computation, leading to the generic equality:

$$\frac{\partial log(\mathcal{Z}(\vec{F}))}{\partial F^{\vec{S}}_{i,i+L-1}} = \langle \delta^{\vec{S}}_{i,i+L-1}(\vec{s}\,) \rangle_p \tag{11}$$

For the second term:

$$\frac{\partial E(\vec{\mathcal{D}}_i|\vec{F})}{\partial h_j^b} = -\frac{1}{\beta}\frac{\partial}{\partial h_j^b}\left( \sum_{i=1}^{\mathcal{N}} \sum_{a\in\mathcal{A}} h_i^a \cdot \delta_i^a(\vec{\mathcal{D}}_i) + \sum_{i=1}^{\mathcal{N}-1} \sum_{a,b\in\mathcal{A}} J^{ab}_{i,i+1} \cdot \delta^{ab}_{i,i+1}(\vec{\mathcal{D}}_i) + \sum_{i=1}^{\mathcal{N}-2} \sum_{a,b,c\in\mathcal{A}} K^{abc}_{i,i+1,i+2} \cdot \delta^{abc}_{i,i+1,i+2}(\vec{\mathcal{D}}_i) \right) =$$

$$= -\frac{1}{\beta} \cdot \delta_j^b(\vec{\mathcal{D}}_i) = -\frac{1}{\beta} \cdot \delta(\mathcal{D}_{i,j}, b)$$

Substituting the two results back into Eq. S10:

$$\frac{\partial \mathcal{L}(\vec{F}|\{\vec{\mathcal{D}}_i\}_{i=1}^M)}{\delta h_j^b} = \sum_{i=1}^M w_i \cdot \langle \delta_j^b(\vec{s}\,) \rangle_p - \not\beta \cdot \sum_{i=1}^M w_i \cdot \frac{\delta(\mathcal{D}_{i,j}, b)}{\not\beta}$$

By multiplying and dividing the second term of the equation by $\sum_{i=1}^M w_i$:

$$\frac{\partial \mathcal{L}(\vec{F}|\{\vec{\mathcal{D}}_i\}_{i=1}^M)}{\delta h_j^b} = \sum_{i=1}^M w_i \cdot \langle \delta_j^b(\vec{s}\,) \rangle_p - \sum_{i=1}^M w_i \cdot \frac{\sum_{i=1}^M w_i \cdot \delta(\mathcal{D}_{i,j}, b)}{\sum_{i=1}^M w_i} = \sum_{i=1}^M w_i \cdot \Big( \langle \delta_j^b(\vec{s}\,) \rangle_p - \langle \delta_j^b \rangle_{\mathcal{D}} \Big) = \sum_{i=1}^M w_i \cdot \Big( \langle \delta_j^b(\vec{s}\,) \rangle_p - f_j^b \Big) = 0$$

Where in the last equality, we have made use of the constraints imposed in Eq. S4. Therefore, by generalizing to all the force parameters, extremizing the loss function correspond to finding the values of the forces which satisfy the wanted constraints:

$$\frac{\partial \mathcal{L}(\vec{F}|\{\vec{\mathcal{D}}_i\}_{i=1}^M)}{\delta F^{\vec{S}}_{i,i+1-L}} = \langle \delta^{\vec{S}}_{i,i+L-1}(\vec{s}\,) \rangle_p - f^{\vec{S}}_{i,i+L-1} = 0 \tag{12}$$

One last proof, for the previous statement, comes from the computation of the second derivative to understand the curvature of the function and to claim that the extreme point represent indeed a minimum:

$$\frac{\partial^2 \mathcal{L}(\vec{F}|\{\vec{\mathcal{D}}_i\}_{i=1}^M)}{\partial^2 h_j^b} = \frac{\partial}{\partial h_j^b}\left( \sum_{i=1}^M w_i \cdot (\langle \delta_j^b((\vec{s}\,)) \rangle_p - f_j^b) \right) = \sum_{i=1}^M w_i \frac{\partial}{\partial h_j^b} \langle \delta_j^b(\vec{s}\,) \rangle_p = \sum_{i=1}^M w_i \cdot \frac{\partial}{\partial h_j^b} \sum_{\vec{s}\in\mathcal{A}^{\mathcal{N}}} \delta_j^b(\vec{s}\,) \cdot p(\vec{s}\,) =$$

$$= \sum_{i=1}^M w_i \cdot \frac{\partial}{\partial h_j^b}\left( \sum_{\vec{s}\in\mathcal{A}^{\mathcal{N}}} \delta_j^b(\vec{s}\,) \cdot \frac{1}{\mathcal{Z}(\vec{s}\,|\vec{F})} \cdot e^{-\beta E(\vec{F})} \right) = \sum_{i=1}^M w_i \cdot \sum_{\vec{s}\in\mathcal{A}^{\mathcal{N}}} \delta_j^b(\vec{s}\,) \cdot \left( -\frac{1}{\mathcal{Z}^2(\vec{F})} \frac{\partial \mathcal{Z}(\vec{F})}{\partial h_j^b} \cdot e^{-\beta \cdot E(\vec{s}\,|\vec{F})} + \frac{1}{\mathcal{Z}(\vec{F})} \frac{\partial e^{-\beta \cdot E(\vec{s}\,|\vec{F})}}{\partial h_j^b} \right) =$$

$$= \sum_{i=1}^M w_i \cdot (-\langle \delta_j^b(\vec{s}\,) \rangle_p^2 + \langle (\delta_j^b(\vec{s}\,))^2 \rangle_p) \geq 0$$

where for the inequality we have exploited the known property of the variance of a random variable:

$$Var(\delta_j^b(\vec{s}\,)) = (\langle (\delta_j^b(\vec{s}\,))^2 \rangle_p - \langle \delta_j^b(\vec{s}\,) \rangle_p^2) \geq 0$$

and the characteristics of the weights, which are always positive, leading to $\sum_{i=1}^M w_i > 0$. The loss function is therefore always non-negative for any given combination of the force parameters. $\mathcal{L}$ being a convex function, admits a global minimum, which corresponds to the one in which the forces satisfy the given constraints from Eq. S4. It is therefore sufficient to apply a numerical minimization algorithm to infer the set of parameters $\vec{F}$.

For this end, we will apply the minimization procedure, from Eq. S10 to:

$$\mathcal{L}'(\vec{F}|\{\vec{\mathcal{D}}_i\}_{i=1}^M) = log(\mathcal{Z}(\vec{F})) - \sum_{i=1}^{\mathcal{N}+1-L} \sum_{\vec{S}\in\mathcal{A}^L} F^{\vec{S}}_{i,i+L-1} \cdot \langle \delta^{\vec{S}}_{i,i+L-1} \rangle_{\mathcal{D}}$$

which is an equivalent quantity to $\mathcal{L}$ (up to the positive constant $\sum_{i=1}^M w_i$) that provides exactly the same minimum as described in Eq. S12. Moreover, to avoid overfitting, we will add to $\mathcal{L}'$ an hyperparameter $\lambda$ to impose a L2-regularization term. The final form of the loss function will correspond to:

$$\tilde{\mathcal{L}}(\vec{F}|\{\vec{\mathcal{D}}_i\}_{i=1}^M) = log(\mathcal{Z}(\vec{F})) - \sum_{i=1}^{\mathcal{N}+1-L} \sum_{\vec{S}\in\mathcal{A}^L} F^{\vec{S}}_{i,i+L-1} \cdot \langle \delta^{\vec{S}}_{i,i+L-1} \rangle_{\mathcal{D}} + \lambda \cdot \sum_{i=1}^{\mathcal{N}+1-L} \sum_{\vec{S}\in\mathcal{A}^L} (F^{\vec{S}}_{i,i+L-1})^2$$

By defining $\eta$ the learning rate, $\xi$ the maximum number of iteration, $\epsilon$ the tolerance and $\gamma$ the momentum term, a gradient descent with momentum can be implemented, iterating, at each time step, over the set of equations:

$$\begin{cases} \vec{v}_t = \gamma \cdot \vec{v}_{t-1} + \eta \cdot \nabla_F \tilde{\mathcal{L}}(\vec{F}_t) \\ \vec{F}_{t+1} = \vec{F}_t - \vec{v}_t \end{cases}$$

where $\vec{v}_{t=0}$ and $\vec{F}_{t=1}$ are initialized at the first step. The recursion ends, either when the maximum element (in absolute value) of the gradient is lower than the tolerance value, $max(|\nabla_F \tilde{\mathcal{L}}(\vec{F}_t)|) < \epsilon$ or the recursion exceed the maximum number of iterations, $t > \xi$. In the procedure the maximum number of iterations has been set to $\xi = 10\,000$, while $\eta = 2$ and $\gamma = 0.8$. The hyperparameters $\epsilon$ and $\lambda$ have been optimized over a given set of data (see main Validation paragraph).

### A3: Transfer Matrix computation

From Eq. S7, it is possible to rewrite the probability distribution, through the full Kronecker delta definition as:

$$p(\vec{s}\,|\vec{F}) = \frac{1}{\mathcal{Z}} e^{\sum_{i=1}^{\mathcal{N}} \sum_{a \in \mathcal{A}} h_i^a \cdot \delta(s_i, a) + \sum_{i=1}^{\mathcal{N}-1} \sum_{a,b \in \mathcal{A}} J_{i,i+1}^{ab} \cdot \delta(s_i, a) \cdot \delta(s_{i+1}, b)} \cdot$$
$$\cdot e^{\sum_{i=1}^{\mathcal{N}-2} \sum_{a,b,c \in \mathcal{A}} K_{i,i+1,i+2}^{abc} \cdot \delta(s_i, a) \cdot \delta(s_{i+1}, b) \cdot \delta(s_{i+2}, c)}$$

with the related partition function:

$$\mathcal{Z}(\vec{F}) = \sum_{s_1 \in \mathcal{A}} \sum_{s_2 \in \mathcal{A}} \dots \sum_{s_{\mathcal{N}} \in \mathcal{A}} e^{\sum_{i=1}^{\mathcal{N}} \sum_{a \in \mathcal{A}} h_i^a \cdot \delta(s_i, a)} \cdot$$
$$\cdot e^{\sum_{i=1}^{\mathcal{N}-1} \sum_{a,b \in \mathcal{A}} J_{i,i+1}^{ab} \cdot \delta(s_i, a) \cdot \delta(s_{i+1}, b)} \cdot$$
$$\cdot e^{\sum_{i=1}^{\mathcal{N}-2} \sum_{a,b,c \in \mathcal{A}} K_{i,i+1,i+2}^{abc} \cdot \delta(s_i, a) \cdot \delta(s_{i+1}, b) \cdot \delta(s_{i+2}, c)}$$

For the analytical computation of the partition function we will exploit the transfer matrix method. At this point, it is not possible to rewrite the partition function in terms of the transfer matrix $\hat{T}_{s_i, s_{i+1}}$, since the statistics of tri-nucleotides cannot be taken into account when focusing into two consecutive sites. The computation needs to be extended to include larger window statistics. To this aim we consider a set of new variables, made of two nucleotides, in such a way that the transfer matrix can be written in the new form as: $\hat{T}_{(s_i, s_{i+1}),(s_{i+1}, s_{i+2})}$. By defining the new coarse-grained nucleotides $\vec{S}_i^* = (s_i, s_{i+1})$, the transfer matrix can be denoted as $\hat{T}_{\vec{S}_i^*, \vec{S}_{i+1}^*}$. In general we can therefore map the problem into a new one, where the sequence $\vec{s}$ is generalized to:

$$\vec{s}^{\,*} = (S_{i,1}^*, S_{i,2}^*, S_{i+1,1}^*, \dots, S_{\mathcal{N}-1,1}^*, S_{\mathcal{N}-1,2})$$

with the additional constraints:

$$S_{i,2}^* = S_{i+1,1}^* = s_{i+1} \quad \forall i = 1, \dots, \mathcal{N} - 2$$

Given the previous equalities, it is possible to rewrite the partition function as follows:

$$\mathcal{Z}(\vec{F}) = \sum_{\vec{S}_1^* \in \mathcal{A}^2} \sum_{\vec{S}_2^* \in \mathcal{A}^2} \dots \sum_{\vec{S}_{\mathcal{N}-1}^* \in \mathcal{A}^2} \prod_{i=1}^{\mathcal{N}-2} \delta(S_{i,2}^*, S_{i+1,1}^*) \cdot$$
$$\cdot e^{\sum_{i=1}^{\mathcal{N}-2} \sum_{a \in \mathcal{A}} \left[ \frac{h_i^a}{3} \cdot \delta(S_{i,1}^*, a) + \frac{h_{i+1}^a}{3} \cdot \delta(S_{i,2}^*, a) + \frac{h_{i+2}^a}{3} \cdot \delta(S_{i+1,1}^*, a) \right]} \cdot$$
$$\cdot e^{\sum_{i=1}^{\mathcal{N}-1} \sum_{a,b \in \mathcal{A}} \left[ \frac{J_{i,i+1}^{ab}}{2} \cdot \delta(S_{i,1}^*, a) \cdot \delta(S_{i,2}^*, b) + \frac{J_{i+1,i+2}^{ab}}{2} \cdot \delta(S_{i+1,1}^*, a) \cdot \delta(S_{i+1,2}^*, b) \right]} \cdot$$

$$\cdot e^{\sum_{i=1}^{\mathcal{N}-2}\sum_{a,b,c\in\mathcal{A}} K^{abc}_{i,i+1,i+2}\cdot\delta(S^*_{i,1},a)\cdot\delta(S^*_{i,2},b)\cdot\delta(S^*_{i+1,1},c)}\cdot$$

$$\cdot e^{\sum_{a\in\mathcal{A}}\left[\frac{2}{3}\cdot\left(h^a_1\cdot\delta(S^*_{1,1},a)+h^a_{\mathcal{N}}\cdot\delta(S^*_{\mathcal{N}-1,2},a)\right)+\frac{1}{3}\cdot\left(h^a_2\cdot\delta(S^*_{1,2}-a)+h^a_{\mathcal{N}-1}\cdot\delta(S^*_{\mathcal{N}-1,1})\right)\right]}\cdot$$

$$\cdot e^{\sum_{a,b\in\mathcal{A}}\frac{1}{2}\left(J^{ab}_{1,2}\cdot\delta(S^*_{1,1},a)\cdot\delta(S^*_{1,2},b)+J^{ab}_{\mathcal{N}-1,\mathcal{N}}\cdot\delta(S^*_{\mathcal{N}-1,1},b)\cdot\delta(S^*_{\mathcal{N}-1,2},b)\right)}$$

By applying definition of the Kronecker deltas and expliciting the dependence of all sum in terms of $\mathcal{N}-2$ terms:

$$\mathcal{Z}(\vec{F})=\sum_{\vec{S}^*_1\in\mathcal{A}^2}\sum_{\vec{S}^*_2\in\mathcal{A}^2}\cdots\sum_{\vec{S}^*_{\mathcal{N}-1}\in\mathcal{A}^2}\prod_{i=1}^{\mathcal{N}-2}\delta(S^*_{i,2},S^*_{i+1,1})\cdot$$

$$\cdot e^{\sum_{i=1}^{\mathcal{N}-2}\left[\frac{h_i^{S^*_{i,1}}}{3}+\frac{h_{i+1}^{S^*_{i,2}}}{3}+\frac{h_{i+2}^{S^*_{i+1,1}}}{3}\right]+\sum_{i=1}^{\mathcal{N}-2}\left[\frac{J_{i,i+1}^{S^*_{i,1},S^*_{i,2}}}{2}+\frac{J_{i+1,i+2}^{S^*_{i+1,1}S^*_{i+1,2}}}{2}\right]+\sum_{i=1}^{\mathcal{N}-2}K_{i,i+1,i+2}^{S^*_{i,1}S^*_{i,2}S^*_{i+1,1}}}\cdot$$

$$\cdot e^{\frac{2}{3}\cdot\left(h_1^{S^*_{1,1}}+h_{\mathcal{N}}^{S^*_{\mathcal{N}-1,2}}\right)+\frac{1}{3}\cdot\left(h_2^{S^*_{1,2}}+h_{\mathcal{N}-1}^{S^*_{\mathcal{N}-1,1}}\right)+\frac{1}{2}\cdot\left(J_{1,2}^{S^*_{1,1}S^*_{1,2}}+J_{\mathcal{N}-1,\mathcal{N}}^{S^*_{\mathcal{N}-1,1}S^*_{\mathcal{N}-1,2}}\right)}$$

Therefore we can group all terms which depend on $\mathcal{N}-2$ separating them from the remaining ones:

$$\mathcal{Z}(\vec{F})=\sum_{\vec{S}^*_1\in\mathcal{A}^2}\sum_{\vec{S}^*_2\in\mathcal{A}^2}\cdots\sum_{\vec{S}^*_{\mathcal{N}-1}\in\mathcal{A}^2}\prod_{i=1}^{\mathcal{N}-2}\Bigg[\delta(S^*_{i,2},S^*_{i+1,1})\cdot$$

$$\cdot e^{\left[\frac{h_i^{S^*_{i,1}}}{3}+\frac{h_{i+1}^{S^*_{i,2}}}{3}+\frac{h_{i+2}^{S^*_{i+1,1}}}{3}\right]+\left[\frac{J_{i,i+1}^{S^*_{i,1},S^*_{i,2}}}{2}+\frac{J_{i+1,i+2}^{S^*_{i+1,1}S^*_{i+1,2}}}{2}\right]+K_{i,i+1,i+2}^{S^*_{i,1}S^*_{i,2}S^*_{i+1,1}}}\cdot$$

$$\cdot e^{\frac{2}{3}\cdot\left(h_1^{S^*_{1,1}}+h_{\mathcal{N}}^{S^*_{\mathcal{N}-1,2}}\right)+\frac{1}{3}\cdot\left(h_2^{S^*_{1,2}}+h_{\mathcal{N}-1}^{S^*_{\mathcal{N}-1,1}}\right)+\frac{1}{2}\cdot\left(J_{1,2}^{S^*_{1,1}S^*_{1,2}}+J_{\mathcal{N}-1,\mathcal{N}}^{S^*_{\mathcal{N}-1,1}S^*_{\mathcal{N}-1,2}}\right)}=$$

$$=\sum_{\vec{S}^*_1\in\mathcal{A}^2}\sum_{\vec{S}^*_2\in\mathcal{A}^2}\cdots\sum_{\vec{S}^*_{\mathcal{N}-1}\in\mathcal{A}^2}\prod_{i=1}^{\mathcal{N}-2}\hat{T}_{\vec{S}^*_i,\vec{S}^*_{i+1}}\cdot\hat{T}_{\vec{S}^*_{\mathcal{N}-1},\vec{S}^*_1}\tag{13}$$

Where in the previous steps we have defined:

$$\hat{T}_{\vec{S}^*_i,\vec{S}^*_{i+1}}=\delta(S^*_{i,2},S^*_{i+1,1})\cdot e^{\frac{h_i^{S^*_{i,1}}}{3}+\frac{h_{i+1}^{S^*_{i,2}}}{3}+\frac{h_{i+2}^{S^*_{i+1,1}}}{3}+\frac{J_{i,i+1}^{S^*_{i,1},S^*_{i,2}}}{2}+\frac{J_{i+1,i+2}^{S^*_{i+1,1}S^*_{i+1,2}}}{2}+K_{i,i+1,i+2}^{S^*_{i,1}S^*_{i,2}S^*_{i+1,1}}}$$

$$\hat{T}_{S^*_{\mathcal{N}-1},S^*_1}=e^{\frac{2}{3}\cdot\left(h_1^{S^*_{1,1}}+h_{\mathcal{N}}^{S^*_{\mathcal{N}-1,2}}\right)+\frac{1}{3}\cdot\left(h_2^{S^*_{1,2}}+h_{\mathcal{N}-1}^{S^*_{\mathcal{N}-1,1}}\right)+\frac{1}{2}\cdot\left(J_{1,2}^{S^*_{1,1}S^*_{1,2}}+J_{\mathcal{N}-1,\mathcal{N}}^{S^*_{\mathcal{N}-1,1}S^*_{\mathcal{N}-1,2}}\right)}$$

The analytical form of $\mathcal{Z}$ is fully known, and can be efficiently computed numerically thanks to Eq. S13. It is possible to infer all the parameters $\vec{F}$ by minimizing the loss function, as described in Appendix A2.

### A4: Gauge Choice

The number of all parameters $\vec{F}$ to infer, for motifs of general length $\mathcal{N}$ is: $4\cdot\mathcal{N}$ for all single forces $h^a_i$, $16\cdot(\mathcal{N}-1)$ for all coupling $J^{ab}_{i,i+1}$ and $64\cdot(\mathcal{N}-2)$ for all three sites forces $K^{abc}_{i,i+1,i+2}$. So the total number of parameters is: $84\cdot\mathcal{N}-144=528$ parameters for 8-mer motifs. Amongst all these parameters, only a much smaller subset consists in actual independent ones, due to the frequency constraints of 1,2,3-mers. Let's now consider the computation to count the amount of all dependent parameter. At first we impose that the frequency of all 1,2,3-mers at a given position must sum up to one:

$$\sum_{a\in\mathcal{A}} f^a_i=1\ \forall i=1,...,\mathcal{N}\rightarrow\#\mathcal{N}$$

$$\sum_{ab\in\mathcal{A}^2} f^{ab}_{i,i+1} = 1 \ \forall i = 1, ..., \mathcal{N}-1 \rightarrow \#\mathcal{N}-1$$

$$\sum_{abc\in\mathcal{A}^3} f^{abc}_{i,i+1,i+2} = 1 \ \forall i = 1, ..., \mathcal{N}-2 \rightarrow \#\mathcal{N}-2$$

This intermediate step accounts for $3 \cdot \mathcal{N} - 3$ free forces. Let's consider now, that at a given position 1-mers are linked to 2-mers. As a matter of fact:

$$\sum_{a\in\mathcal{A}} f^{ax}_{i,i+1} = f^{x}_{i+1} \ \forall i = 1, ..., \mathcal{N}-1 \ \wedge \ \forall x \in \mathcal{A} \rightarrow \#4 \cdot (\mathcal{N}-1)$$

$$\sum_{a\in\mathcal{A}} f^{xa}_{i,i+1} = f^{x}_{i} \ \forall i = 1, ..., \mathcal{N}-1 \ \wedge \ \forall x \in \mathcal{A} \rightarrow \#4 \cdot (\mathcal{N}-1)$$

Giving a total of additional dependent parameters of: $8 \cdot \mathcal{N} - 8$. In the same way 3-mers are linked to 1-mers:

$$\sum_{ab\in\mathcal{A}} f^{abx}_{i,i+1,i+2} = f^{x}_{i+2} \ \forall i = 1, ..., \mathcal{N}-2 \ \wedge \ x \in \mathcal{A} \rightarrow \#4 \cdot (\mathcal{N}-2)$$

$$\sum_{ab\in\mathcal{A}} f^{axb}_{i,i+1,i+2} = f^{x}_{i+1} \ \forall i = 1, ..., \mathcal{N}-2 \ \wedge \ x \in \mathcal{A} \rightarrow \#4 \cdot (\mathcal{N}-2)$$

$$\sum_{ab\in\mathcal{A}} f^{xab}_{i,i+1,i+2} = f^{x}_{i} \ \forall i = 1, ..., \mathcal{N}-2 \ \wedge \ x \in \mathcal{A} \rightarrow \#4 \cdot (\mathcal{N}-2)$$

with additional $12 \cdot \mathcal{N} - 24$ free parameters. Finally even 3-mers are linked to 2-mers by the following constraints:

$$\sum_{a\in\mathcal{A}} f^{xya}_{i,i+1,i+2} = f^{xy}_{i,i+1} \ \forall i = 1, ..., \mathcal{N}-2 \ \wedge \ xy \in \mathcal{A}^2 \rightarrow \#16 \cdot (\mathcal{N}-2)$$

$$\sum_{a\in\mathcal{A}} f^{axy}_{i,i+1,i+2} = f^{xy}_{i+1,i+2} \ \forall i = 1, ..., \mathcal{N}-2 \ \wedge \ xy \in \mathcal{A}^2 \rightarrow \#16 \cdot (\mathcal{N}-2)$$

accounting of $32 \cdot \mathcal{N} - 64$ dependent forces. By considering all free parameters together:

$$(3 \cdot \mathcal{N} - 3) + (8 \cdot \mathcal{N} - 8) + (12 \cdot \mathcal{N} - 24) + (32 \cdot \mathcal{N} - 64) = 55 \cdot \mathcal{N} - 99$$

which corresponds in the case of 8-mers to 341 free parameters. By putting together the whole number of parameters and the amount of dependent ones, it is possible to obtain the final number of independent forces:

$$(84 \cdot \mathcal{N} - 144) - (55 \cdot \mathcal{N} - 99) = 29 \cdot \mathcal{N} - 45$$

which for the 8-mers correspond to 187 independent parameters out the total 528. For the inference process, it has been chosen the zero-gauge choice, in which all dependent parameter are set to zero values. Any different gauge choice, will always lead to the same probability and with the same properties shown at Eq. S4.

### A5: Bayesian posterior

Since we will always define a distribution for methylated and non-methylated sites, we will indicate the forces that allow to parametrize the probability distribution from Eq. S7 with methylated label $m = 1$ and non-methylated $m = 0$: $p(\vec{s}\,|1)$ and $p(\vec{s}\,|0)$. In this way, by defining $m = \{0, 1\}$ the two possible labels, we will indicate the previously mentioned equation, developing the Kronecker deltas as in the transfer matrix computation, as follows:

$$p(\vec{s}\,|m) \propto e^{\sum_{i=1}^{\mathcal{N}} h^{s_i,m}_i + \sum_{i=1}^{\mathcal{N}-1} J^{s_i s_{i+1},m}_{i,i+1} + \sum_{i=1}^{\mathcal{N}-2} K^{s_i,s_{i+1},s_{i+2},m}_{i,i+1,i+2}} =$$
$$= e^{\sum_{i=1}^{\mathcal{N}} h^{m}_i(s_i) + \sum_{i=1}^{\mathcal{N}-1} J^{m}_{i,i+1}(s_i, s_{i+1},) + \sum_{i=1}^{\mathcal{N}-2} K^{m}_{i,i+1,i+2}(s_i, s_{i+1}, s_{i+2},)} \qquad (14)$$

where in the last equality, the notation has changed, for simplicity, to include the parametrization of the dataset from which the inference has been performed. In this way, it is possible to retrieve back the equations as presented in the main article

(Eq. 3), where $\mathcal{Z}(\vec{F}) \to \mathcal{Z}^m$ and $E(\vec{s}\,|\vec{F}) \to E(\vec{s}\,|m)$. Since we are interested in predicting the probability of a given motif to be methylated, we can then apply Bayes theorem:

$$p(m|\vec{s}\,) \propto p(\vec{s}\,|m) \cdot p_0(m)$$

where $p_0(m = 1), p_0(m = 0)$ are *prior* knowledge relatively to the methylation imbalance, which quantify, independently of the motif, the ratio of methylated or non-methylated sites. The two probabilities, are strictly linked by the following simple relation:

$$\sum_{m\in\{0,1\}} p(m|\vec{s}\,) = p(1|\vec{s}\,) + p(0|\vec{s}\,) = 1$$

Therefore, introducing the common normalization factor $k(\vec{s}\,)$, since it must be independent of the methylation label, we can obtain:

$$k(\vec{s}\,)\Big[p(\vec{s}\,|1) \cdot p_0(1) + p(\vec{s}\,|0) \cdot p_0(0)\Big] = 1 \to k(\vec{s}\,) = \frac{1}{\sum_m p(\vec{s}\,|m) \cdot p_0(m)} \tag{15}$$

Therefore, fixing $k$ for the motif $\vec{s}$, we can finally obtained the methylation probability (as in main Eq. 4):

$$p(1|\vec{s}\,) = \frac{p(\vec{s}\,|1) \cdot p_0(1)}{\sum\limits_{m=\{0,1\}} p(\vec{s}\,|m) \cdot p_0(m)} \tag{16}$$

### A6: $m^6A$ Kinetics

#### Two-State Model

Let's consider a simple dynamical process based only on $m^6A$ writers and erasers, leading to methylated $m = 1$ or non-methylated $m = 0$ states. From a chemical point of view this can be represented as the following process:

$$0 \underset{k_{off}}{\overset{k_{on}}{\rightleftharpoons}} 1$$

where $k_{on}$ and $k_{off}$ correspond to the reaction rates for the writer complex to perform the methylation and for the eraser to de-methylate a given site. It is possible to write the following set of differential equations:

$$\begin{cases} \frac{d\rho(1|t)}{dt} = -k_{off} \cdot \rho(1|t) + k_{on} \cdot \rho(0|t) \\ \frac{d\rho(0|t)}{dt} = +k_{off} \cdot \rho(1|t) - k_{on} \cdot \rho(0|t) \end{cases}$$

where $\rho(1|t)$ and $\rho(0|t)$ represent the probability to be in a given state at time $t$. The two equations are additionally linked by the following constraint:

$$\rho(1|t) + \rho(0|t) = 1 \qquad \forall t$$

From this set of equations it is possible to find the general solution:

$$\rho(1|t) = \frac{k_{on}}{k_{on} + k_{off}} \cdot \left(1 - e^{-t \cdot (k_{on} + k_{off})}\right)$$

In the following we will assume that the methylation reaction $k_{on}$ and de-methylation $k_{off}$ are considered only parametrically dependent on the motif surrounding the adenosine:

$$\begin{cases} k_{on} = k_{on}(\vec{s}\,) \\ k_{off} = k_{off}(\vec{s}\,) \end{cases}$$

by writing in an explicit way the probability parameters, the final expression results:

$$\rho(1|t, k_{on}, k_{off}, \vec{s}\,) = \frac{k_{on}(\vec{s}\,)}{k_{on}(\vec{s}\,) + k_{off}(\vec{s}\,)} \cdot \left(1 - e^{-t \cdot (k_{on}(\vec{s}\,) + k_{off}(\vec{s}\,))}\right) \tag{17}$$

By considering the equilibrium probability, it is possible to find a relationship linking the set of reaction rates:

$$\lim_{t\to\infty} \rho(1|t, k_{off}, \vec{s}\,) = \frac{k_{on}(\vec{s}\,)}{k_{on}(\vec{s}\,) + k_{off}(\vec{s}\,)}$$

Moreover, if now we assume that the detected $m^6A$ sites in a given experiemental dataset are obtained after the equilibrium condition is reached, it is possible to connect the two-state model with MEMAR-C:

$$\lim_{t\to\infty} \rho(1|t, k_{off}, \vec{s}\,) = \frac{k_{on}(\vec{s}\,)}{k_{on}(\vec{s}\,) + k_{off}(\vec{s}\,)} = p(1|\vec{s}\,) \tag{18}$$

#### AUPRC from $m^6A$ dynamical process

We consider a generic dataset $\mathcal{D} = \mathcal{D}_{methylated} \cup \mathcal{D}_{non-methylated}$ and define the vector $\vec{N}^{\mathcal{D}} = (N_{\vec{s}_1}, ...N_{\vec{s}_{4^{\mathcal{N}}}})$ where $N_{\vec{s}_i}$ represent the total number of times a given motif, amongst all the possible combination of $\mathcal{N}$ letters, is present in the dataset. Based on the two-state dynamical process defined in Eq. S17, it is possible to quantify the amount of methylated sites in a given dataset for any given time $t$. This can be used to later assess the $m^6A$ predictor performance, computing the AUPRC as a function of time. To this end, we consider a random variable $x$ which can take as values either 0 or 1 based on the methylation state:

$$x = \begin{cases} 1 & methylated \\ 0 & non-methylated \end{cases} \tag{19}$$

Then let's consider the set $\vec{x}_1, ..., \vec{x}_{4^{\mathcal{N}}}$ as the total number of random variables, associated to each motif $\vec{s}_i$ in the given dataset, such that: $\vec{x_1} = (x_1^1, ..., x_1^{N_{\vec{s}_1}}), ..., \vec{x}_{4^{\mathcal{N}}} = (x_{4^{\mathcal{N}}}^1, ..., x_{4^{\mathcal{N}}}^{N_{\vec{s}_{4^{\mathcal{N}}}}})$. Each of this random variable is distributed as a Bernoulli:

$$\mathcal{P}(x_i^j|\rho) = \begin{cases} \rho(1|t, k_{on}, k_{off}, \vec{s_j}) & x_i^j = 1 \\ 1 - \rho(1|t, k_{on}, k_{off}, \vec{s_j}) & x_i^j = 0 \end{cases} \tag{20}$$

We can for instance compute the probability that $c$ motifs $\vec{s}_j$ are methylated:

$$\mathcal{P}\left(\sum_{i=1}^{N_j} x_j^i = c|\rho\right) = \binom{N_j}{c} \cdot \rho^c(1|t, k_{on}, k_{off}, \vec{s_j}) \cdot \rho^{N_j - c}(0|t, k_{on}, k_{off}, \vec{s_j})$$

which is equivalent to a Binomial distribution since the set $\vec{x}_j$ is identically and independently distributed. The resulting average number of methylated $\vec{s}_j$ motifs results in:

$$\left\langle \sum_{i=1}^{N_j} x_j^i \right\rangle = N_{\vec{s_j}} \cdot \rho(1|t, k_{on}, k_{off}, \vec{s_j}) \tag{21}$$

If we want now to compute the total average number of methylated sites in the whole dataset, this corresponds to:

$$N_{m^6A}^{\mathcal{D}}(\rho) = \left\langle \sum_{j=1}^{4^{\mathcal{N}}} \sum_{i=1}^{N_{\vec{s}_j}} x_j^i \right\rangle = \left\langle \sum_{i=1}^{N_{\vec{s}_1}} x_1^i + \sum_{i=1}^{N_{\vec{s}_2}} x_2^i + ... + \sum_{i=1}^{N_{\vec{s}_{4^{\mathcal{N}}}}} x_{4^{\mathcal{N}}}^i \right\rangle = \left\langle \sum_{i=1}^{N_{\vec{s}_1}} x_1^i \right\rangle + \left\langle \sum_{i=1}^{N_{\vec{s}_2}} x_2^i \right\rangle + ... + \left\langle \sum_{i=1}^{N_{\vec{s}_{4^{\mathcal{N}}}}} x_{4^{\mathcal{N}}}^i \right\rangle =$$

$$= N_{\vec{s_1}} \cdot \rho(1|t, k_{on}, k_{off}, \vec{s_1}) + N_{\vec{s_2}} \cdot \rho(1|t, k_{on}, k_{off}, \vec{s_2}) + ... + N_{\vec{s}_{4^{\mathcal{N}}}} \cdot \rho(1|t, k_{on}, k_{off}, \vec{s}_{4^{\mathcal{N}}}) \tag{22}$$

where we have exploited the independence of the set of variables to express the expected values. If we want to compute the classification performance of MEMAR we will consider, without any loss of generality, that the set of random variables $\vec{x}$ are ordered in decreasing direction according to the probability assigned by the predictor. We can then consider a threshold, which takes as values the probability of the motifs in the same decreasing order. By indicating with $n$ the discrete step of the threshold, we can compute the following quantities:

$$precision(n) = \frac{\sum_{i=1}^{n} N_{\vec{s}_i} \cdot \rho(x = 1|t, k_{on}, k_{off}, \vec{s}\ )}{\sum_{i=1}^{n} N_{\vec{s}_i}}$$

$$recall(n) = \frac{\sum_{i=1}^{n} N_{\vec{s}_i} \cdot \rho(x = 1|t, K_{on}, K_{off}, \vec{s_D})}{\sum_{i=1}^{4^{\mathcal{N}}} N_{\vec{s}_i} \cdot \rho(x = 1|t, K_{on}, K_{off}, \vec{s_D})} = \frac{\sum_{i=1}^{n} N_{\vec{s}_i} \cdot \rho(x = 1|t, K_{on}, K_{off}, \vec{s_D})}{N_{m^6A}^{\mathcal{D}}(\rho)}$$

For any given time, it is possible to compute the precision recall curve and the corresponding area under the curve as a function of time: $AUPRC(t)$. In the same way, the ratio of $m^6A$ in the dataset, can be computed, at a given time, as:

$$ratio = \frac{N_{m^6A}^{\mathcal{D}}(\rho)}{\sum_{i=1}^{4^{\mathcal{N}}} N_{\vec{s}_i}} = \frac{\sum_{i=1}^{4^{\mathcal{N}}} N_{\vec{s}_i} \cdot \rho(1|t, k_{on}, k_{off}, \vec{s}_i)}{\sum_{i=1}^{4^{\mathcal{N}}} N_{\vec{s}_i}} \tag{23}$$

If the motifs are present in the dataset in an equivalent amount, $\vec{N}^{\mathcal{D}} = (N_{\vec{s}_1}, ...N_{\vec{s}_{4^{\mathcal{N}}}}) = (N_{uni}, ..., N_{uni})$, the quantities computed simplify as shown in section *AUPRC from the two-state model* of the main article (Eqs. 13-15).

#### Binding Parameter variations between datasets

Starting from a given set of binding constants for $m^6A$ writer and erasers, $k_{on}(\vec{s}\ )$ and $k_{off}(\vec{s}\ )$ from a dataset $\mathcal{D}$, it is possible to assess potential parameter variations, $k'_{on}(\vec{s}\ )$ and $k'_{off}(\vec{s}\ )$, given a different, generic dataset $\mathcal{D}'$. Assuming that only $m^6A$ writer constants or equivalently only erasers ones vary, according to the biological or experimental differences between datasets $\mathcal{D}$ and $\mathcal{D}'$, we will exploit two different hypothesis to mathematically describe variations. The first is considering only a constant uniform rescaling $\alpha_{on}/\alpha_{off}$ of parameters, independently of the actual motif: $k'_{on}(\vec{s}\,) = \alpha_{on} \cdot k_{on}(\vec{s}\,)$ (or equivalently $k'_{off}(\vec{s}\,) = \alpha_{off} \cdot k_{off}(\vec{s}\,)$). The second hypothesis consists in the relaxation of the previous assumption, considering a non-uniform rescaling $\alpha(\vec{s}\,)$: $k'_{on}(\vec{s}\,) = \alpha_{on}(\vec{s}\,) \cdot k_{on}(\vec{s}\,)$ (or equivalently $k'_{off}(\vec{s}\,) = \alpha_{off}(\vec{s}\,) \cdot k_{off}(\vec{s}\,)$). In the first case, it is possible to numerically compute the value of the actual rescaling constant by minimizing the error between the inferred MEMAR-C probability $p'(1|\vec{s}\,)$ from the dataset $\mathcal{D}'$ and the dynamical two-state model $\rho$ with modified methylation writer binding constant (or equivalently of de-methylation one):

$$\mathcal{L}_1(\alpha) = \sum_{\vec{s} \in \mathcal{A}^{\mathcal{N}}} \mathcal{L}_1^{\vec{s}}(\alpha) = \sum_{\vec{s} \in \mathcal{A}^{\mathcal{N}}} \left| \rho(1|t, \alpha_{on} \cdot k_{on}, k_{off}, \vec{s}\,) - p'(1|\vec{s}\,) \right| \tag{24}$$

The minimization of this loss can give close to zero-values only if the uniform assumption is valid. Quantification of this validity can be assessed as explained in main article section *Kinetic parameter variations in altered-methylation experiments.* In the second case, it is possible to obtain each non-uniform rescaling parameter $\alpha_{on}(\vec{s}\,)$ by a motif-dependent minimization:

$$\mathcal{L}_1^{\vec{s}}(\alpha) = |\rho(1|t, \alpha_{on} \cdot k_{on}, k_{off}, \vec{s}\,) - p'(1|\vec{s}\,)| \qquad \forall \vec{s} \tag{25}$$

This will ensure a zero-loss value, and therefore a perfect description of inferred probabilities from the two-state dynamical model.

**From inferred dataset, to control conditions and altered methylation levels** We detail here the variations of parameters from all the datasets considered in the main article. Starting from the training set from GLORI data in HEK293T we first consider the two-state dynamical model to describe the methylation probability $p(1|\vec{s}\,)$ considering the equilibrium condition as described in paragraph *Two-state dynamical model* and Eq. S18:

1. $\rho(1|t, k_{on}, k_{off}, \vec{s}\,) = \dfrac{k_{on}(\vec{s}\,)}{k_{on}(\vec{s}\,) + k_{off}(\vec{s}\,)}$, with $k_{on}(\vec{s}\,) = p(1|\vec{s}\,)$, $k_{off}(\vec{s}\,) = 1 - p(1|\vec{s}\,)$ and $t \to \infty$ .

By construction, the two-state model before mentioned perfectly describes MEMAR-C methylation probability of motifs in the given dataset. We can then relax the equilibrium hypothesis considering a out-of-equilibrium framework. For this end, we will consider the two-state model at finite time $t = \tilde{t}$, and we will then considered a set modified datasets, with different equilibrium probabilities $\tilde{\rho}$, with higher $m^6A$ ratio: $\tilde{\rho} > \rho$. This can be obtained, for instance, decreasing by a constant factor the de-methylation rates: $\tilde{k}_{off}(\vec{s}\,) = \alpha_{off} \cdot k_{off}(\vec{s}\,)$ (with $\alpha_{off} < 1$). Indeed, if we consider $\tilde{k}_{on}(\vec{s}\,) = k_{on}(\vec{s}\,)$ for each motif and the de-methylation rates as indicated above, we can write the constraint as follows:

$$\tilde{\rho} > \rho \to \frac{\tilde{k}_{on}}{\tilde{k}_{on} + \tilde{k}_{off}} > \frac{k_{on}}{k_{on} + k_{off}} \to \frac{p}{p + \alpha_{off} \cdot (1-p)} > \frac{p}{p + 1 - p}$$

$$p + a \cdot (1-p) < 1 \to \alpha_{off} < 1$$

with the condition that $p \in (0, 1)$. We will in this case consider the generic two-state probability distribution as follows:

2. $\rho(1|t, k_{on}, k_{off}, \vec{s}\,) = \dfrac{k_{on}(\vec{s}\,)}{k_{on}(\vec{s}\,) + k_{off}(\vec{s}\,)} \cdot \left(1 - e^{-t \,\cdot\, (k_{on}(\vec{s}\,) + k_{off}(\vec{s}\,))}\right)$, with $k_{on}(\vec{s}\,) = p(1|\vec{s}\,)$, $k_{off}(\vec{s}\,) = \alpha_{off} \cdot (1 - p(1|\vec{s}\,))$ and $t = \tilde{t}$.

The set of new parameters $\{\alpha_{off}, \tilde{t}\}$ can be chosen as done in main section *A two-state dynamical model provides interpretation of performance scaling* in order to go from an equilibrium framework (1), to a non-equilibrium one (2).

For the following, we will always consider the equilibrium model, therefore as in (1), only varying either the methylation/de-methylation binding constants to describe variations to different datasets. We now take into account the deep-sequencing dataset in HEK293T cell line, in control conditions. This dataset varies from the training one for the higher reading depth of transcripts, leading to much more detected methylated sites. Since we would like to distinguishing binding parameters variations from control conditions to altered methylation conditions (obtained as well at higher sequencing depths), we infer the exact values of the binding parameters to have a zero-deviation from the inferred probabilities of the new dataset (Suppl. Fig. S13A), minimizing Eq. S25, and exploiting the relation:

3. $k_{on}^{CC_{HEK293T}}(\vec{s}\,) = \alpha_{on}^{CC_{HEK293T}}(\vec{s}\,) \cdot p(1|\vec{s}\,)$ and $k_{off}^{CC_{HEK293T}}(\vec{s}\,) = (1 - p(1|\vec{s}\,))$

Finally, a similar analysis can be obtained by considering the HeLa cell line (Suppl. Fig. S13B). The full set of parameters which perfectly describe the methylation probability of motifs, is obtained similarly to the control conditions of HEK293T at (3), by minimizing Eq. S25, following the relationships below:

4. $k_{on}^{CC_{HeLa}}(\vec{s}\,) = \alpha_{on}^{CC_{HeLa}}(\vec{s}\,) \cdot k_{on}^{CC_{HEK293T}}(\vec{s}\,)$ and $k_{off}^{CC_{HeLa}}(\vec{s}\,) = k_{off}^{CC_{HEK293T}}(\vec{s}\,)$

Having obtained the values of the binding parameters in control conditions for both cell lines, it is possible to modify either writer or erase parameters, by an uniform rescaling ($\alpha_{on}/\alpha_{off}$), obtained from (3) or (4), following Eq. S24 to describe altered methylation datasets as detailed in main paragraph *Kinetic parameter variations in altered-methylation experiments* (Suppl. Fig. S14).

An alternative approach, is to apply directly the uniform-rescaling of the two-state model, starting from (1), corresponding to the binding parameters of the standard inference HEK293T dataset, without having to infer the control conditions parameters from steps (3) or (4) (main Fig. 6B, Suppl. Fig. S12).
