## Supplementary Figures for "A maximum entropy model for *m*^6^*A* prediction: from human transcripts to influenza A viral segments"

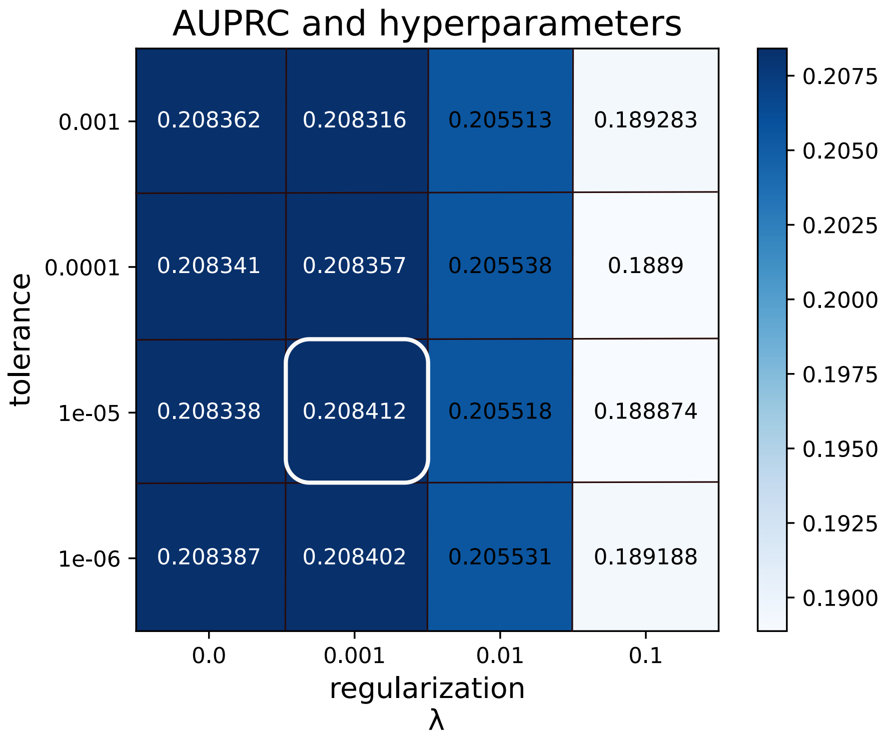


Figure 1: **MEMAR hyperparameters choice.** Heatmap showing MEMAR-C AUPRCs at distinct couples of hyperparameter values. By focusing on tolerance values only, negligible difference can be seen across rows at fixed column. Regarding the L2-regularization strengths $\lambda$, great variations, with decreasing classification performance, can be seen across columns (from second to last) at fixed row, with overall negligible difference amongst first and second columns. The set of hyperparameters which maximize model performance (shown in a rectangular shape) are chosen for MEMAR training.

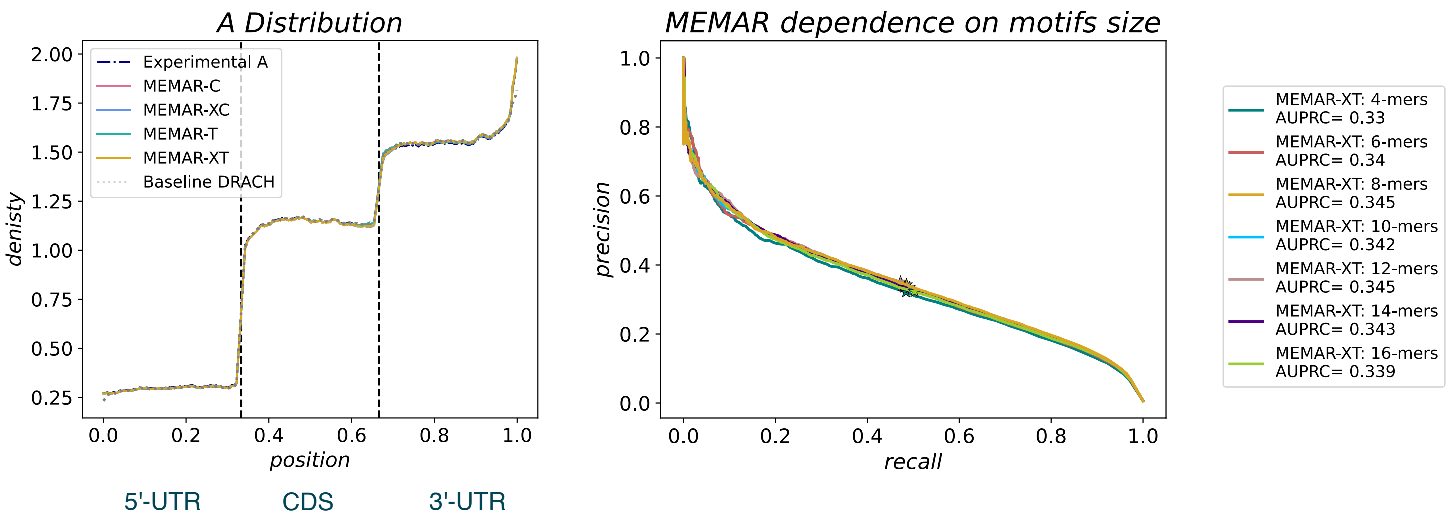


Figure 2: **Distribution of non-methylated adenosines and MEMAR dependence on motifs size. Left**) The position-distribution of non-methylated adenosines $d(x|0)$ is shown for experimental GLORI data on HEK293T cells compared to predicted ones by different models. The distribution is well captured by all MEMAR classes and also the baseline DRACH model in all 5'-UTR, CDS and 3'-UTR regions. **Right**) MEMAR-XT precision-recall curves varying training and testing scheme with larger windows size (from four to sixteen nucleotides surrounding a central adenosine). AUPRC best result of 0.345 is found by modeling 8-mer motifs, with an equivalent result for 12-mers, although being much more parametrized. Lower values are retrieved in all remaining cases.

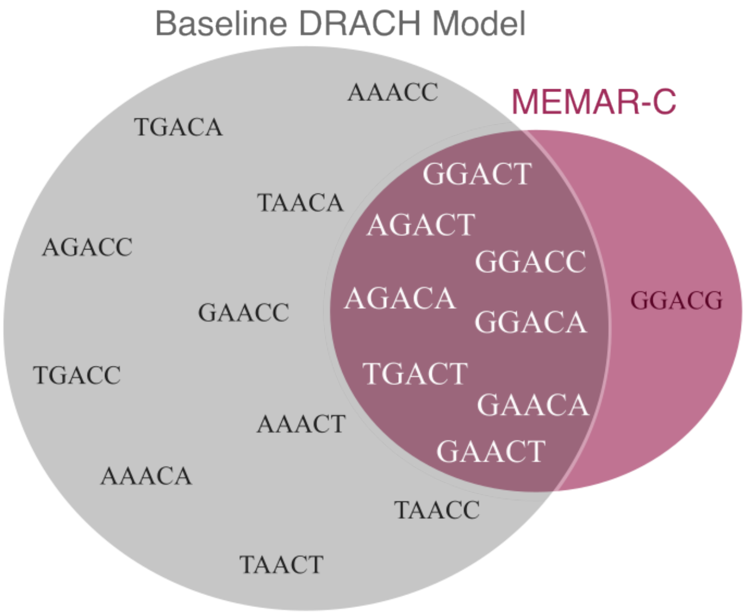


Figure 3: **Classification comparison between two binary classification methods: the baseline DRACH model and MEMAR-C at the maximum F1 threshold.** Representation of all predicted methylated motifs of the baseline DRACH model shown inside the left circle. On the right side, the subset of all possible 5-mers contained in the full list of predicted methylated 8-mer motifs surrounding a central adenosine are shown from MEMAR-C predictions.

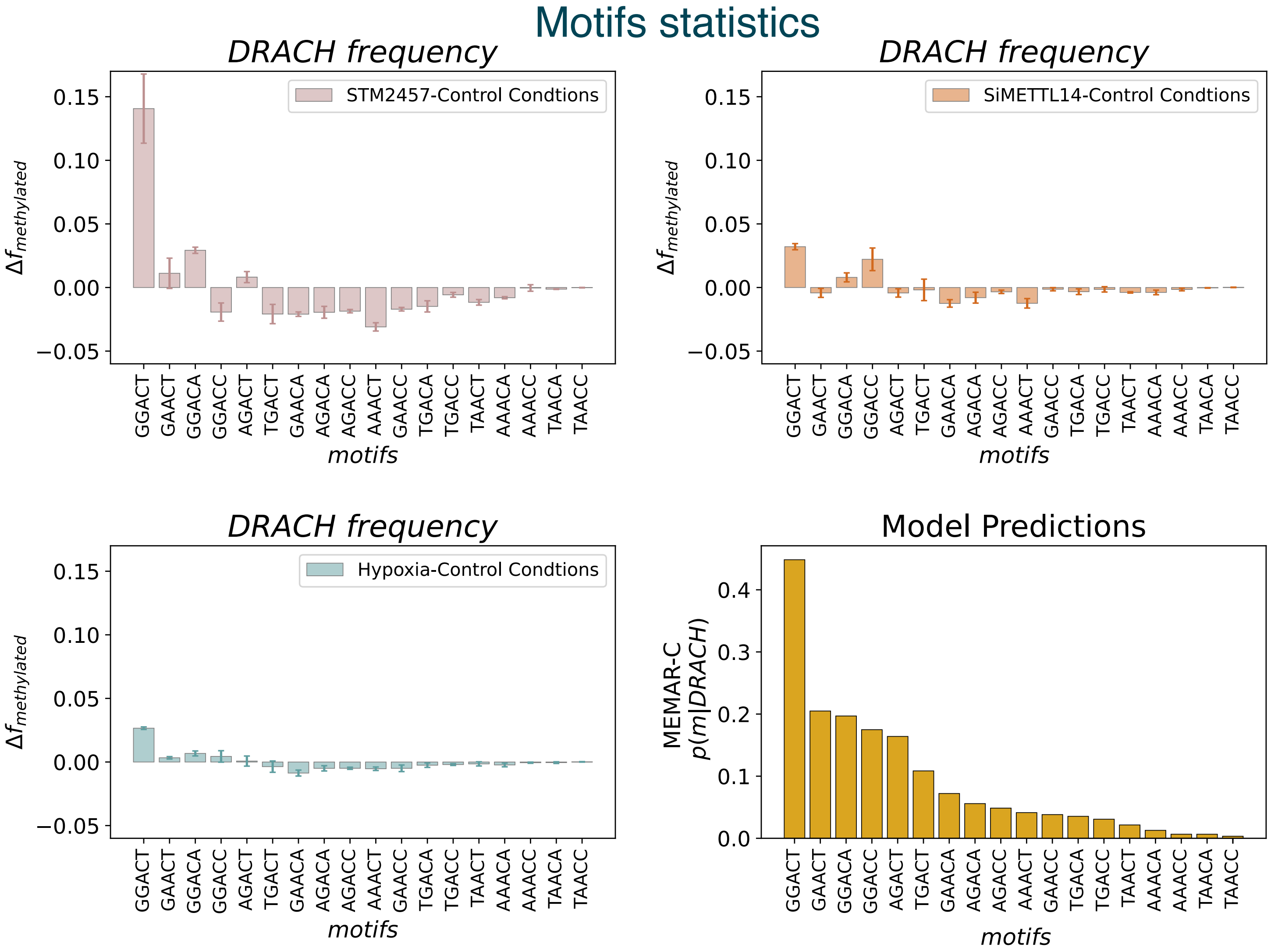


Figure 4: **Difference in the experimental DRACH methylated distribution between STM2457, siYTHDF2 and siMETTL14 with control conditions in HeLa cell line ranked by MEMAR-C predicted probabilities.** A strong variations in the motifs distribution is observed for METTL3 inhibition through STM2457, with a higher proportion of the first ranked motif *GGACT*. Regarding METTL14 knock-down and hypoxia, much lower variations are seen compared to STM2457. In particular, despite a common higher proportion of the first-ranked motif *GGACT*, slightly higher variations are observed in siMETTL14. Lastly, the predictions of MEMAR-C DRACH probabilities to be methylated are shown from highest to lowest.

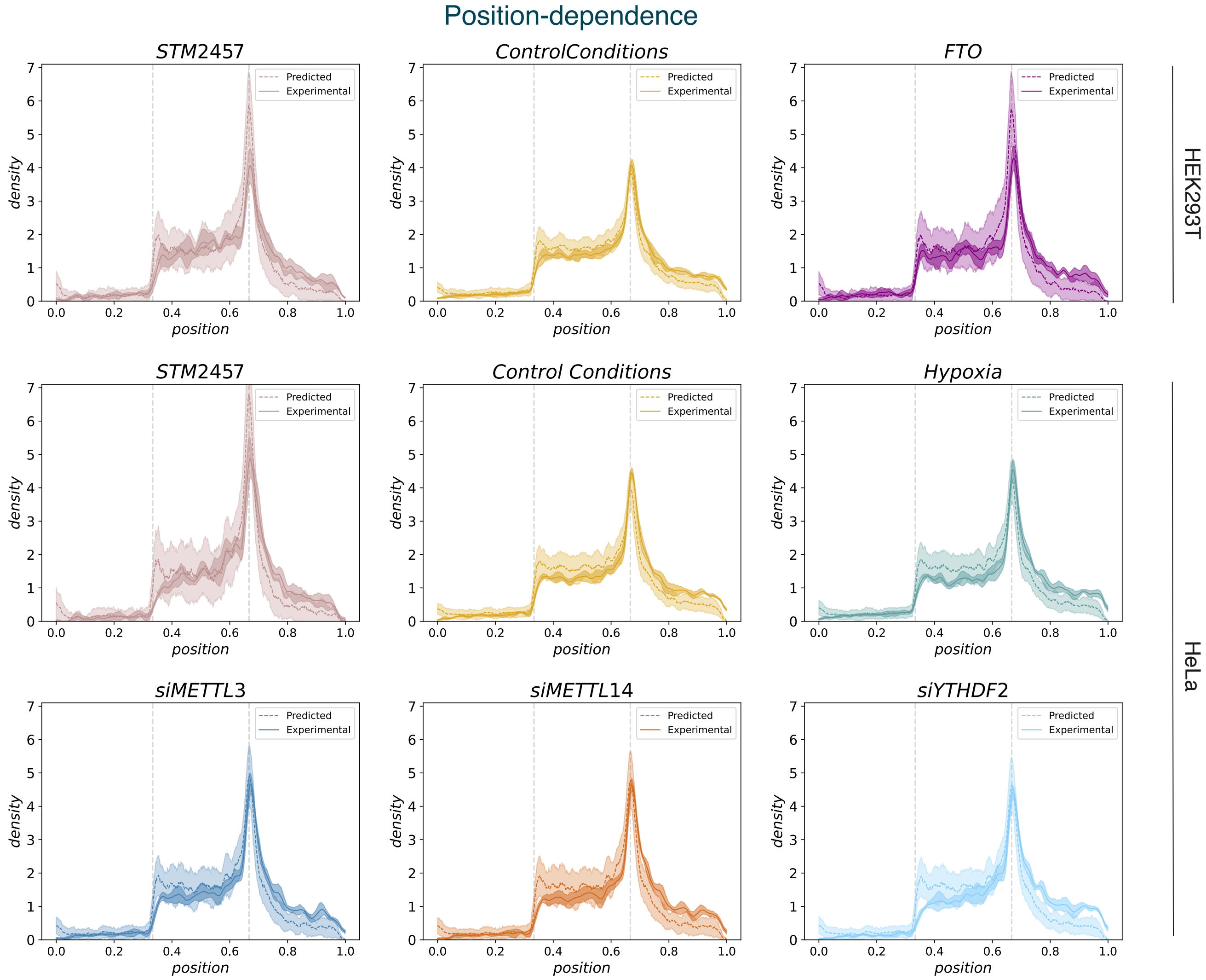


Figure 5: **Comparison of the position-dependent $m^6A$ distribution between control and altered methylation conditions against MEMAR-XT predicted methylated-sites distributions.** Overall, a similar experimental position-dependent $m^6A$ distribution $d(x|1)$ is seen across all experimental datasets and cell lines, with general agreement with MEMAR-XT predictions. An higher deviation between experimental data, and MEMAR-XT density distributions is observed particularly for FTO and STM2457 near the STOP codon. This follows the higher thresholds (at the maximum F1 score) found at Precision-Recall Curves for these datasets (see Suppl. Tabs. S8, S9). The higher thresholds leave as predicted methylated sites only strong methylation motifs which are more localized around this region.

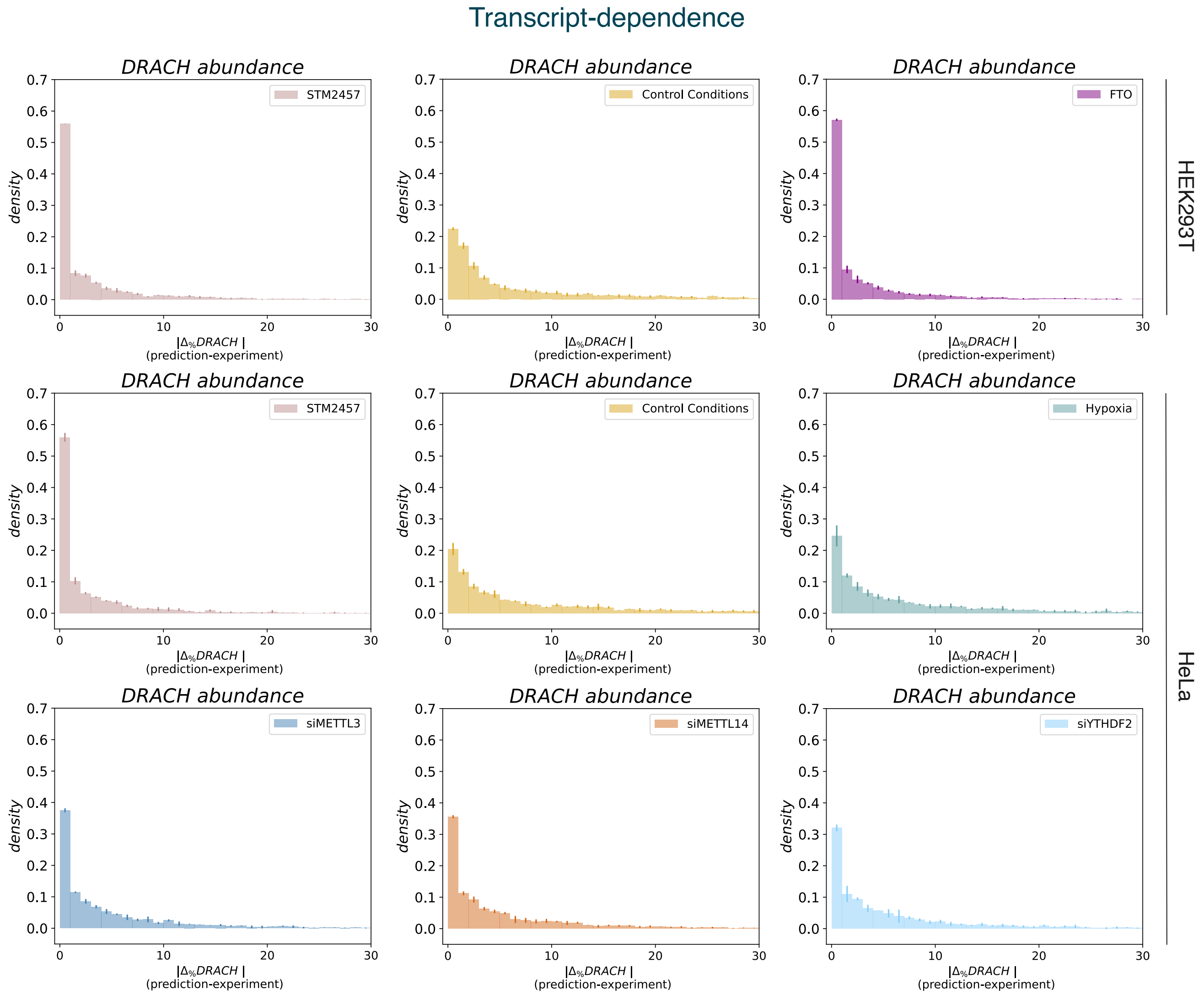


Figure 6: **Distribution of the deviation between the amount of predicted and experimental methylated DRACH sites for each transcript in control and altered methylation conditions.** Better quantification are found in strongly altered datasets as FTO and STM2457. These results follow the higher threshold for more impacted datasets on the amount of methylated sites compared to control conditions (Suppl. Tabs. S8, S9).

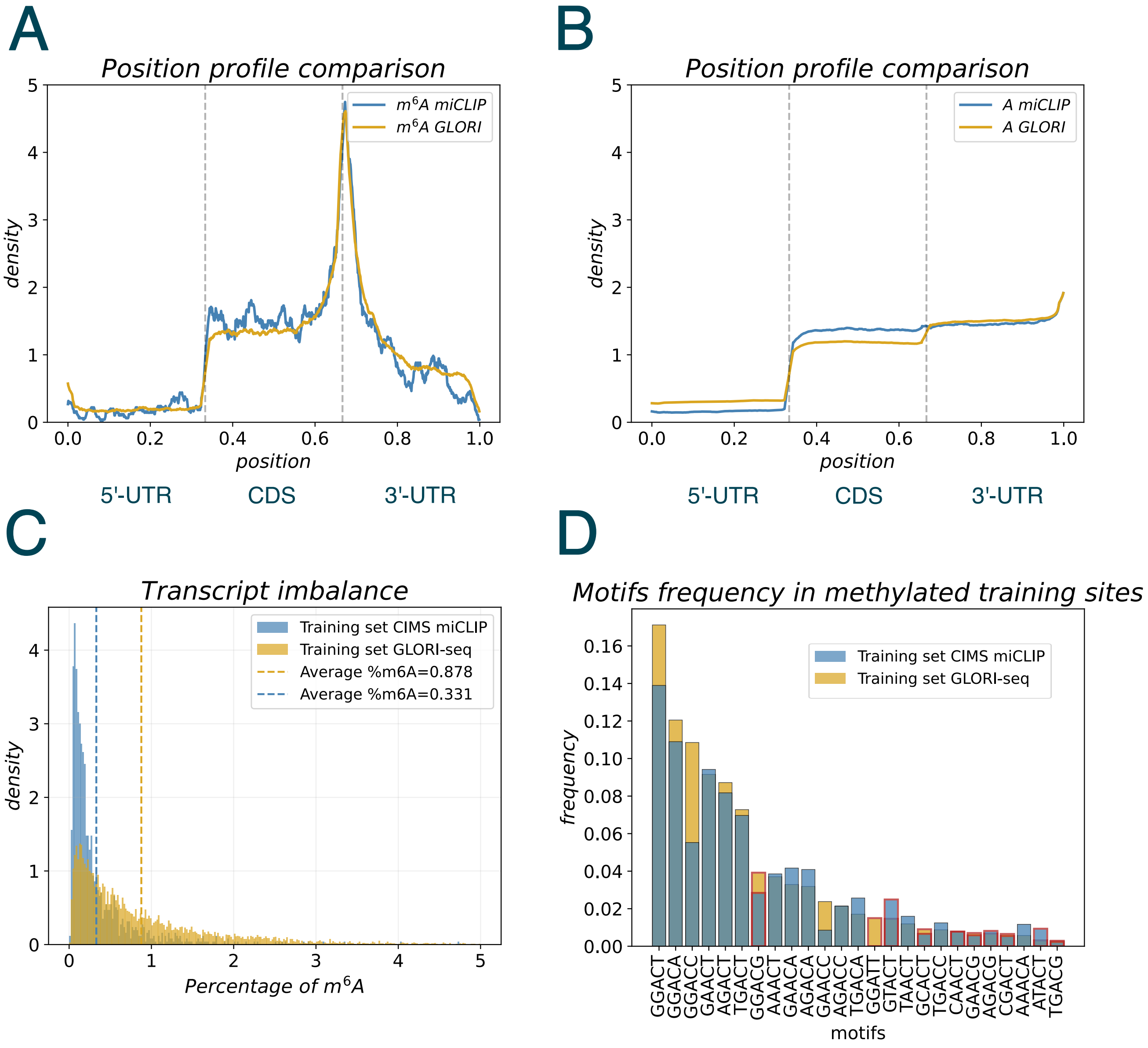


Figure 7: **Training sets details.** **A**) Position-distribution of methylated adenosines $d(x|1)$ shown for experimentally detected sites in GLORI and CIMS miCLIP training datasets, showing strong agreement. **B**) Position-distribution of methylated adenosines $d(x|0)$ shown for the same experimental datasets in **A** with overall agreement, with some deviations over 5'-UTR and CDS regions. **C**) Distribution of the percentage of $m^6A$ amongst the training transcripts from the two datasets considered. GLORI data show a longer-tailed distribution compared to miCLIP ones, indicating higher detected $m^6A$ levels in transcripts. Average values, shown with dotted lines, display a significant difference in the average percentage of methylated sites. **D**) Distribution of methylated motifs in the two training datasets, sorted by frequency values, in decreasing order, from GLORI data (focusing on the first 25 motifs only). Non-DRACH motifs have been highlighted through red rectangles.

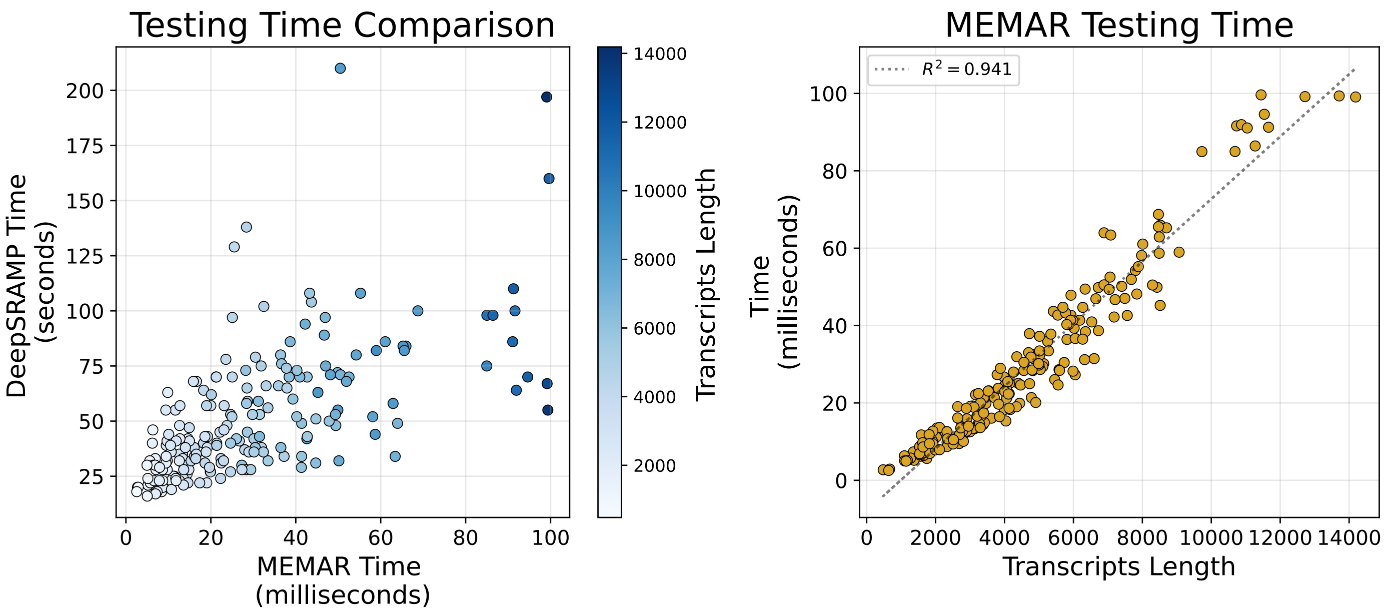


Figure 8: **Testing time analysis. Left**) Comparison between MEMAR testing time (in milliseconds) and DeepSRAMP (in seconds) with a color gradient as a function of transcripts length. MEMAR is a 1000 factor faster than DeepSRAMP. **Right**) MEMAR testing time as function of the transcripts length. A linear regression fit is shown with a dashed line. The model scales linearly, with a squared Person correlation coefficient of 0.941.

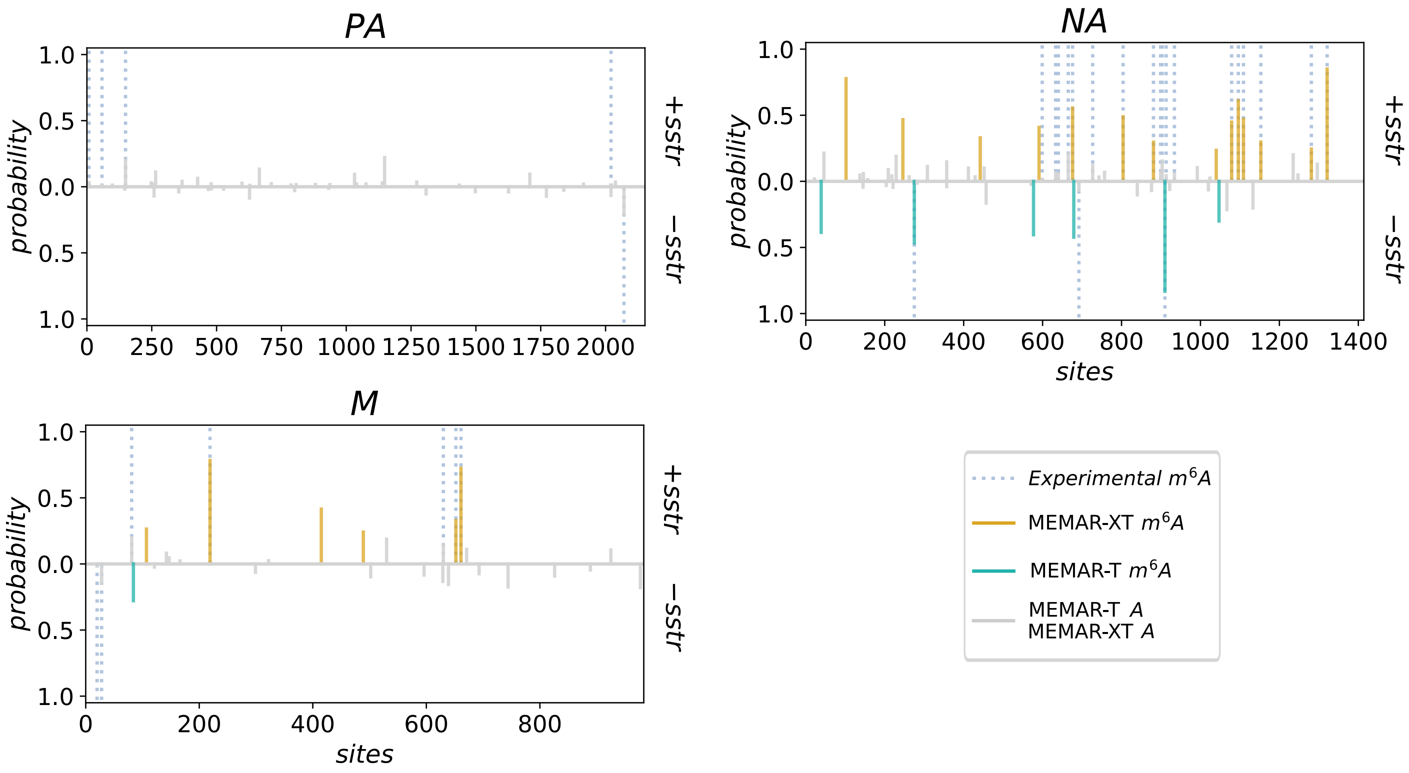


Figure 9: **MEMAR predictions over IAV segments PA, NA and M.** MEMAR-XT prediction over +sstr (upper graphs) and MEMAR-T over -sstr (lower graphs). No predicted methylated site is found in the polymerase segment PA, whilst M and NA show better classification over positive strands.

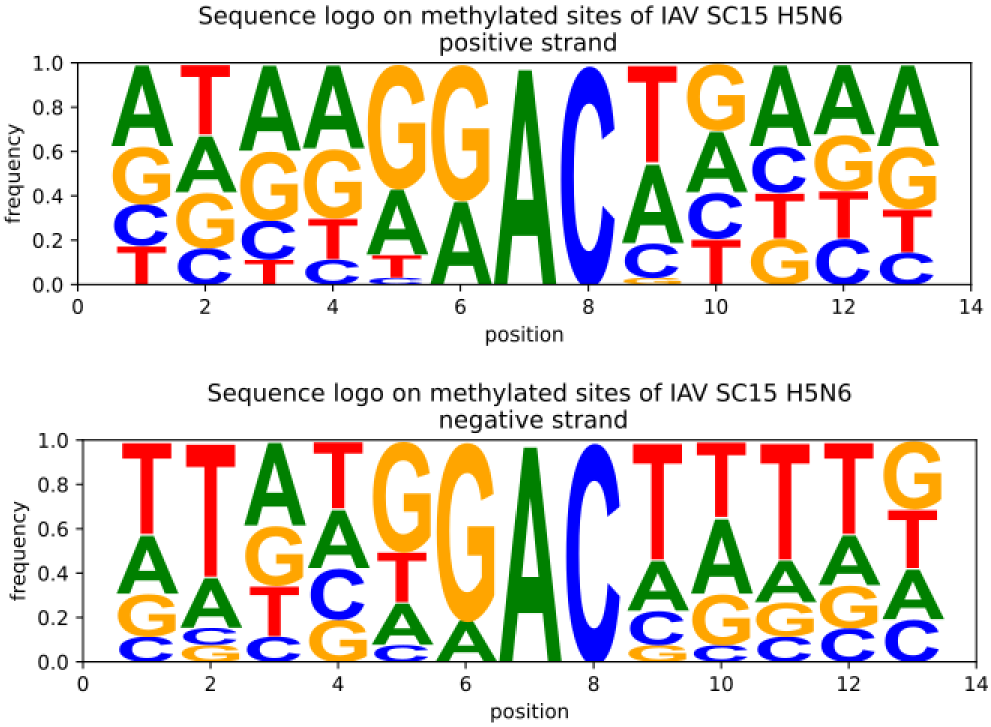


Figure 10: **Sequence logo of detected methylated sites for positive and negative strands of A/Goose/Sichuan/SC15/2015(H5N6).** Sequence logos show only RAC motifs as identified $m^6A$ sites, independently of the strand.

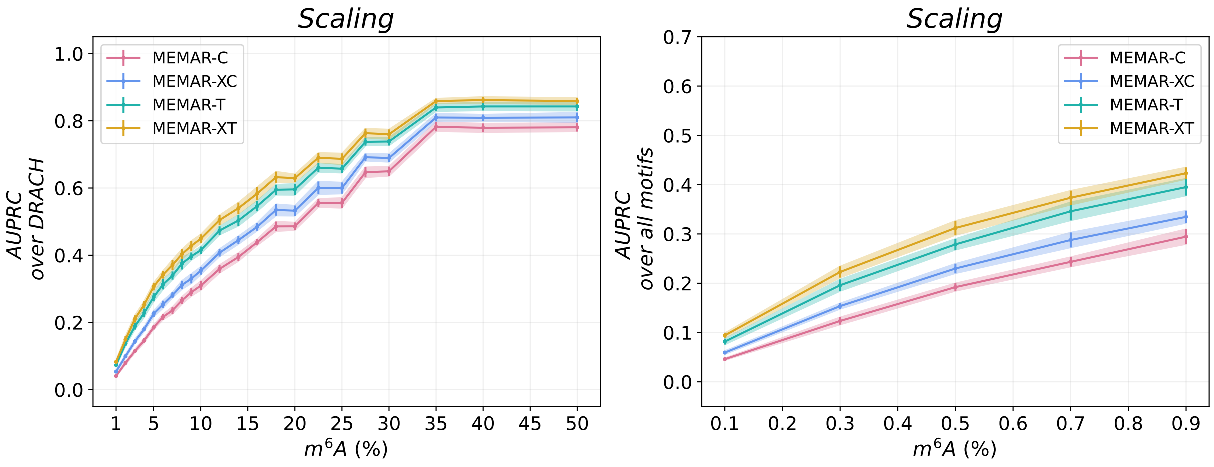


Figure 11: **Scaling of all MEMAR classes at different $m^6A$ percentages. A**) AUPRC of MEMAR classes filtering over DRACH motifs in HEK293T test set at different $m^6A$ percentages till the balance case. All classes reach at the same point, around 35%, a saturation plateau. **B**) AUPRC of MEMAR classes over all motifs at different $m^6A$ percentages. These are taken lower than panel **A** to reflect real imbalance values, which are lower due to the higher number of non-methylated motifs in transcripts without filtering over DRACH motifs.

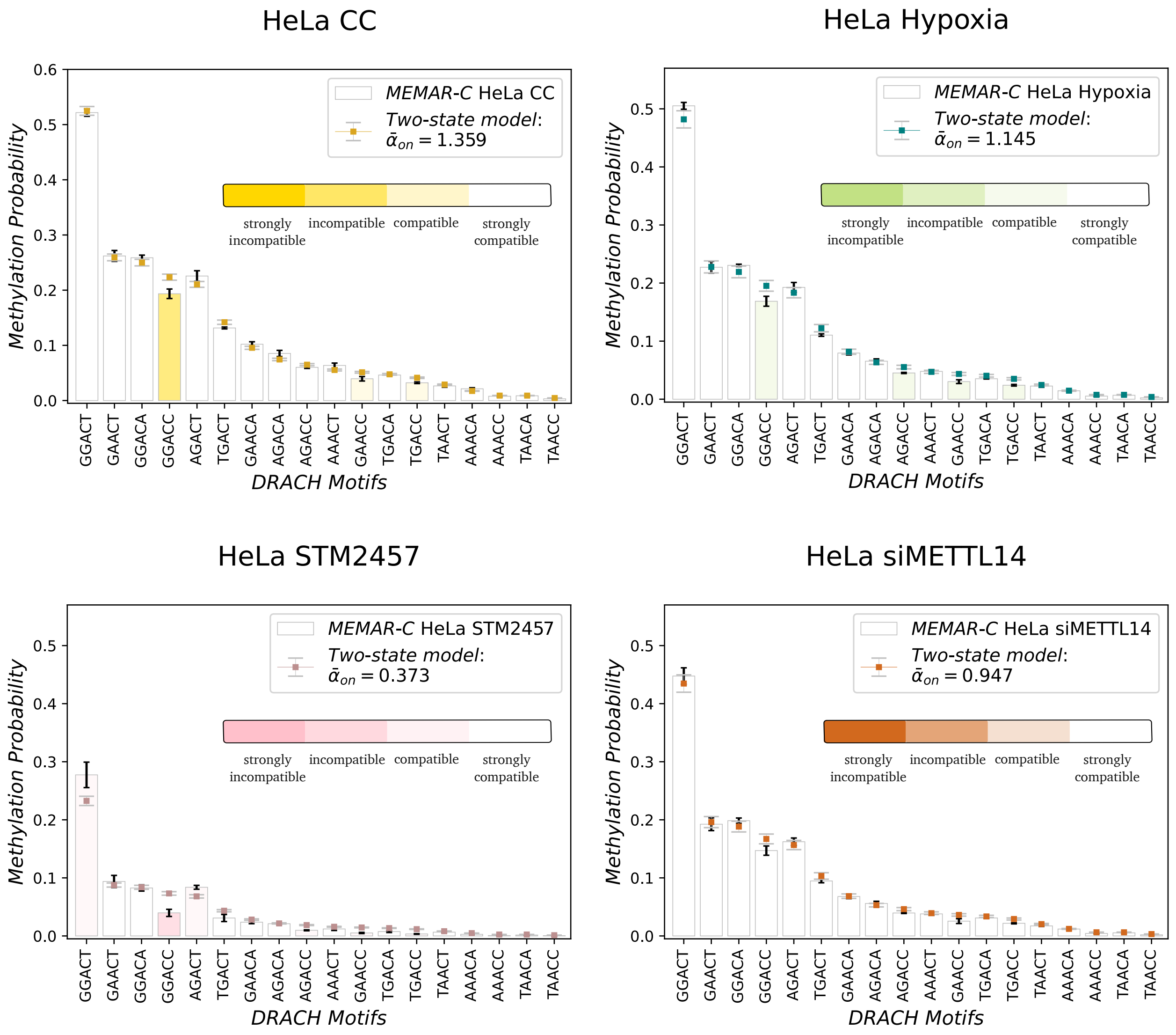


Figure 12: **Variations of methylation writer binding parameters from training dataset in HEK293T cells from GLORI data, to control or altered methylation GLORI datasets with higher sequencing depths in HeLa cells.** Re-inferred MEMAR-C probabilities in each altered dataset, consisting of hypoxia, silencing of METTL14 and inhibition of METTL3 through STM2457, or control one, are compared against the two-state model probabilities, rescaling by an uniform factor the writer binding constant from MEMAR-C trained over GLORI data in HEK293T cells. In control, the uniform rescaling parameter $\alpha_{on} > 1$ consistently indicate the higher methylation probabilities due to higher sequencing depths. Only the fourth motif presents a relevant deviation. In altered conditions, an overall agreement for the uniform rescaling is seen, although a significant deviation for the fourth ranked-motif STM2457 is present, as well as a small deviation for the first-ranked motif in the same condition, and additional small variations in hypoxia.

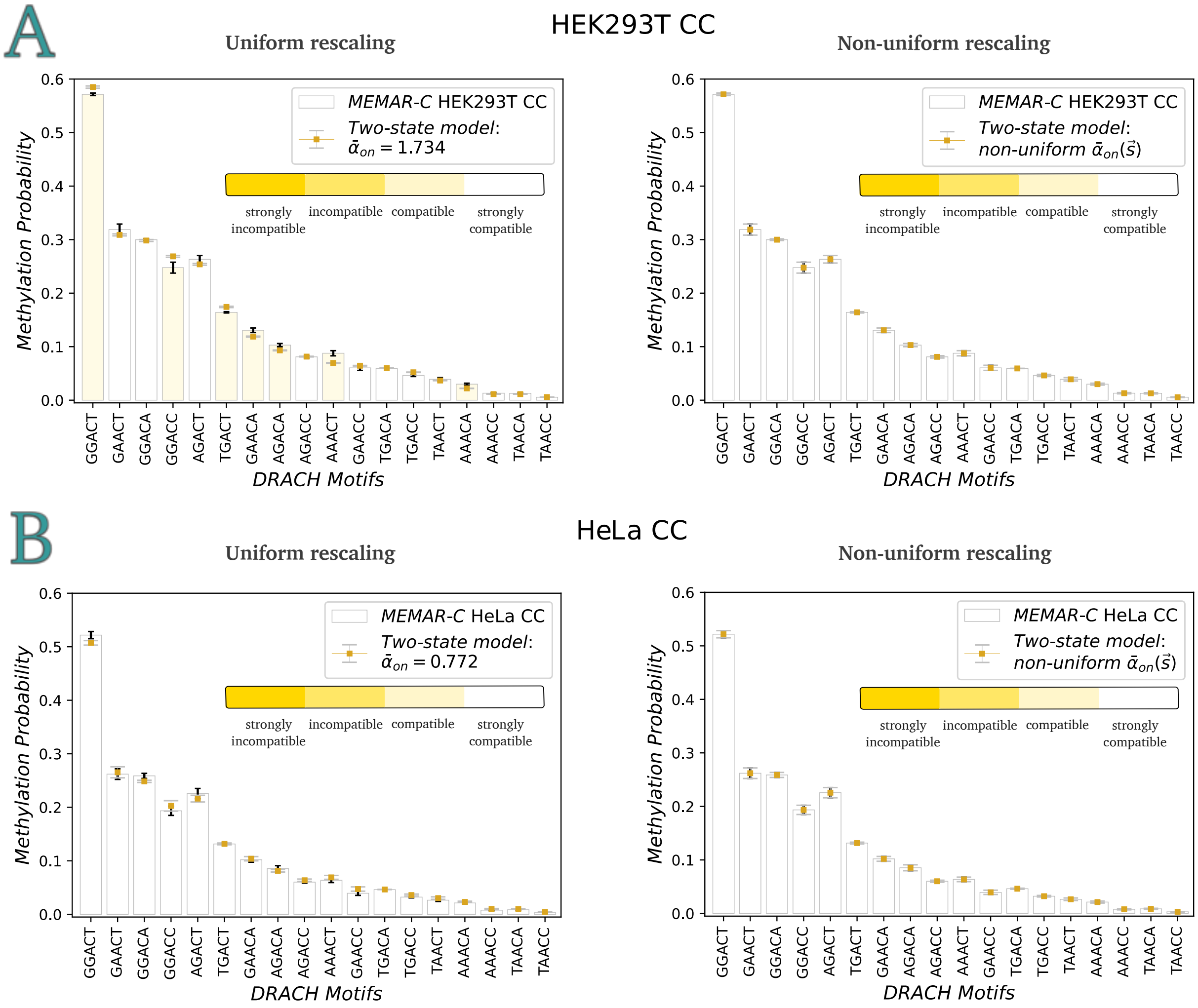


Figure 13: **Variations of methylation writer binding parameters in control conditions with deep-sequencing GLORI in HEK293T and HeLa cell lines.** Standard deviations of methylation probabilities are displayed from a three-set independent partition of each dataset. **A)** Methylation probabilities of the two state-model at equilibrium from the training dataset in GLORI HEK293T cell line to higher sequencing depth data, rescaling uniformly or non-uniformly writer binding constants. To take into account the higher-sequencing reads, a simple higher concentration of writer proteins, given by the uniform change ($\bar{\alpha}_{on} > 1$), can explain the variation in the abundance of methylated sites. Small deviations appear for a few sets of motifs, in particular for the first, the forth and the tenth motif. The non-uniform change allows to have a perfect match between the MEMAR-C methylation probabilities and the two-state ones. **B)** Same as **A**, considering a variations of writer binding constants from HEK293T in control conditions with deep-sequencing (Appendix A6). All motifs are explained by the uniform-change, with negligible differences to the non-uniform one. The two cell lines differ in the amount of deposited $m^6A$ sites ($\bar{\alpha}_{on} < 1$).

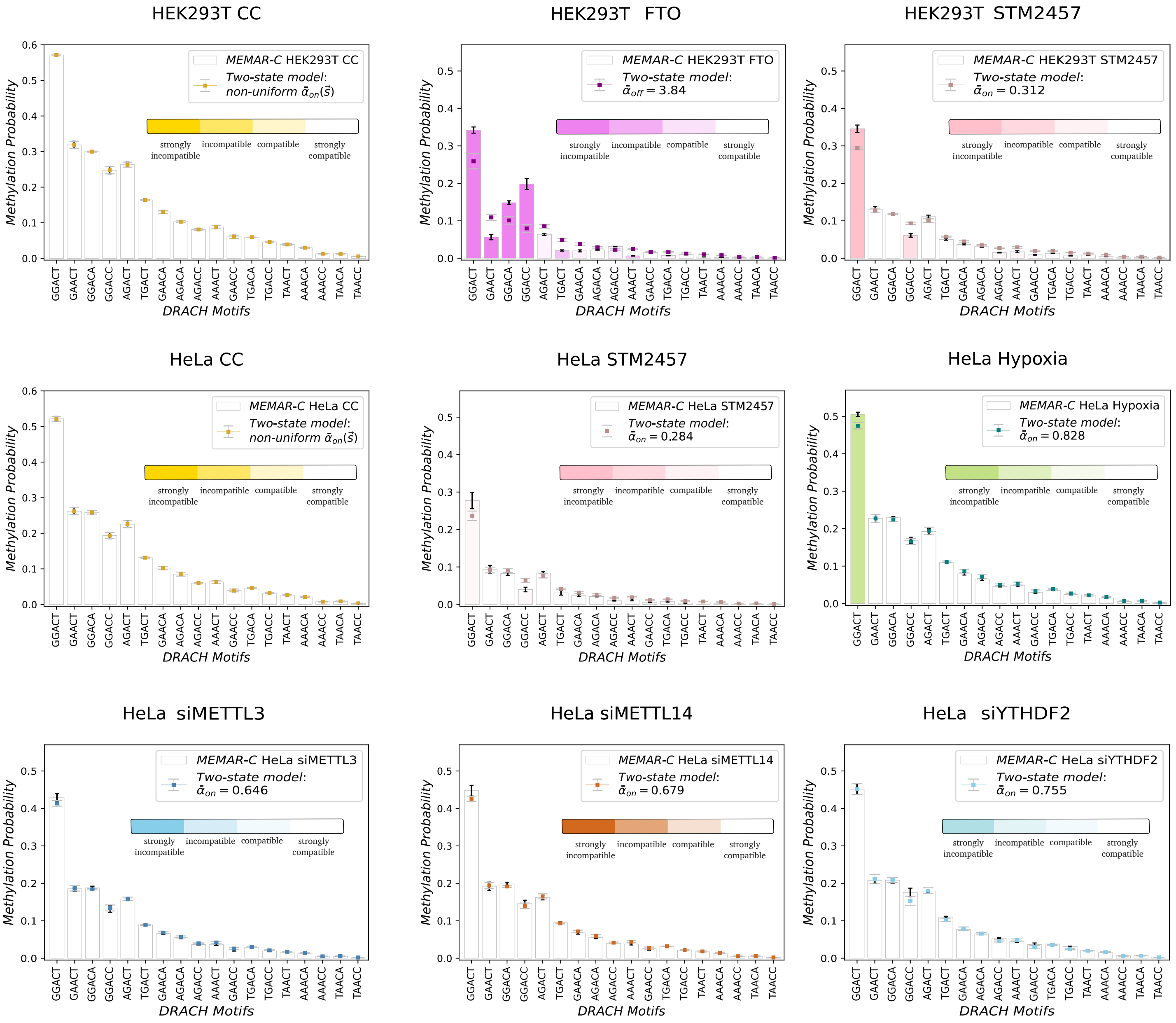


Figure 14: **Variations of the methylation writer/eraser binding parameters in altered methylation conditions from HEK293T or HeLa GLORI data in control conditions.** In the figure, the two-state model is adjusted either to match MEMAR-C predicted probabilities in HEK293T or HeLa GLORI data in control conditions, through a non-uniform rescaling of the binding parameters (Suppl. Fig. S13). Uniform rescaling of the two-state model are made for all altered conditions from control conditions in the given cell line. In particular $\bar{\alpha}_{on} < 1$ or equivalently $\bar{\alpha}_{off} > 1$ agree to a lower amount of methylated sites found in altered conditions compared to control ones. Deviations from an uniform rescaling of the two-state model are found for few motifs in HEK293T FTO and STM2457, and only to much a smaller extent to HeLa STM2457. All remaining altered conditions through silencing of $m^6A$ writers or readers, are perfectly explained by the uniform-rescaling, with only a small deviation for the first ranked motif in hypoxia.

.

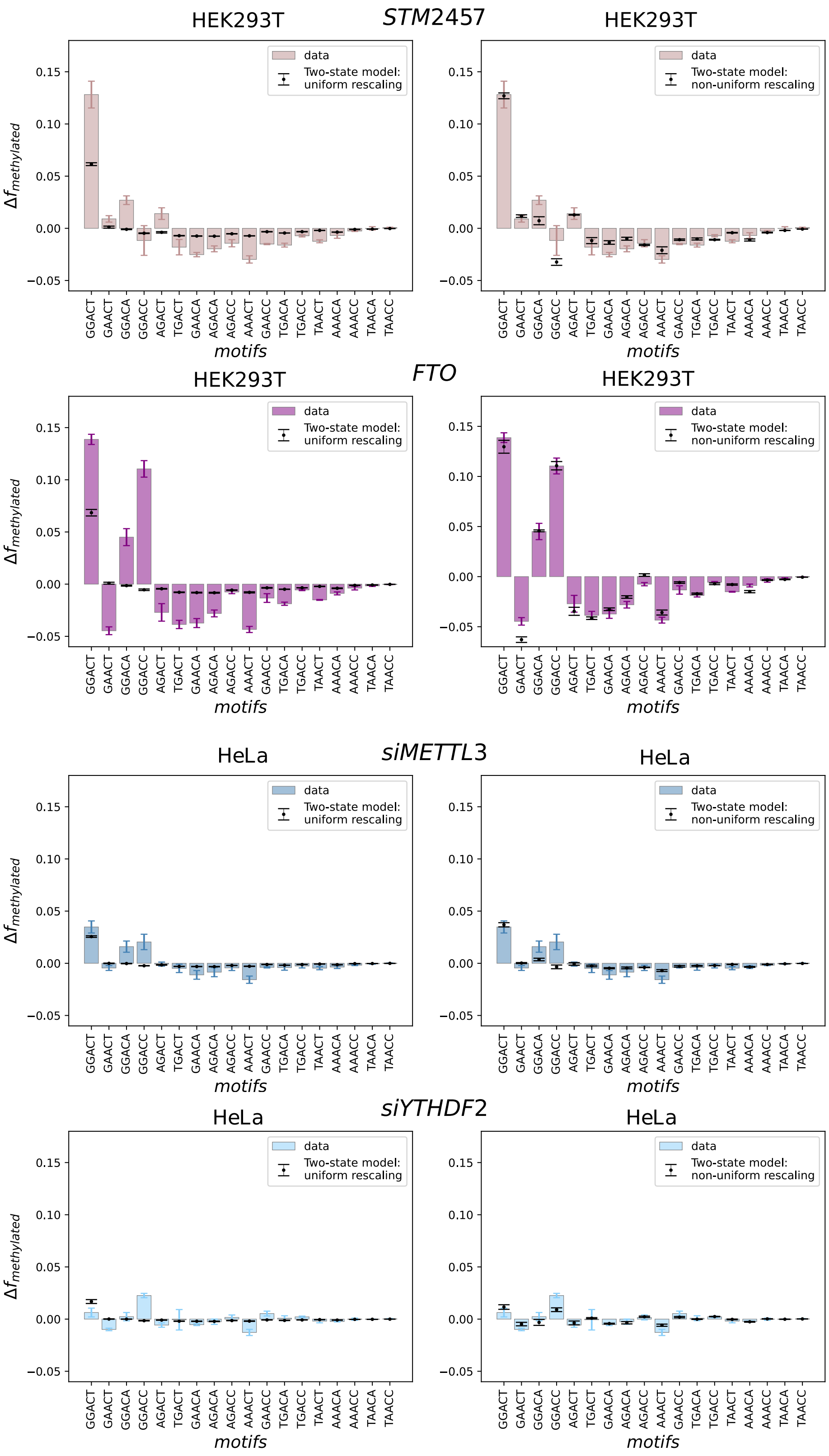


Figure 15: **Differences between DRACH methylation frequencies in altered and control conditions: observed data against two-state model predictions** Motifs are displayed following the ranking of MEMAR-C predicted probabilities (from highest to lowest). The uniform rescaling of the binding parameters in the two-state model for FTO and STM2457 in HEK293T (Suppl. Fig. S14) increase the proportion of the first-ranked motif. This, although, is not sufficient to explain the observed data (in particular large discrepancies can be seen for FTO) in agreement to the anomalies in the methylation process seen in Suppl. Fig. S14. These can be taken into account by a non-uniform rescaling of the two-state model from control conditions, improving the agreement, and greatly reproducing the observed data. In HeLa cell line, the uniform rescaling is enough to explain, to a first order, the proportion of the first ranked motif in the data, with much smaller deviations to the previous cases. The non-uniform rescaling only slightly affect the predicted profiles.
