## Supplementary Tables for "A maximum entropy model for *m*^6^*A* prediction: from human transcripts to influenza A viral segments"

| ***prior*** | GLORI | miCLIP | IAV |
|---|---|---|---|
| percentage | 0.878% | 0.331% | 1.865% |

Table 1: **Constant *prior* values expressed in percentage.**

| ***prior*** | PB2 | PB1 | PA | HA | NP | NA | M | NS |
|---|---|---|---|---|---|---|---|---|
| percentage | 0.341% | 0.181% | 0.243% | 1.970% | 5.649% | 2.791% | 2.200% | 1.543% |

Table 2: **Transcript-dependent IAV *prior* expressed in percentage.**

| $m^6A$ **imbalance** | PB2 | PB1 | PA | HA | NP | NA | M | NS |
|---|---|---|---|---|---|---|---|---|
| percentage | 1.314% | 0.633% | 0.555% | 2.671% | 4.852% | 3.950% | 1.785% | 4.633% |

Table 3: **Experimental imbalance in IAV positive strands expressed in percentage.**

| +sstr | m6A SC15 | m6A NCXJ16 | m6A full identity |
|---|---|---|---|
| PB2 | 10 | 10 | 9 |
| PB1 | 5 | 5 | 5 |
| PA | 4 | 4 | 4 |
| HA | 16 | 16 | 15 |
| NP | 23 | 23 | 20 |
| NA | 19 | 19 | 17 |
| M | 5 | 5 | 4 |
| NS | 14 | 12 | 9 |

Table 4: **Details of $m^6A$ sites from the detected experimental results on positive strands of A/Goose/Sichuan/SC15/2015(H5N6) and the ones mapped to A/Duck/Sichuan/NCXJ16/2014(H5N6).** For each segment, the total number of methylations found experimentally is shown, comparing the amount of mapped sites, accepting at most one nucleotide mutation in a window of 13 nucleotides centered over a given adenosine. On the last column, amongst the motifs found, the total number of methylations with full identity is shown.

| +sstr | position | 13-mers | NCXJ16 → SC15 |
|---|---|---|---|
| PB2 | 1984 | AAGGCGACAAAA | C → A |
| HA | 711 | GAACGGACAGCGT | C → T |
| NP | 1324 | ATGAGGACAGAAA | A → T |
| NP | 227 | GAAGGAACAGGTA | G → A |
| NP | 138 | CACAGAACTCAAA | A → G |
| NA | 804 | CGAAGAACTGAAG | G → T |
| NA | 1007 | ATGGGAAATGCGA | A→ C |
| M | 661 | ATTGGGACTCATC | G → A |
| NS | 554 | ATGATAACACAGT | C→ A |
| NS | 489 | TCCAGGACATACT | C → T |
| NS | 158 | GTCTGGACATCGA | C→ T |

Table 5: **Details of the methylated motifs mapped to the positive strands of A/Duck/Sichuan/NCXJ16/2014(H5N6) with one-nucleotide distance from original detected sites over A/Goose/Sichuan/SC15/2015(H5N6).** For each IAV segment, the position of the methylations found on NCXJ6 is shown, as well as the 13-mers motif centered over the methylated $A$, with the mutated nucleotide highlighted in red. Moreover, on the final column, the resulting mutations from NCXJ6 to SC15 are shown.

| -sstr | m6A SC15 | m6A NCXJ16 | m6A full identity |
|---|---|---|---|
| PB2 | 4 | 4 | 4 |
| PB1 | 2 | 2 | 2 |
| PA | 2 | 1 | 1 |
| HA | 3 | 3 | 3 |
| NP | 6 | 6 | 6 |
| NA | 3 | 3 | 3 |
| M | 4 | 4 | 3 |
| NS | 2 | 2 | 2 |

Table 6: **Details of $m^6A$ sites from the detected experimental results on negative strands of A/Goose/Sichuan/SC15/2015(H5N6) and the ones mapped to A/Duck/Sichuan/NCXJ16/2014(H5N6).** Information provided as in Suppl. Tab. S4.

| -sstr | position | 13-mers | NCXJ16 → SC15 |
|---|---|---|---|
| M | 136 | AAGGCGACGATAA | A → G |

Table 7: **Details of the methylated motifs mapped to the negative strands of A/Duck/Sichuan/NCXJ16/2014(H5N6) with one-nucleotide distance from original detected sites over A/Goose/Sichuan/SC15/2015(H5N6).** Information provided as in Supp. Tab. S5.

| **Experimental Condition** | Mean Threshold |
|---|---|
| Control | 0.153 |
| FTO | 0.400 |
| STM2457 | 0.426 |

Table 8: **Mean threshold values at the maximum F1 scores of the corresponding Precision-Recall curves in GLORI HEK293T cells.** Averages are computed over three sub-partitions of testing genes. Altered methylation conditions, which affect strongly methylation levels (see main Tab. 4), have much higher mean threshold values.

| **Experimental Condition** | Mean Threshold |
|---|---|
| Control | 0.183 |
| siYTDHF2 | 0.183 |
| Hypoxia | 0.210 |
| siMETTL3 | 0.300 |
| siMETTL14 | 0.300 |
| STM2457 | 0.483 |

Table 9: **Mean threshold values at the maximum F1 scores of the corresponding Precision-Recall curves in GLORI HeLa cells.** Averages are computed over three sub-partitions of testing genes. Altered methylation conditions, which affect methylation levels (see main Tab. 4), have different threshold values. Silencing RNA of YTHDF2, which affect only $m^6A$ reader, has the same threshold value as control condition, while for hypoxia an higher one is found. Silencing of $m^6A$ writers lead to increased threshold values. Methylation knock-down through increased expression of STM2457, which strongly affect $m^6A$ levels, gives the highest mean threshold over all cases considered.
